# Heterogeneity of human insular cortex: Five principles of functional organization across multiple cognitive domains

**DOI:** 10.1101/2025.03.28.646039

**Authors:** Weidong Cai, Vinod Menon

## Abstract

The insular cortex serves as a critical hub for human cognition, but how its anatomically distinct subregions coordinate diverse cognitive, emotional, and social functions remains unclear. Using the Human Connectome Project’s multi-task fMRI dataset (N=524), we investigated how insular subregions dynamically engage during seven different cognitive tasks spanning executive function, social cognition, emotion, language, and motor control. Our findings reveal five key principles of human insular organization. First, insular subregions maintain distinct functional signatures that enable reliable differentiation based on activation and connectivity patterns across cognitive domains. Second, these subregions dynamically reconfigure their network interactions in response to specific task demands while preserving their core functional architecture. Third, clear functional specialization exists along the insula’s dorsal-ventral axis: the dorsal anterior insula selectively responds to cognitive control demands through interactions with frontoparietal networks, while the ventral anterior insula preferentially processes emotional and social information via connections with limbic and default mode networks. Fourth, we observed counterintuitive connectivity patterns during demanding cognitive tasks, with the dorsal anterior insula decreasing connectivity to frontoparietal networks while increasing connectivity to default mode networks – suggesting a complex information routing mechanism rather than simple co-activation of task-relevant networks. Fifth, while a basic tripartite model captures core functional distinctions, finer-grained parcellations revealed additional cognitive domain-specific advantages that are obscured by simpler parcellation approaches. Our results illuminate how the insula’s organization supports its diverse functional roles through selective engagement of distinct neural networks, providing a new framework for understanding both normal cognitive function and clinical disorders involving insular dysfunction.

## Introduction

The insular cortex serves as a critical neural hub that integrates diverse signals necessary for human cognition, emotion, and behavior {Menon, 2024 #359}. This functionally rich brain region processes a remarkable range of information: it helps us recognize our heartbeat, feel empathy for others’ pain, maintain attention during challenging tasks, and make complex decisions under uncertainty ^1-13^. This breadth of function makes the insula unique among brain regions – it participates in both basic physiological regulation and sophisticated cognitive operations ^1,3-8,12-14^. Insular dysfunction is manifest across multiple disorders including anxiety, depression, ADHD, autism, schizophrenia, and frontotemporal dementia ^15-29^. The diversity of symptoms associated with insular dysfunction highlights a fundamental question in neuroscience: how does a single brain region coordinate such a wide array of functions?

The complexity of insular organization presents a fundamental challenge for understanding its function ^7,30-34^. The insula’s cytoarchitecture varies systematically across its extent, revealing an intricate structural gradient ^3,5,10,30,35-38^. At its anterior ventral aspect, the insula shows an agranular structure characterized by undifferentiated cortical layers II/III, distinct from the fully developed granular cortex with a canonical 6-layer architecture. Moving dorsally and posteriorly, it transitions through dysgranular regions before reaching a fully developed six-layer granular structure in posterior regions. This anatomical gradient is marked by distinct cellular features, most notably the presence of specialized von Economo neurons in anterior agranular regions ^3,10,39,40^. These distinctive neurons, characterized by their large spindle-shaped cell bodies and thick dendrites, may enable rapid information processing critical for complex cognitive functions^40,41^.

The anatomical heterogeneity of the insula has led to multiple parcellation schemes, each attempting to capture meaningful functional subdivisions. The most widely adopted approach divides the insula into three main regions: ventral anterior (vAI), dorsal anterior (dAI), and posterior (PI) insula ^7,42,43^. However, more detailed analyses have suggested finer subdivisions, with some schemes identifying up to fifteen distinct subregions ^44-47^ (**Figure 1a**). The Deen and Ryali atlases employ a tripartite parcellation based on intrinsic connectivity patterns derived from resting-state fMRI^42,48^, while the Faillenot parcellation defines six clusters based on gyral and sulcal morphology^45^. The most granular approach, the Julich atlas, identifies fifteen distinct areas based on cytoarchitectonic boundaries^44^. Each parcellation approach reflects different aspects of insular organization—intrinsic connectivity, morphological characteristics, or cytoarchitecture—raising crucial questions about which level of anatomical granularity best captures the functional heterogeneity of the insula. Whether the widely-used tripartite organization sufficiently delineates functional specialization or more fine-grained parcellations provide additional insights remains an open question with important implications for understanding this complex brain region.

**Figure 1.**
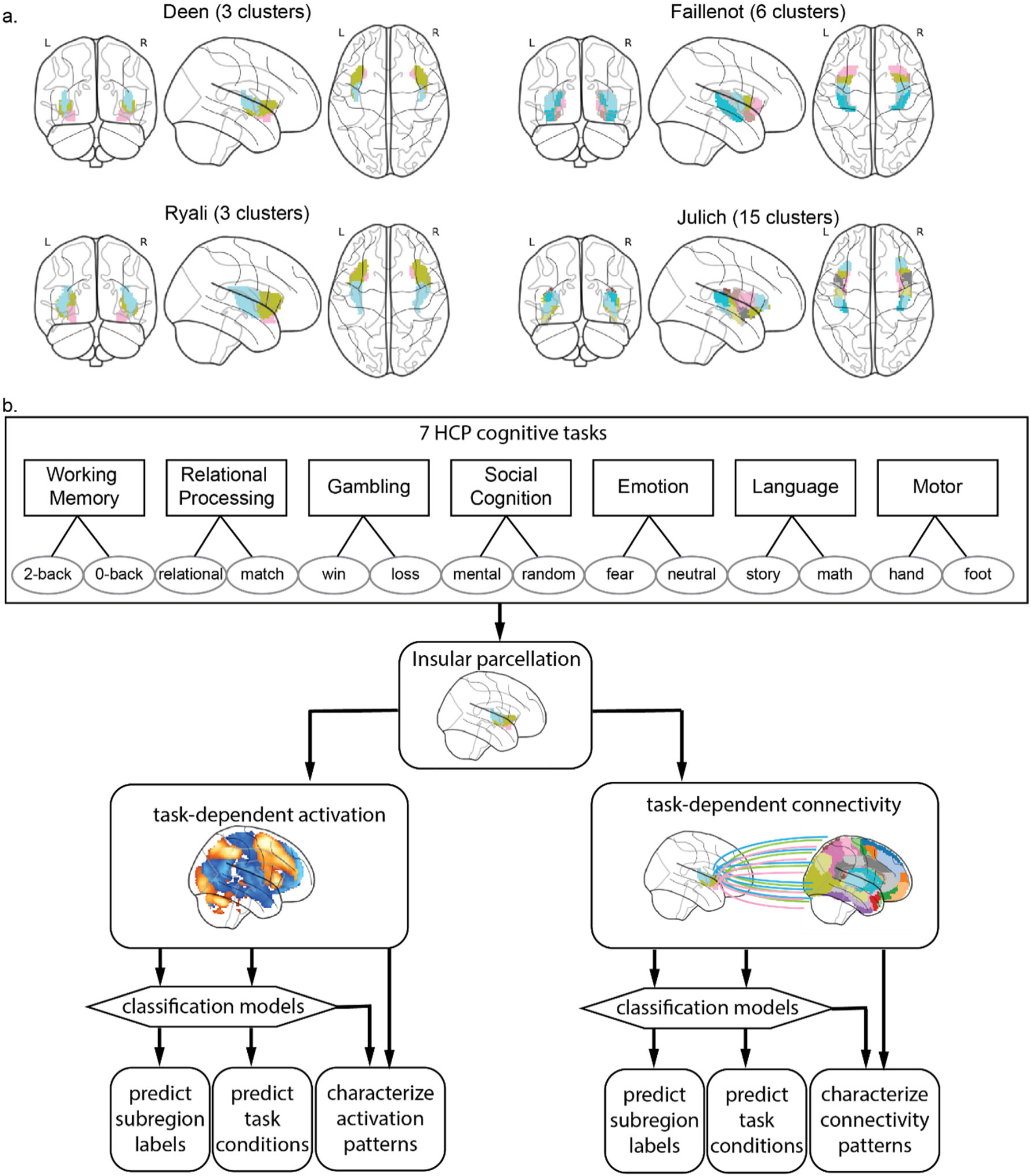
Insular parcellation schemes and analytical framework. (a) Comparison of four anatomical parcellation approaches: Deen and Ryali (3 subregions based on intrinsic connectivity), Faillenot (6 subregions based on gyral/sulcal morphology), and Julich (15 subregions based on cytoarchitecture). (b) Comprehensive data analysis pipeline using seven HCP cognitive tasks. Task-dependent activation patterns within insular subregions and connectivity patterns between insular subregions and distributed brain networks were extracted. These multivariate patterns served as features in machine learning models to differentiate insular subregions and task conditions. Cross-validation procedures evaluated classification performance, followed by detailed characterization of subregion-specific activation and connectivity profiles across cognitive domains.

Current evidence suggests broad functional distinctions across insular subregions, though the precise nature of these specializations remains debated. The anterior insula appears crucial for detecting behaviorally relevant stimuli and orchestrating cognitive control ^6,49-51^, while posterior regions primarily integrate bodily signals ^52,53^. Within the anterior insula, evidence suggests a dorsal-ventral distinction: dorsal regions may preferentially process cognitive information, while ventral regions handle emotional and motivational signals. These functional differences align with distinct intrinsic functional connectivity patterns ^31-33,54-56^. The dAI preferentially connects with cognitive control networks, particularly the dorsal anterior cingulate cortex ^32,33,42^. In contrast, the vAI connects prominently with limbic and subcortical regions involved in emotion and reward processing ^31-33,42^. The PI maintains extensive connections with sensorimotor networks, consistent with its role in processing bodily signals.

Meta-analytic studies have consistently identified the anterior insula as one of the most frequently activated regions during cognitively demanding tasks ^6,49,57^. Its network interactions are dynamically modulated by cognitive control and attention demands ^58-62^, and disruptions to these interactions correlate with impaired cognitive functions ^63-65^. Beyond cognitive control, the anterior insula, especially the vAI, plays a crucial role in processing reward and emotional information ^66,67^, supported by strong structural connections with key affect- and reward-processing regions such as the amygdala and nucleus accumbens ^31,68,69^. However, most studies examining these functions have not systematically distinguished between vAI and dAI contributions ^38,66,67,70^, leaving important questions about functional specialization unanswered. Moreover, few studies have systematically explored functional specificity of anterior insula across a wide range of cognitive task contexts and load manipulation in the same individuals or evaluated consistency across different insular atlases.

Crucially, while previous studies have characterized insular network interactions primarily through resting-state connectivity and meta-analyses across different populations ^31,32,42,54-56,71^, we lack a systematic understanding of how insular subregions dynamically reconfigure their network interactions during active task performance. By examining task-related functional connectivity and network dynamics at the single-subject level across multiple cognitive domains, we sought to uncover fundamental principles of how the insula achieves flexible network integration while maintaining stable functional specialization. This approach allows us to distinguish between static connectivity patterns and dynamic network reorganization that supports diverse cognitive operations.

We address these fundamental questions about insular organization using the Human Connectome Project’s multi-task dataset, which offers several unique advantages for mapping functional architecture. The dataset includes seven diverse cognitive tasks that probe different aspects of insular function, from basic motor control to working memory to complex social cognition. The fMRI battery consists of Working Memory, Relational Processing, Gambling, Social Cognition, Emotion Processing, Language Processing and Motor tasks, with two key load or context manipulations (**Figure 1b**). Crucially, the large sample size (N∼1200) provides robust statistical power for detecting subtle functional distinctions. Each task systematically varies cognitive demand while maintaining consistent task structure and sensorimotor requirements, enabling the isolation of cognitive load from task-specific processes.

Our investigation employs multiple complementary approaches to characterize insular function. We examine both task-dependent insular activation patterns and insula-to-network interactions across seven cognitive domains while systematically comparing four anatomical parcellation schemes ranging from basic tripartite models to complex subdivisions with up to fifteen regions ^42-45^.

Using machine learning approaches, we address several fundamental questions: (1) Do insular subregions show reliable functional boundaries across cognitive domains? (2) How does parcellation granularity affect our ability to capture distinct cognitive and affective processes? (3) How does the insular cortex participate in multiple functions while maintaining core functional specialization?

We hypothesized that insular subregions would show distinct but complementary functional profiles: dorsal anterior regions coordinating cognitive control through frontoparietal network interactions, and ventral anterior regions integrating emotional and social information through limbic network connections. We further predicted that while basic tripartite parcellation may capture overall task differentiation, fine-grained parcellations will reveal additional specialization. Finally, we expected to find evidence for flexible, rather than fixed, task-dependent interaction between insular subregions and brain networks to support various cognitive and affective processes, reflecting a computational architecture optimized for dynamic integration of information.

## Results

### Multi-task condition-specific activation patterns differentiate insular subregions

To determine whether functional specialization exists within insular subregions, we first examined whether activation patterns across the seven diverse HCP tasks (Working Memory, Relational Processing, Gambling, Social Cognition, Emotion Processing, Language Processing and Motor) could differentiate distinct insular areas. We extracted task-dependent ROI-wise activation values (e.g., 0-back and 2-back in working memory, and loss and win in gambling) across all seven HCP cognitive tasks and used these activation profiles to distinguish between insular subregions. A linear support vector machine (linearSVM) model was trained to differentiate insular subregions, with performance evaluated using 5-fold cross validation. Statistical significance was determined through permutation testing (500 iterations). This multivariate approach allowed us to quantify the distinctiveness of functional activation patterns across insular subdivisions.

We found that task-dependent activation patterns accurately differentiated insular subregions across all parcellation schemes (all *ps*<0.002, **Figure 2a**). Classification accuracies substantially exceeded chance levels for each atlas (33.3% for 3 clusters, 16.7% for 6 clusters, and 6.7% for 15 clusters), demonstrating robust functional differentiation between insular subregions regardless of parcellation granularity. These results provide strong evidence that insular subregions maintain distinct functional profiles across diverse cognitive domains.

**Figure 2.**
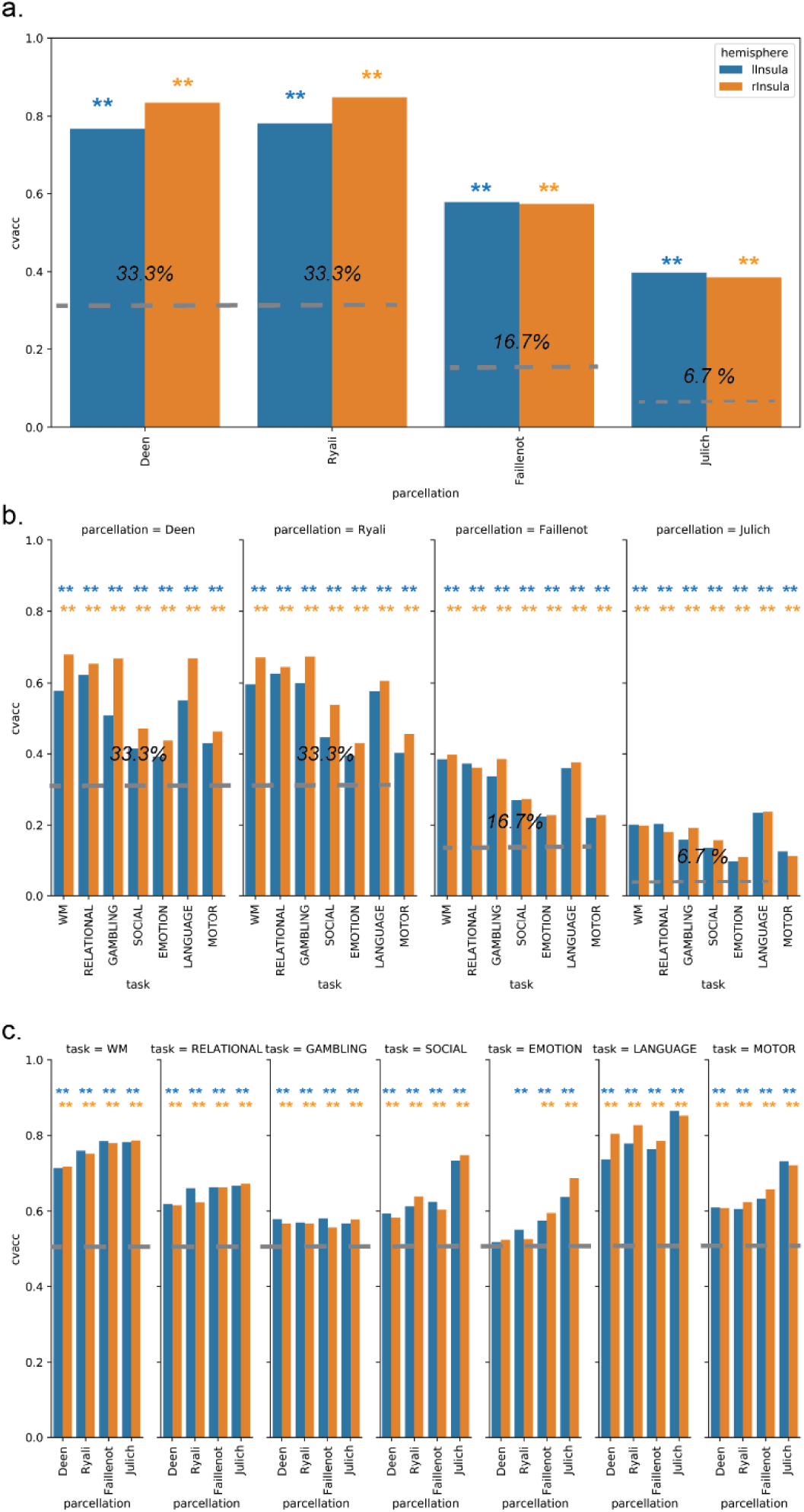
Insular subregions show distinct functional signatures across cognitive domains. (a) Classification of insular subregions using activation patterns across all seven cognitive tasks, demonstrating successful differentiation across all parcellation schemes (dashed lines indicate chance levels: Deen/Ryali: 33%, Faillenot: 16.7%, Julich: 6.7%). (b) Within-task classification of insular subregions, showing reliable differentiation even within single cognitive domains. (c) Classification of task conditions (e.g., 2-back vs. 0-back) using activation patterns across insular subregions. Blue and orange asterisks indicate significant classification (p<0.05, FWE-corrected) using left and right insular patterns, respectively, assessed via permutation testing.

### Within-task condition-specific activation patterns differentiate insular subregions

To assess whether functional differentiation was evident even within individual task contexts, we next examined whether activation patterns from single tasks could distinguish insular subregions. Using the same linearSVM approach, we found that task activation patterns from each HCP task reliably differentiated insular subregions across all parcellation schemes (all *ps*<0.002, **Figure 2b**). Notably, classification performance was highest for cognitively demanding tasks (working memory, relational processing, and language) compared to social-emotional and motor tasks. This pattern was consistent across all four parcellation atlases, suggesting that cognitive control operations may elicit more distinctive activation patterns across insular subregions than affective or sensorimotor processes. This finding aligns with the hypothesis that the insula plays a particularly important role in cognitive control through its selective engagement of specific subregions during demanding cognitive operations.

### Insular activation patterns encode task-specific cognitive states

After establishing that insular subregions maintain distinct functional profiles, we next investigated whether these regional activation patterns contained sufficient information to discriminate between different cognitive conditions within each task. Using the same linearSVM framework, we trained classifiers to differentiate between main task conditions (e.g., 0-back versus 2-back in working memory) based solely on activation patterns across insular subregions. This approach assessed the functional significance of insular activation patterns in representing distinct cognitive states.

Results revealed that activation patterns across insular subregions successfully discriminated between task conditions in most contexts (all *ps*<0.002, **Figure 2c**), with three notable exceptions: insular activation patterns from Deen’s and Ryali’s parcellations failed to significantly differentiate conditions in the emotional processing task. This selective failure suggests that emotional processing may engage insular subregions in a more distributed or overlapping manner than other cognitive domains.

Classification performance varied systematically across cognitive domains, with highest accuracies observed for working memory and language tasks (70-80%) compared to other domains (∼60%). This suggests that insular activation patterns most strongly encode cognitive states related to executive function and language processing. Interestingly, the more fine-grained Julich atlas (15 subregions) yielded superior classification performance for social, emotional, language, and motor tasks, while showing comparable performance to coarser parcellations for working memory, relational processing, and gambling tasks. This pattern indicates that finer anatomical distinctions may capture subtle functional specialization particularly relevant to social-emotional processing, while coarser distinctions adequately capture cognitive control-related states.

### Distinct activation profiles across insular subregions

To characterize functional specialization within the insula, we examined mean activation levels across insular subregions during seven HCP tasks. Repeated measures ANOVA with factors of SUBREGION (vAI, dAI, PI) and CONDITION (2 main conditions within each task) revealed significant main effects of SUBREGION across all tasks (p<0.01, FDR corrected, **Figure 3**, **Supplementary Table S6**). The dAI showed significantly higher activation levels than both vAI and PI during Working Memory and Relational Processing tasks (all ps<0.001, FDR corrected, **Figure 3**). This functional profile was consistent across parcellation schemes - regions corresponding to the dAI in Deen and Ryali atlases showed similar activation patterns to the ASG in Faillenot and Id7/Id6 in Julich atlases (**Figure 3**, **Supplementary Tabled S6-9)**.

**Figure 3.**
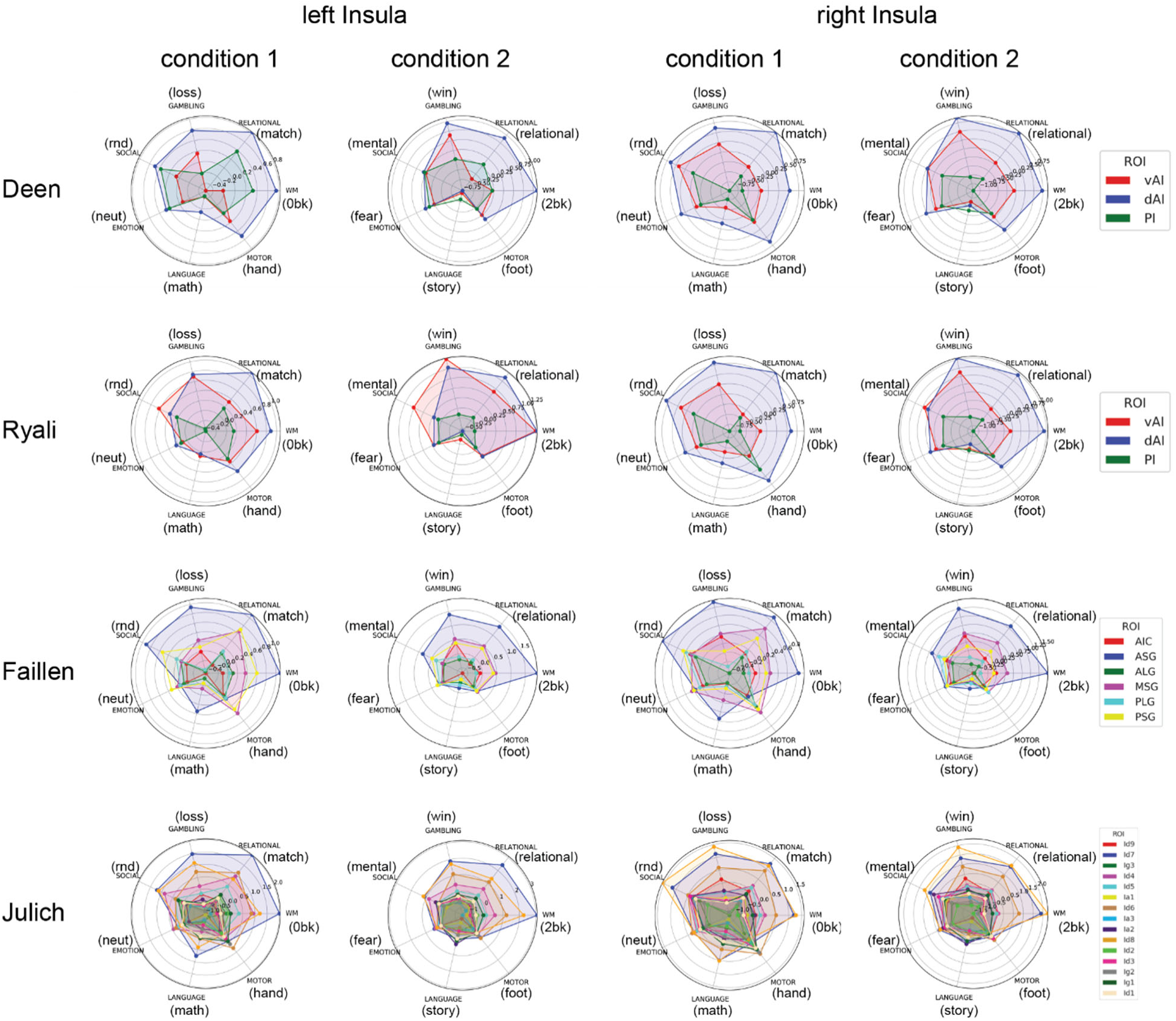
Distinct task-activation profiles reveal functional specialization among insular subregions. Radar plots illustrate condition-specific activation patterns across insular subregions for each parcellation scheme. Conditions include 2-back/0-back (Working Memory), match/relational (Relational Processing), loss/win (Gambling), random/mental (Social Cognition), fear/neutral (Emotion), story/math (Language), and hand/foot (Motor). Results reveal consistent functional organization across parcellation schemes despite different anatomical definitions.

Significant SUBREGION and CONDITION interactions (p<0.01, FDR corrected, **Figure 4**, **Supplementary Table S6**) revealed selective responses of insular subregions to specific cognitive demands. Specifically, the dAI showed heightened sensitivity to cognitive load, with significantly greater increased activation during demanding conditions in the working memory (2-back vs. 0-back) task compared to the vAI and PI (*p<*0.01, FDR corrected, **Figure 4**). Interestingly, the dAI showed significantly reduced activation during social processing (mental vs. random) and during language processing (story vs. math) compared to vAI (*p<0.01*, FDR corrected, **Figure 4**). In contrast, the vAI exhibited significantly greater activation increases during social processing (mental vs. random) compared to dAI and PI (*p<0.01*, FDR corrected, **Figure 4**), while the PI showed enhanced activation during language processing (story vs. math) and reduced activation in demanding working memory conditions (2-back vs. 0-back) than dAI and vAI (*p<0.01*, FDR corrected, **Figure 4**). These distinctive response profiles remained consistent across all four parcellation schemes (**Figure 4**, **Supplementary Tabled S6-9)**.

**Figure 4.**
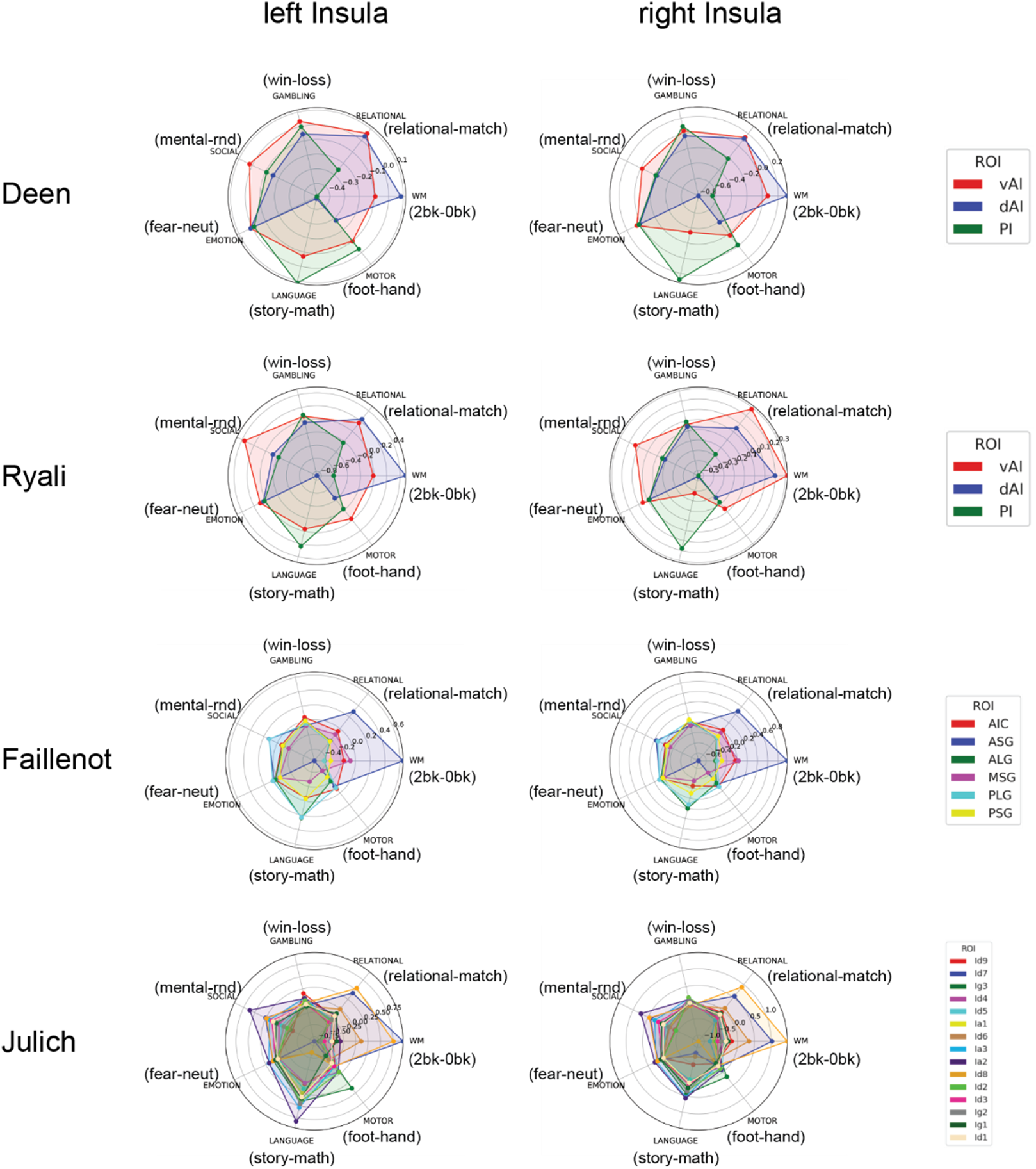
Task demands differentially modulate activation across insular subregions. Radar plots display contrast values (condition 2 vs. condition 1) across tasks for each insular subregion. Contrasts include 2-back>0-back (Working Memory), relational>match (Relational), win>loss (Gambling), mental>random (Social), fear>neutral (Emotion), story>math (Language), and foot>hand (Motor). Results reveal a dissociation between dorsal and ventral anterior insula responses to cognitive versus emotional demands.

These findings demonstrate systematic functional specialization within the insular cortex, with dAI selectively responsive to cognitive control demands, vAI preferentially engaged by social-emotional processing, and PI showing more variable responses.

### Multivariate patterns within individual insular subregions encode task-specific information

To further characterize functional specialization, we examined whether voxel-wise activation patterns within each insular subregion contained information sufficient to differentiate task conditions. Using a linearSVM approach, we evaluated classification performance for each subregion across all insular atlases.

Classification results revealed striking functional dissociations (**Figure 5, Supplementary Tables S2-5**). Within the dAI, voxel patterns achieved highest classification accuracy for discriminating between conditions in working memory (0-back vs. 2-back), relational processing (match vs. relational), and language tasks (story vs. math). In contrast, the vAI showed superior classification performance for distinguishing between conditions in gambling (loss vs. win) and emotion processing (neutral vs. fear) tasks. The PI exhibited highest classification accuracy for motor task conditions (foot vs. hand) among all subregions. This triple dissociation was consistent across all parcellation schemes, providing converging evidence for distinct computational roles within the insular cortex. Similar results were found in the Ryali, Faillenot and Julich atlases.

**Figure 5.**
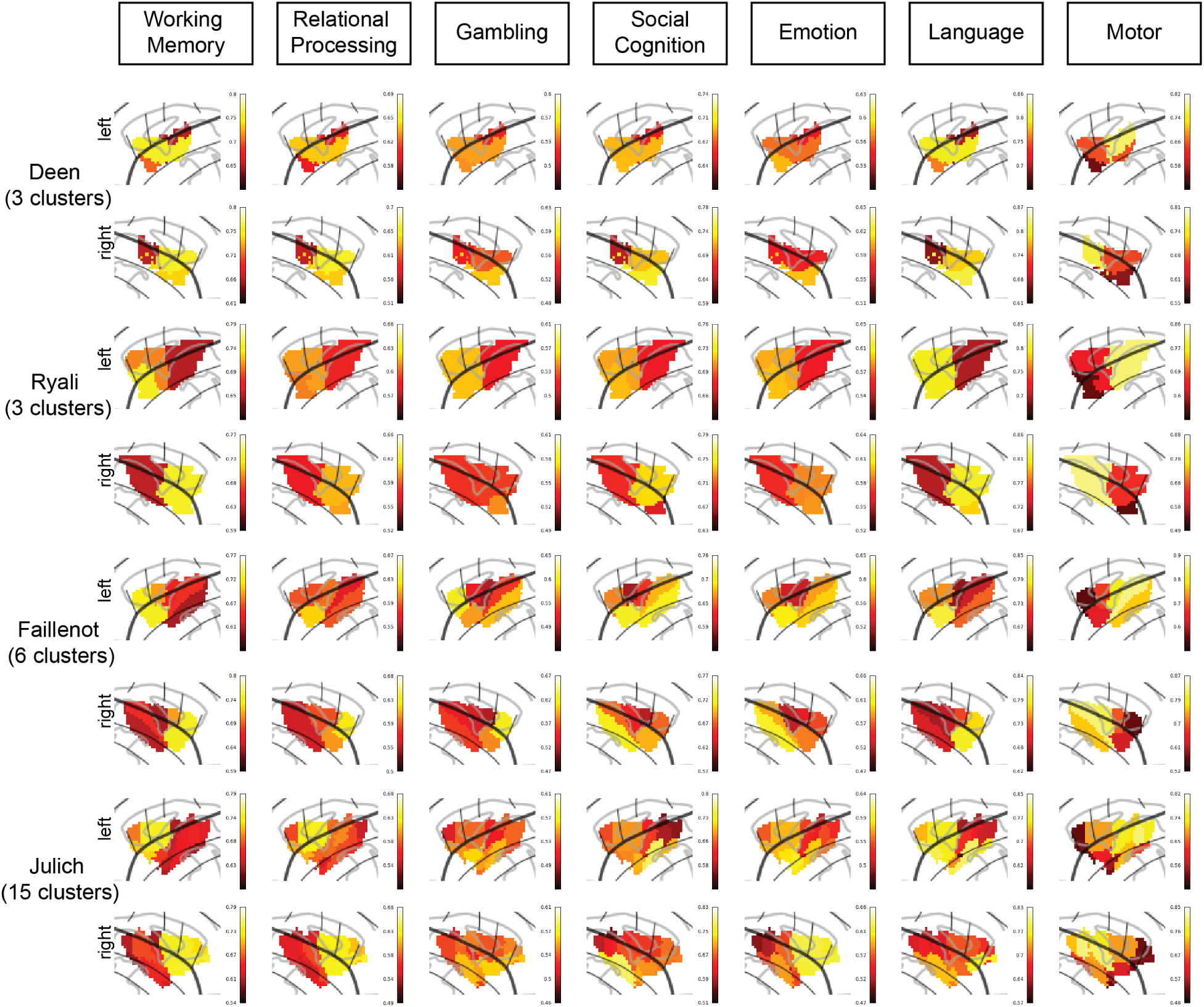
Multivariate activation patterns within individual insular subregions encode task-specific information. Heat map displaying classification accuracy for discriminating between task conditions based on voxel-wise activation patterns within each insular subregion. Brighter colors indicate higher accuracy. Results reveal a triple dissociation: dorsal anterior insula best discriminates cognitive conditions, ventral anterior insula best discriminates emotional/reward conditions, and posterior insula best discriminates motor conditions.

These results reveal a fundamental organization principle: the dAI specializes in representing cognitive control states, the vAI in encoding affective, reward and social information, and the PI in representing sensorimotor states. Crucially, these results generalize across insular parcellations with different levels of granularity.

### Multi-task condition-specific network connectivity patterns differentiate insular subregions

Having established functional specialization through activation patterns, we next investigated whether task-dependent connectivity profiles could also differentiate insular subregions. Task-dependent connectivity was computed between each insular subregion and 246 regions across the brain, then averaged by network membership using the Brainnetcome atlas (**Supplementary Figure S2**).

LinearSVM classification based on connectivity profiles successfully differentiated insular subregions across all parcellation schemes (all *ps*<0.002, **Figure 6a**), with accuracies substantially exceeding chance levels. This indicates that insular subregions maintain distinct network integration patterns that transcend specific task demands.

**Figure 6.**
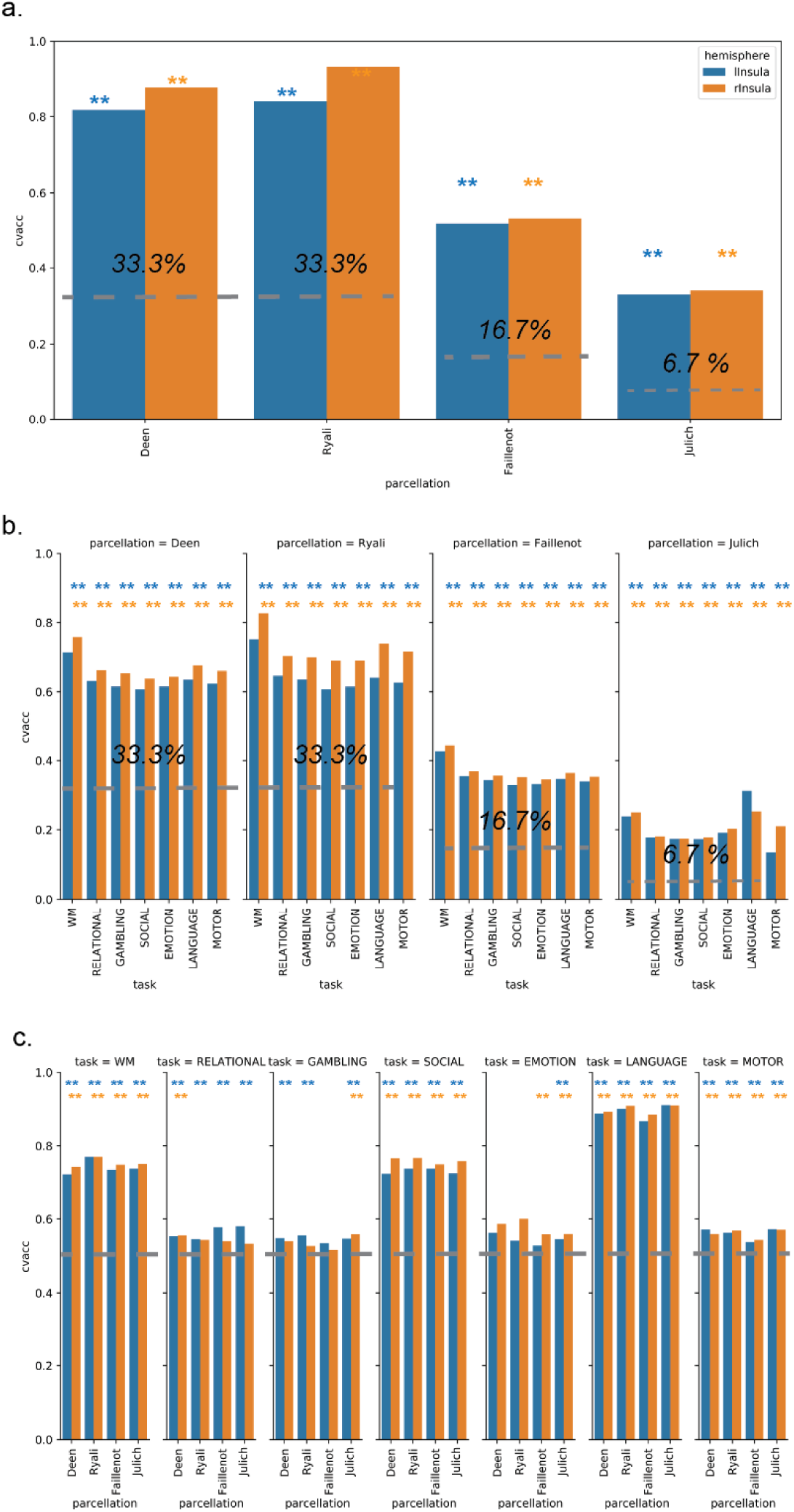
Network connectivity patterns differentiate insular subregions and task conditions. (a) Classification of insular subregions using connectivity patterns across all seven tasks. (b) Within-task classification of insular subregions based on task-specific connectivity patterns. (c) Classification of task conditions using insular-network connectivity patterns. Results reveal particularly high classification accuracy for Working Memory, Social Cognition, and Language tasks. Dashed lines indicate chance levels, and asterisks indicate significant classification results (p<0.05, FWE-corrected).

### Within-task condition-specific network connectivity patterns differentiate insular subregions

When examining individual tasks, connectivity patterns within each task context successfully differentiated insular subregions across all parcellation schemes (all *ps*<0.002, **Figure 6b**).

Notably, classification performance was highest for the working memory task across almost all parcellation schemes, suggesting that cognitive control demands induce particularly distinctive network configurations across insular subregions. This pattern was consistent across parcellation atlases with one exception: connectivity during the language task in the Julich atlas. The superior discriminability during working memory suggests that cognitive control operations may drive the most distinct network reconfiguration among insular subregions.

### Insular-network connectivity encodes task-specific cognitive states

We next investigated whether task-dependent connectivity between insular subregions and brain networks could differentiate between task conditions. Using the same linearSVM framework, we found that connectivity patterns successfully discriminated between task conditions across most cognitive domains (all *ps*<0.002, **Figure 6c**).

Classification accuracy varied systematically across tasks, with highest performance for working memory, social cognition, and language tasks (70-90%) compared to other domains (∼60%). This suggests that these particular cognitive operations induce more distinctive network reconfiguration within the insular cortex, potentially reflecting the insula’s crucial role in integrating cognitive control, social processing, and linguistic functions.

### Task-specific modulation of insula-network connectivity

To characterize the specific network interactions of insular subregions, we examined connectivity between each subregion and major brain networks across task conditions. Given similar classification results across parcellation schemes, we focused on the Deen atlas for detailed network analysis, with results from other atlases reported in Supplementary Materials (**Figures S3-S10, Tables S10-S13**).

Repeated measures ANOVA revealed significant main effects of SUBREGION across most brain networks and tasks (**Supplementary Table S10; Supplementary Figure S3**). The dAI showed significantly stronger connectivity with frontoparietal networks (FPN-1, FPN-2) and salience/ventral attention networks than other subregions during working memory, relational processing, gambling, and language tasks (all *ps*<0.01, FDR corrected, **Figure 7**). The vAI exhibited stronger connectivity with default mode networks (DMN-1, DMN-2) during emotion and motor tasks, while the PI showed stronger connectivity with somatomotor networks and limbic regions, particularly the amygdala-hippocampal complex (all ps<0.01, FDR corrected, **Figure 7**). These distinct connectivity preferences support specialized functional roles that adapt to task context rather than merely reflecting intrinsic connectivity differences.

**Figure 7.**
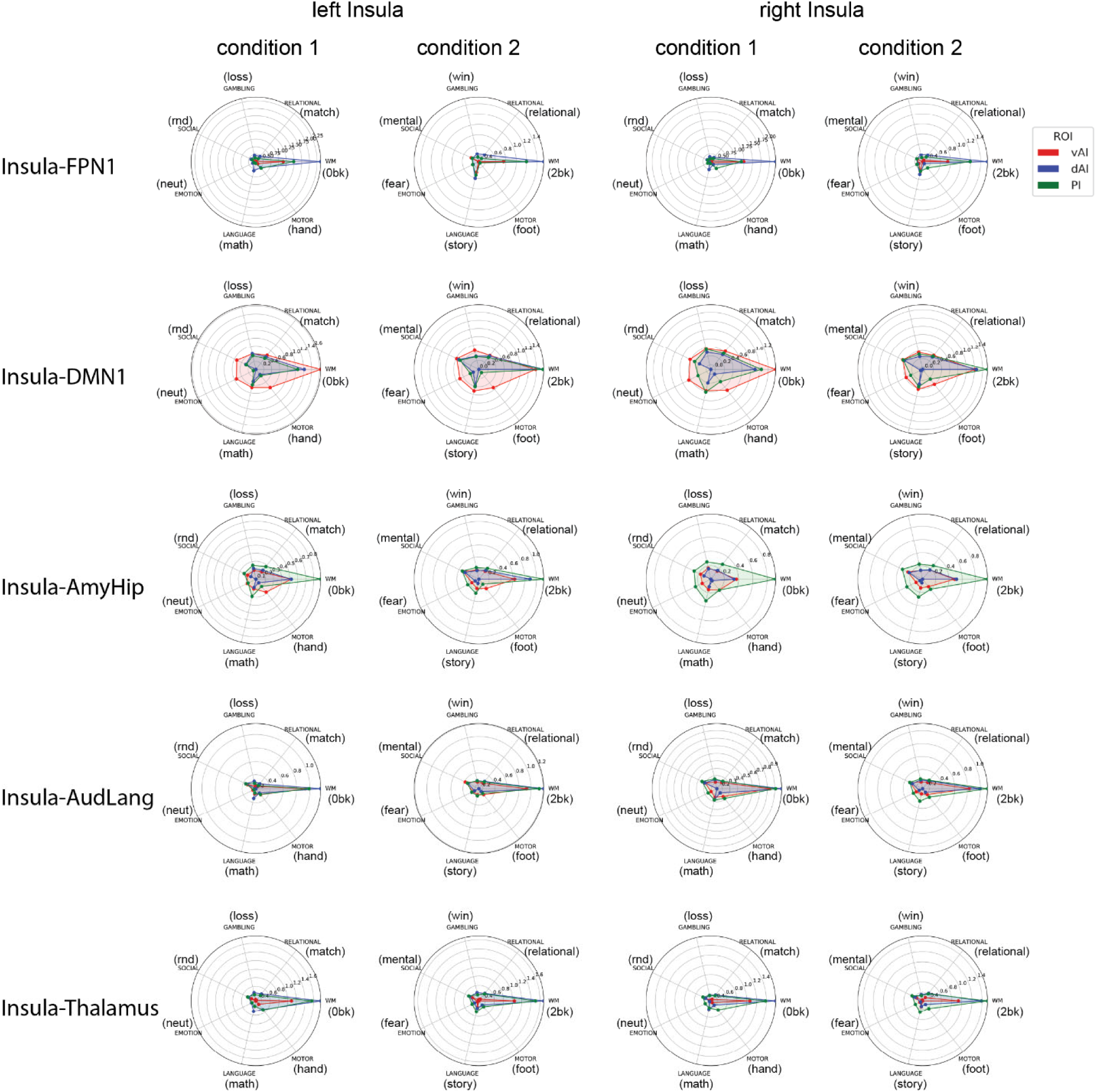
Insular subregions maintain distinct network connectivity preferences across cognitive domains. Task-dependent connectivity patterns between insular subregions (Deen atlas) and selected brain networks: frontoparietal network (FPN-1), default mode network (DMN-1), amygdala-hippocampus complex (AmyHip), auditory-language network (AudLang), and thalamus. Note the consistently stronger connectivity between dorsal anterior insula and frontoparietal regions, and between ventral anterior insula and limbic structures. See Supplementary Figures S3-S4 for complete network connectivity profiles.

Significant SUBREGION × CONDITION interactions revealed differential connectivity modulation by task demands (*p*<0.01, FDR corrected, **Figure 8, Supplementary Figure S3, S4, S11, Supplementary Table S10**). During working memory, significant interactions were observed with frontoparietal, default mode, limbic, dorsal attention, and visual networks. During emotional and social tasks, significant interactions were primarily with default mode and dorsal attention networks, while during language processing, interactions spanned nearly all brain networks. Motor task manipulations selectively modulated somatomotor connectivity. These domain-specific interaction patterns demonstrate how insular subregions selectively reconfigure their network interactions based on cognitive demands.

**Figure 8.**
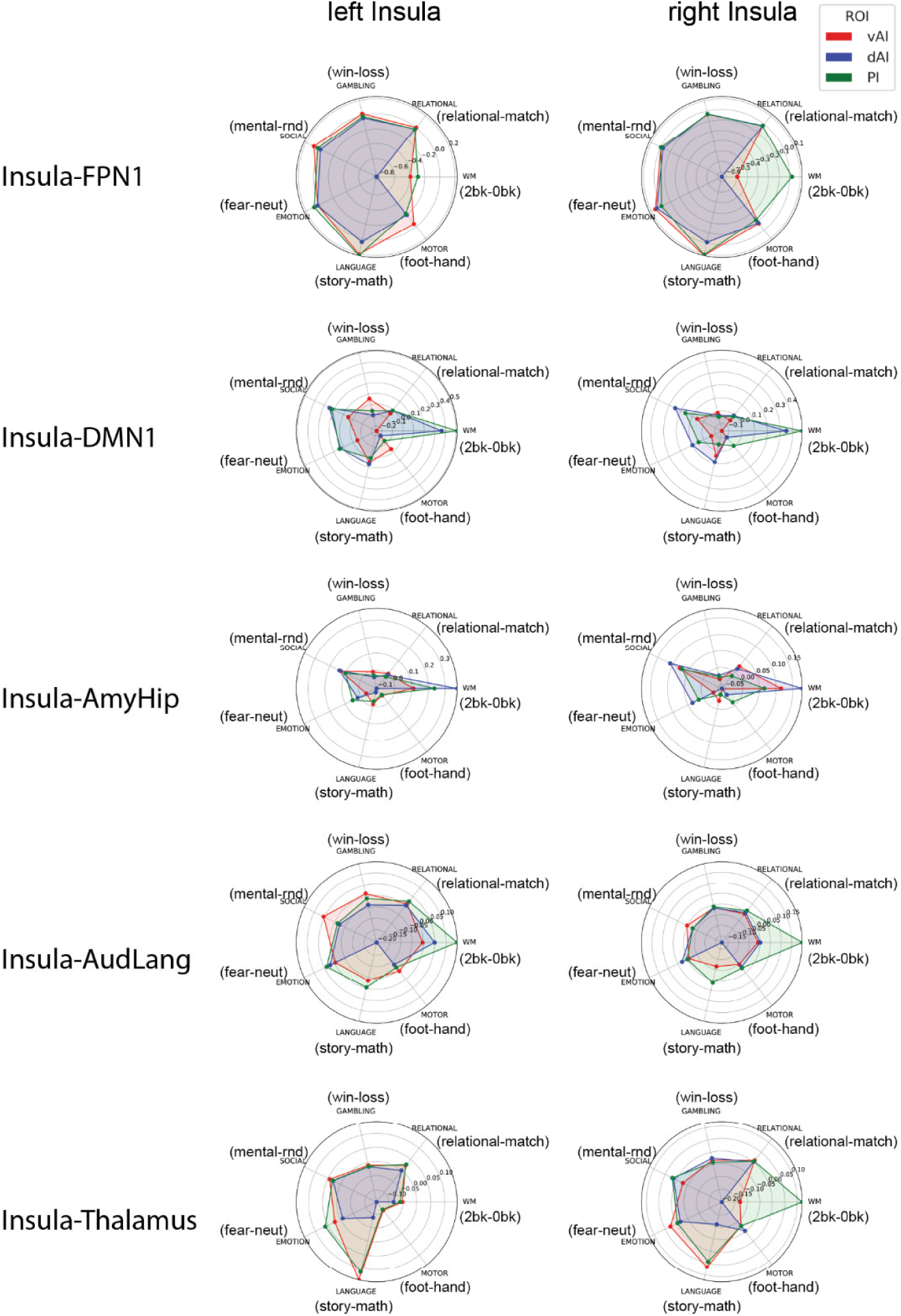
Cognitive demands differentially modulate network connectivity across insular subregions. Contrast values showing task-dependent modulation of connectivity between insular subregions and key brain networks. Note the counterintuitive patterns: during demanding cognitive tasks, dorsal anterior insula shows decreased connectivity with frontoparietal networks but increased connectivity with default mode regions. Ventral anterior insula shows decreased connectivity with limbic regions during emotional processing. These patterns suggest complex information routing mechanisms beyond simple co-activation of task-relevant networks. AudLang = auditory language network; DMN-1= dorsal default mode network; FPN-1 = frontoparietal network; Amy-Hip = amygdala–hippocampus network; and Thalamus.

Follow-up comparisons revealed counterintuitive patterns: during working memory (2-back vs. 0-back) and language tasks (story vs. math), the dAI showed decreased connectivity with frontoparietal networks but increased connectivity with default mode and limbic networks. Conversely, the vAI showed decreased connectivity with default mode networks during working memory, social, and emotion tasks. The vAI’s connectivity with limbic regions, particularly the amygdala-hippocampal complex, was significantly reduced during emotional processing (fear vs. neutral) (all *ps*<0.01, FDR corrected, **Figure 8, Supplementary Figure S3, S4**). Similar results were found for other insular parcellation schemes (**Supplementary Figure S5-10, S12-14**).

These findings reveal task-dependent network reconfiguration patterns that challenge simple interpretations of network engagement. Rather than merely increasing connectivity with task-relevant networks, insular subregions exhibit distinctive patterns of both increased and decreased connectivity that may reflect complex information routing mechanisms supporting diverse cognitive and affective functions.

## Discussion

The insular cortex has long been recognized as a critical neural hub involved in diverse cognitive, emotional, and physiological functions. Our comprehensive analysis using the Human Connectome Project’s multi-task dataset and a within-subjects design reveals five principles of insular functional organization. First, insular subregions maintain distinct functional signatures that enable reliable differentiation based on both activation and connectivity patterns across cognitive domains. Second, these subregions dynamically reconfigure their network interactions in response to specific task demands while preserving their core functional architecture. Third, clear functional specialization exists along the insula’s dorsal-ventral axis, with dorsal regions processing cognitive control information and ventral regions processing emotional and social information. Fourth, counterintuitive connectivity patterns during demanding cognitive tasks suggest complex information routing mechanisms rather than simple co-activation of task-relevant networks. Fifth, while a basic tripartite model captures core functional distinctions, finer-grained parcellations revealed additional cognitive domain-specific advantages that are obscured by simpler parcellation approaches. Collectively, these findings provide a comprehensive framework for understanding how the insula’s structural organization supports its diverse functional roles, with important implications for both basic neuroscience and clinical disorders involving insular dysfunction.

### Distinct functional signatures enable reliable differentiation of insular subregions

Our classification analyses demonstrate that insular subregions maintain distinct functional profiles that transcend specific task contexts. Using linearSVM classifiers, we successfully differentiated insular subregions based on both activation and connectivity patterns with remarkably high accuracy. Cross-task activation patterns yielded classification accuracies of 80% for three-cluster models (Deen, Ryali), 60% for the six-cluster model (Faillenot), and 40% for the fifteen-cluster model (Julich) – all substantially exceeding their respective chance levels (33%, 16.7%, and 6.7%). Similar results were obtained using connectivity patterns, with classification accuracies of 80%, 50%, and 30% for the three-, six-, and fifteen-cluster models, respectively.

Importantly, functional differentiation was evident even within individual task contexts. Classification accuracies remained significantly above chance for all parcellation schemes across all seven tasks, though with some task-dependent variation. The Working Memory, Relational Processing, and Language tasks yielded the highest classification accuracies, while Social Cognition and Emotion tasks showed relatively lower, but still significant, differentiation. This pattern suggests that cognitive control operations may induce more distinctive activation and connectivity patterns across insular subregions than affective processes.

These findings extend previous meta-analytic work^7,33,42,45^ by demonstrating functional specialization within the same individuals across multiple cognitive domains. The consistency of classification performance across parcellation schemes, cognitive domains, and analysis approaches provides compelling evidence that insular subregions maintain distinct functional profiles that reflect fundamental organizational properties rather than methodological artifacts.

### Dynamic network reconfiguration with preserved core functional architecture

Our connectivity analyses reveal that insular subregions maintain distinct network connectivity preferences while dynamically adapting to task demands. Across seven cognitive tasks, insular subregions demonstrated consistent but distinct connectivity profiles: the dorsal anterior insula (dAI) showed preferential connectivity with salience and frontoparietal networks^32,33^, the ventral anterior insula (vAI) exhibited stronger connections with default mode networks, and the posterior insula (PI) maintained strong connectivity with limbic networks, particularly the amygdala-hippocampus complex.

Despite these stable connectivity profiles, we observed task-specific modulation patterns that differed across insular subregions. In particular, the working memory task induced the most distinctive network configurations, yielding the highest classification accuracy for differentiating insular subregions across almost all parcellation schemes. This suggests that cognitive control demands may drive particularly distinct patterns of network reconfiguration in the insula.

Significant interactions between insular subregions and task conditions were observed for specific brain networks in different cognitive domains. During working memory, significant interactions spanned frontoparietal, default mode, limbic, dorsal attention, and visual networks. During emotional and social tasks, significant interactions were primarily with default mode and limbic networks, while during language processing, interactions spanned nearly all brain networks. Motor task manipulations selectively modulated somatomotor connectivity. These domain-specific interaction patterns demonstrate how insular subregions selectively reconfigure their network interactions based on cognitive demands.

This dynamic network organization contrasts markedly with the connectivity patterns observed during resting state^33,42,43,66,71,72^ and reveals broader principles about brain network organization. The ability of the insula to rapidly reconfigure its network interactions while preserving functional specialization balances the competing demands of stability and flexibility – a fundamental organizing principle that may extend beyond the insula to other brain systems.

### Dorsal-ventral functional specialization in the anterior insula

Our multi-task approach provides strong evidence for a fundamental functional dissociation along the dorsal-ventral axis of the anterior insula. The dAI showed selective sensitivity to cognitive control demands, with significantly greater activation during challenging conditions in working memory (2-back vs. 0-back) and relational processing tasks compared to other subregions. In contrast, the vAI demonstrated preferential activation during social processing (mental vs. random) and emotional tasks. This functional dissociation was further supported by our voxel-wise pattern classification analyses. Within the dAI, multivariate patterns achieved highest classification accuracy for discriminating between conditions in working memory, relational processing, and language tasks. In contrast, the vAI showed superior classification performance for distinguishing between conditions in gambling and emotion processing tasks. The posterior insula (PI) exhibited highest classification accuracy for motor task conditions. This triple dissociation provides compelling evidence for distinct computational roles within the insular cortex.

The functional specialization extends to network connectivity patterns. The dAI showed strong task-dependent coupling with frontoparietal networks during cognitively demanding tasks, while the vAI exhibited enhanced connectivity with the default mode network, amygdala, and hippocampus during tasks involving emotional and social processing. These distinct connectivity profiles proved crucial for distinguishing between task conditions, with dAI-frontoparietal connectivity patterns strongly differentiating cognitive control conditions and vAI-limbic connectivity patterns differentiating emotional and social conditions.

This dorsal-ventral dissociation aligns with theoretical models and extends previous structural and intrinsic connectivity findings^31,32,42,54-56,71^. The dAI’s role in cognitive control is consistent with its strong connections to dorsal anterior cingulate cortex and dorsolateral prefrontal cortex, while the vAI’s involvement in affective processing aligns with its connections to ventral anterior cingulate cortex and limbic structures. Importantly, our findings are supported by recent causal evidence from a neuromodulation study showing that transcranial ultrasound stimulation over dAI, but not vAI, impacts cognitive control functions ^73^. This convergence of functional, connectivity, and causal evidence confirms that the dorsal-ventral specialization reflects fundamental organizational properties rather than merely correlational differences.

### Counterintuitive connectivity patterns suggest complex information routing mechanisms

A surprising finding was the pattern of connectivity modulation during demanding cognitive tasks. Contrary to simple models of network engagement, we observed complex, sometimes counterintuitive patterns of connectivity that challenge traditional interpretations. During working memory, the dAI showed increased activation during the more demanding 2-back condition but decreased connectivity with frontoparietal networks compared to the 0-back condition. Simultaneously, the dAI’s connectivity with default mode and limbic networks increased with higher cognitive load – a pattern also observed previously^65^. Similar paradoxical patterns were observed for the vAI, which showed decreased connectivity with default mode networks during tasks requiring enhanced cognitive control (2-back vs. 0-back) and emotional processing (fear vs. neutral). These findings stand in contrast to simplistic models that would predict increased connectivity with task-relevant networks during demanding conditions.

This dissociation between activation and connectivity patterns illuminates an important principle of network organization. During the 0-back condition, which primarily requires external attention and stimulus detection, the dAI maintains strong connectivity with frontoparietal networks. However, during the 2-back condition, which requires maintained internal representations, the dAI reduces frontoparietal connectivity while increasing connectivity with default mode regions. This suggests a fundamental shift in network architecture from a configuration optimized for external stimulus processing to one specialized for internal representation maintenance^74^.

Rather than conceptualizing connectivity as simply increasing or decreasing with task demands, our results suggest complex patterns of network reconfiguration that may reflect strategic routing of information. During high cognitive load, the dAI may decrease frontoparietal connectivity to allow more independent operations of these networks while increasing connectivity with default mode regions to suppress task-irrelevant processing. This interpretation aligns with dynamic causal brain circuit models and suggests that effective cognitive function depends not just on activating appropriate networks but on establishing specific patterns of communication between them.

These findings have important implications for understanding brain network dynamics more broadly. They suggest that cognitive processes rely on complex, context-dependent reconfiguration of network interactions rather than stable, task-invariant connectivity patterns. Further research with high temporal resolution methods will be crucial for testing these hypotheses and clarifying the mechanisms underlying these complex network dynamics.

### Consistency across parcellation schemes indicates fundamental organizational principles

A key strength of our study is the systematic comparison of functional organization across four established insular atlases: Deen and Ryali (3 clusters), Faillenot (6 clusters), and Julich (15 clusters). These insular atlases were generated based on distinct neurobiological features: intrinsic connectivity (Deen, Ryali), morphological characteristics (Faillenot), and cytoarchitecture (Julich). Despite these different anatomical definitions, we observed remarkably consistent functional organization across parcellation schemes.

The dorsal-ventral functional gradient was preserved across atlases, with regions corresponding to the dAI in Deen and Ryali atlases showing similar activation and connectivity patterns to the ASG in Faillenot and Id7/Id6 regions in Julich. Similarly, the differential responses to cognitive control, emotional, and motor tasks were consistent across parcellation schemes. This cross-atlas consistency strongly suggests that our findings reflect fundamental organizational principles rather than artifacts of specific parcellation approaches.

While classification accuracy predictably decreased with more fine-grained parcellations, all schemes showed above-chance performance in differentiating functional subregions. This suggests that even fine-grained anatomical distinctions reflect meaningful functional boundaries. However, the additional granularity of finer parcellations showed domain-specific benefits. The Julich atlas with 15 clusters demonstrated particular advantages in differentiating social, emotional, and motor processes, while showing comparable performance to simpler models for core cognitive tasks.

This pattern suggests a hierarchical organization where broad functional distinctions (captured by the tripartite model) are complemented by finer-grained specialization for specific domains. The basic tripartite model appears sufficient for investigating cognitive control functions and network-level interactions, while more fine-grained parcellations may be particularly valuable for studies focusing on social, emotional, or motor processes. This finding has important methodological implications, suggesting that researchers should select parcellation granularity based on their specific research questions and cognitive domains of interest.

The consistency of our findings across different anatomical definitions strengthens the generalizability of our conclusions and suggests that the functional principles we identified reflect fundamental properties of insular organization rather than methodological artifacts. This multi-atlas approach represents an important methodological advance that could be applied to other brain regions with complex anatomical and functional organization.

### Conclusions and future directions

Our systematic investigation of insular function across multiple cognitive domains reveals five fundamental principles of organization that advance our understanding of this critical brain region. The insula exhibits clear functional specialization implemented through selective engagement with different brain networks. The dorsal anterior insula coordinates cognitive control through dynamic interactions with frontoparietal networks, while the ventral anterior insula integrates emotional and social information through connections with limbic and default mode networks. This functional architecture is optimized for flexible network integration, allowing insular subregions to maintain their core preferences while dynamically reconfiguring network interactions based on task demands.

These findings have important implications for understanding clinical disorders involving insular dysfunction. Many neuropsychiatric conditions – including anxiety, depression, ADHD, autism, and schizophrenia—show altered insular function^15-29^. Our findings suggest that these disorders might differentially affect specific insular subregions or disrupt the dynamic balance of network interactions rather than causing global insular dysfunction. Future research should extend our approach to clinical populations, examining whether specific disorders show characteristic patterns of disruption in insular organization.

Our findings illuminate how the insula’s structural organization supports its diverse functional roles through selective engagement of distinct neural networks. The ability to reconfigure network interactions while maintaining functional specialization may represent a fundamental principle of brain organization that extends beyond the insula, with important implications for understanding both normal cognitive function and the network pathophysiology underlying clinical disorders.

## Methods

### Ethics statement

Data acquisition for the Human Connectome Project (HCP) was approved by the Institutional Review Board of The Washington University in St. Louis ^75^.

### Human Connectome Project (HCP) dataset

HCP dataset ^75^ included 1200 subjects, completing 7 different cognitive tasks within MRI scanner: (1) n-back working memory task; (2) relational processing task; (3) gambling task; (4) language task; (5) social cognition task; (6) emotion processing task; and (7) motor task. Data selection was based on the following criteria: (1) range of head motion in any translational and rotational direction is less than 1 voxel; (2) average scan-to-scan head motion is less than 0.25 mm; (3) each task performance above the criteria (see below); and (4) subjects are right-handed. To ensure that subjects are engaged in each cognitive task, we set performance criterion for each task: WM_Task_Acc > 50%; Relational_Task_Acc > 50%; Language_Task_Acc > 50%; Emotion_Task_Acc > 50%; Social_Task_Perc_Tom > 25%; Gambling_Task_Perc_Larger > 25%; no criteria used in Motor task. Our final sample included 524 subjects (22-36 years old).

Each task contains two major conditions. The n-back working memory task has the 0-back and 2-back conditions. The relational processing task has the match and relational processing conditions. The gambling task has the loss and win conditions. The language task has the story and math conditions. The social cognition task has the random and mental conditions. The emotion processing task has the neutral and fear conditions. The motor task has the foot and hand conditions. All the task uses mini-block designs. Task design details are described in the Supplementary Methods.

### fMRI acquisition

HCP fMRI data was acquired using the multiband, gradient-echo planar imaging with the following parameters: TR=720 ms; TE=33.1 ms; flip angle=52°; and in-plane resolution=2 mm^75^.

### fMRI preprocessing

We downloaded minimally preprocessed fMRI data, which was preprocessed using *fMRIVolume pipeline* ^76^, including correction of gradient-nonlinearity-induced distortion, realignment for motion correction, registration, and normalization in 2mm MNI space. Details of the preprocessing steps are described in previous studies ^76^. We applied spatial smoothing with a Gaussian kernel of 6mm FWHM in the minimally preprocessed data to improve signal-to-noise ratio as well as anatomical correspondence between individuals.

### Task-dependent activation analysis

A general linear model (GLM) analysis was used to determine task-dependent activation in each of seven cognitive control tasks ^77^. Six motion parameters were entered as covariates of no interest, and both canonical hemodynamic response function (HRF) and its time-derivative were used to convolve the stimulus function to form task regressors. Two main task conditions were modelled in the GLM in each cognitive task. The Working Memory task had the 0-back and 2-back conditions; the Relational Processing task had the match and relational processing conditions; the Gambling task had the win and loss conditions; the Language task had the story and math conditions; the Social Cognition task had the mental and random conditions; the Emotion Processing had the fear and neutral conditions; and the Motor task had the hand and foot conditions.

### Task-dependent connectivity analysis

Seed-based generalized psychophysiological interaction (gPPI) was used to determine task-dependent functional connectivity ^78^. Seeds were placed in the insular subregions. The gPPI model consisted of a physiological variable (the raw time series of a seed), multiple psychological variables (hemodynamic response function convolved main effect of condition of interest), and multiple interaction variables (deconvolved raw time series of the seed multiplied by main effect of condition of interest, and then convolved with the hemodynamic response function). Task-dependent connectivity was computed between each insular subregion and regions of interest from the rest of the brain.

### Multi-task condition-specific activation patterns differentiate insular subregions

We conducted a classification analysis to determine whether multi-task condition-specific activation patterns can differentiate insular subregions. Using a linear support vector machine (linearSVM) algorithm (C=1), we trained a model to predict subregion labels (e.g. dAI vs. vAI vs. PI). For each subregion (e.g. dAI), the input features consist of condition-specific mean activation values across seven tasks, yielding a feature vector of 14 activation values (7 tasks x 2 conditions per task).

Model performance was evaluated using 5-fold cross-validation. The dataset was randomly partitioned into five folds, with one-fold serving as the test set while the remaining folds were used for training. This process was repeated five times, ensuring that each fold served as the test set once. Classification accuracies were averaged across all the five test folds to obtain the final cross-validation accuracy.

To assess the statistical significance of model performance, we conducted a permutation test. In each permutation, subregion labels were randomly shuffled, and the linearSVM model was trained on the permuted data, applying the same 5-fold cross-validation procedure. This procedure was repeated 500 times to construct an empirical null distribution of classification accuracies, from which a p value for the observed classification accuracy was derived.

### Within-task condition-specific activation patterns differentiate insular subregions

We further examined whether condition-specific activation patterns within individual cognitive task could differentiate insular subregions. Using the same linearSVM algorithm (C=1), we trained a model to predict subregion labels (e.g. dAI, vAI and PI). For each insular subregion (e.g. dAI), input features consisted of condition-specific mean activation values within a given task, yielding a two-value feature vector (e.g. 0-back and 2-back conditions in the WM task). The same cross-validation procedure and permutation approach were applied to evaluate model performance.

### Insular activation patterns encode task-specific cognitive states

We investigated whether insular activation patterns contained sufficient information to discriminate between cognitive conditions within each task. Using the same linear SVM algorithm (C=1), we trained a model to predict task conditions (e.g. 0-back versus 2-back in the WM task). For each task condition (e.g. 0-back), the input features comprised subregion-specific mean activation values across all insular subregions. The number of input features corresponded to the number of parcels the insular atlas (e.g., 3 for Deen and Ryali, 6 for Faillenot, and 15 for Julich). The same cross-validation and permutation testing methods were implemented to assess model performance.

### Multivariate patterns within individual insular subregions encode task-specific information

We extended our analysis to test whether voxel-wise activation patterns within each insular subregion could differentiate task conditions for each cognitive task. In this analysis, voxel-wise activation values were extracted from each insular subregion per task condition and used as features in the linearSVM model for task condition classification within each task (e.g. 0-back vs. 2-back). Model performance was evaluated using the same cross-validation procedure.

### Multi-task condition-specific network connectivity patterns differentiate insular subregions

Next, we examined whether task-dependent subregion-to-network connectivity could differentiate insular subregions. Again, we employed the same linearSVM algorithm (C=1) to train a classification model to predict subregion labels (e.g. dAI vs. vAI vs. PI). For each subregion (e.g. dAI), the input features consist of condition-specific gPPI values of connectivity between each insular subregion and each brain network, resulting a vector of 280 connectivity values (7 tasks x 2 conditions per task x 20 brain networks). The same cross-validation and permutation testing methods were applied to evaluate model performance.

### Within-task condition-specific network connectivity patterns differentiate insular subregions

We further explored whether condition-specific subregion-to-network connectivity patterns within individual cognitive task could differentiate insular subregions. Using the same linearSVM algorithm (C=1), we trained a model to predict subregion labels (e.g. dAI, vAI and PI). For each insular subregion (e.g. dAI), the input features consisted of condition-specific subregion-to-network connectivity within each task, yielding a vector of 40 gPPI values (e.g. 2 task conditions x 20 brain networks). The same cross-validation and permutation testing methods were applied to assess model performance.

### Insular network connectivity patterns encode task-specific cognitive states

Finally, we tested whether insula-to-network connectivity patterns contain sufficient information to discriminate between cognitive conditions within each task. Using the same linearSVM algorithm (C=1), we trained a model to predict task conditions (e.g. 0-back versus 2-back in the WM task). For each task condition (e.g. 0-back), the input features consisted of condition-specific subregion-to-network connectivity patterns across all the subregions. The number of input features was determined by the product of the number of the insular atlas parcels and the number of brain networks (for instance, 60 for Deen and Ryali atlas, 120 for Faillenot atlas and 300 for Julich). The same cross-validation and permutation testing approach were employed to assess model performance.

### Insular atlases and parcellation schemes

Four different insular atlases were tested in this study, including Deen, Ryali, Faillenot and Julich. Deen and Ryali had 3 clusters in each hemisphere of the insular cortex. Both atlases have the vAI, dAI and PI subregions. Faillenot had 6 clusters in each hemisphere of the insular cortex, including anterior short gyrus (ASG), middle short gyrus (MSG), posterior short gyrus (PSG), anterior inferior cortex (AIG), anterior long gyrus (ALG), posterior long gyrus (PLG). The Julich had 15 clusters in each hemisphere of the insular cortex, involving Area Ia1, Ia2, Ia3, Id1, Id2, Id3, Id4, Id5, Id6, Id7, Id8, Id9, Ig1, Ig2 and Ig3. **Figure 1a** illustrates the insular parcellation of each atlas. More details were provided in the **Supplementary Results**.

### ROI correspondence

To determine ROI correspondence between insular atlases, we computed a voxel overlapping index. Because of different anatomical templates were used when generating parcellations, the four insular atlases were not completely overlapped. To estimate the extent to which two ROIs are overlapped, the voxel overlapping index was calculated as the ratio of the number of voxels in ROI_A and in ROI_B versus the number of voxels in ROI_A or in ROI_B. The ROI correspondence were reported in the **Supplementary Results** and **Supplementary Table S1**

## Data availability

The task fMRI data is accessible from the HCP database (https://db.humanconnectome.org/).

## Code availability

Functional MRI data analyses were conducted using SPM12 on Matlab R2022b. Classification analyses were conducted using scikit-learn 1.6.1 (https://scikit-learn.org/) and figures were plotted using seaborn 0.13.2 (https://seaborn.pydata.org/) and Nilearn (https://nilearn.github.io/). Data analyses scripts can be accessed at Github (https://github.com/scsnl/Cai_2025_Insula_HCP_gPPI).

## Acknowledgements

This research was supported by National Institutes of Health MH124816 (WC), EB022907 (VM), NS086085 (VM), and MH121069 (VM).

## Author Contributions

Conceptualization: W.C., V.M.; Methodology: W.C.; Investigation: W.C., Writing: W.C., V.M.; Review & Editing: W.C., V.M.

