## Supplementary Material for "Heterogeneity of human insular cortex: Five principles of functional organization across multiple cognitive domains"

### Heterogeneity of human insular cortex: from task-dependent activation and connectivity to cognitive functions

[1]Department of Psychiatry and Behavioral Sciences
[2]Wu Tsai Neurosciences Institute
[3]Wu Tsai Human Performance Alliance
[4]Maternal & Child Health Research Institute
[5]Department of Neurology & Neurological Sciences

Stanford University School of Medicine
Stanford, CA 94305, USA

### I. Supplementary Methods

#### HCP dataset

The HCP dataset has seven different cognitive tasks: (1) n-back working memory task, (2) relational processing task, (3) gambling task, (4) language task, (5) social cognition task, (6) emotion processing task, and (7) motor task.

**N-back working memory (WM) task** The WM task combines the category specific representation task and the *n*-back working memory task in a single task across two sessions. Participants were presented with blocks of trials that consisted of pictures of faces, places, tools and body parts. Within each session, the 4 different stimulus types were presented in separate blocks. Furthermore, within each session, half of the blocks are 2-back working memory and half are 0-back working memory task. In the 2-back working memory task blocks, subjects were requested to determine whether the current stimulus matches the stimulus in two presentations of stimuli prior within the same block. In the 0-back working memory task blocks, subjects were requested to determine whether the current stimulus matches the target that was presented in the beginning of each block (cue). A 2.5 second cue indicated the task type (and target for 0-back task) at the beginning of each block. Each task session contained 8 task blocks (10 trials of 2.5 seconds each, for 25 seconds) and 4 fixation (“rest”) blocks (15 seconds). On each trial, the stimulus was presented for 2 seconds, followed by a 0.5 second inter-trial-interval (ITI).

**Relational processing (Relational) task** The Relational task involved a relational processing task and a control condition. In the relational processing condition, participants were presented two pairs of objects, with one pair at the top of the screen and the other pair at the bottom. Participants were required to determine whether the top pair of objects and the bottom pairs of objects are different in the same dimension or not. For example, the top pair differed in texture and the bottom pair differs in shape. In the control task, there were two objects presented at the top of the screen, one object at the bottom of the screen and a word in the middle to indicate dimension (e.ge. “shape” or “texture”). Participants were told to judge whether the bottom object matches either of the top objects in that dimension. Each condition had 3 blocks and each block lasts 18 seconds. There were 4 trials in each relational processing block and 5 trials in each control matching block. In the relational condition, the stimuli were presented for 3500 ms, with a 500 ms ITI. In the matching condition, the stimuli were presented for 2800 ms, with a 400 ms ITI.

**Gambling (Gambling) task** The Gambling Task involves a card-guessing game where participants aim to predict the number on a mystery card (represented by a "?") to win or lose money. Participants are informed that the card numbers range from 1 to 9 and must decide whether the mystery card's number is greater or less than 5 by pressing one of two buttons on a response box. Feedback reveals the actual card number (generated based on whether the trial is classified as a reward, loss, or neutral trial) and is accompanied by one of the following outcomes: (1) A green upward arrow with "$1" for reward trials, (2) A red downward arrow with "-$0.50" for loss trials, and (3) The number 5 with a gray double-headed arrow for neutral trials. The mystery card ("?") is displayed for up to 1500 milliseconds (ms). If the participant responds before the allotted time, a fixation cross replaces the mystery card for the remainder of the duration. Feedback is then displayed for 1000 ms, followed by a 1000 ms inter-trial interval (ITI) marked by a "+" on the screen. The task is organized into blocks of 8 trials, which are either predominantly reward-based (6 reward trials intermixed pseudo-randomly with 1 or 2 neutral or loss trials) or predominantly loss-based (6 loss trials intermixed pseudo-randomly with 1 or 2 neutral or reward trials). Each run consists of 2 mostly reward and 2 mostly loss blocks, interspersed with 4 fixation blocks lasting 15 seconds each. The task spans two runs in total.

**Social cognition (Social) task** The Social Task presents participants with 20-second video clips depicting objects (squares, circles, and triangles) that either interact meaningfully or move randomly on the screen. After watching each video, participants are asked to evaluate whether the objects were engaged in a mental interaction (appearing to consider each other's thoughts and feelings), showed no interaction (movement appears random), or uncertain (Not Sure). Each task run includes 5 video blocks and 5 fixation blocks (15 seconds each). One run contains 2 Mental Interaction videos and 3 Random videos, and the other run contains 3 Mental Interaction videos and 2 Random videos.

**Emotion (Emotion) task** The Emotion Task requires participants to complete blocks of trials where they match either faces or shapes. In face-matching trials, participants decide which of two faces at the bottom of the screen matches the face at the top, with the faces displaying either angry or fearful expressions. In shape-matching trials, participants select the shape at the bottom that matches the shape at the top. Trials are grouped into blocks of 6 trials, each focusing on a single task type (faces or shapes). Each trial consists of a stimulus displayed for 2000 milliseconds (ms), followed by a 1000 ms inter-trial interval (ITI). A 3000 ms task cue ("shape" or "face") precedes each block, making the total block duration 21 seconds. Each of the two runs includes 3 face blocks and 3 shape blocks, with an additional 8-second fixation period at the end of each run.

**Language (Language) task** The Language Task comprises two runs, each alternating between 4 blocks of a story task and 4 blocks of a math task. Block durations vary, averaging approximately 30 seconds, but the math task blocks are matched in length to the story task blocks. Additional math trials are included at the end of each run, as needed, to complete the 3.8-minute duration. In the story task, participants listen to brief auditory stories (5–9 sentences) adapted from Aesop's fables. Following each story, participants answer a two-alternative forced-choice question about the story's topic. In the math task, participants hear addition or subtraction problems and select the correct answer by pressing a button corresponding to one of two options. The math task is adaptive, adjusting its difficulty to maintain a consistent challenge across participants.

**Motor (Motor) task** The Motor Task assesses motor function by requiring participants to perform specific movements: tapping their left or right fingers, squeezing their left or right toes, or moving their tongue. Each movement block lasts 12 seconds and consists of 10 repetitions of the assigned movement, preceded by a 3-second cue signaling the upcoming action. Each of the two task runs includes 13 movement blocks, including 2 blocks of tongue movements, 4 blocks of hand movements (2 left-hand and 2 right-hand), and 4 blocks of foot movements (2 left-foot and 2 right-foot). Additionally, each run incorporates 3 fixation blocks, each lasting 15 seconds, to serve as a baseline for comparison.

#### Supplementary Results

*Insular atlases*

While some studies found 3 clusters in the insular cortex, involving vAI, dAI and PI, others suggested more fine-grained parcellation. To test the robustness of our findings, we used multiple established insular atlases with the number of clusters in each hemisphere ranging from 3 to 15, including Deen [36], Ryali [63], Faillenot [39] and Julich [38]. The Last Name of the first author of the corresponding study is used here as the name of the insular parcellation except the Julich.

Deen and Ryali parcellations had 3 clusters in each hemisphere of the insular cortex, all including the vAI, dAI and PI. In both Deen and Ryali's studies, insular parcellation were determined by intrinsic connectivity derived from resting-fMRI data. Faillenot parcellation had 6 clusters in each hemisphere of the insular cortex, including anterior short gyrus (ASG), middle short gyrus (MSG), posterior short gyrus (PSG), anterior inferior cortex (AIG), anterior long gyrus (ALG), posterior long gyrus (PLG). Segmentation between insular subregions in Faillenot's study was based on gyral and sulcal characteristics of insula and neighboring structures. The Julich had 15 clusters in each hemisphere of the insular cortex, involving Area Ia1, Ia2, Ia3, Id1, Id2, Id3, Id4, Id5, Id6, Id7, Id8, Id9, Ig1, Ig2 and Ig3. The Julich atlas was built on human brain's cytoarchitecture. **Figure 1a** illustrates the insular parcellation of each atlas.

*ROI correspondence*

To test whether task-dependent activation or connectivity patterns remain stable across different parcellations or atlases, we need to first determine the spatial correspondence between ROIs across atlases. We developed a ROI overlapping index, which calculates the proportion of voxels overlapped between two ROIs. Then the best matching ROIs between two atlases were the ones with the highest ROI overlapping index (**Supplementary Table S1**).

The vAI, dAI and PI in both hemispheres are well matched between Deen and Ryali.

The left vAI and dAI in both Deen and Ryali were matched to AIC and ASG in Faillenot, respectively; the left PI in Deen was matched to PSG whereas the left PI in Ryali were matched to ALG in Faillenot. The right vAI, dAI, and PI in both Deen and Ryali were matched to AIC in Faillenot, ASG and ALG, respectively.

The left vAI, dAI, and PI in both Deen and Ryali were matched to Id8, Id6, and Id5 in Julich, respectively. The right vAI and dAI in both Deen and Ryali were matched to Id9 and Id6 in Julich, respectively. The right PI in Deen was matched to Ig3 or Id2 and the right PI in Ryali was matched to Ig2 in Julich.

The AIC, ALG, ASG, MSG, PLG, and PSG in both hemisphere in Faillenot were matched to Id9, Id5, Id8, Id6, Ig2 and Id4 in Julich, respectively.

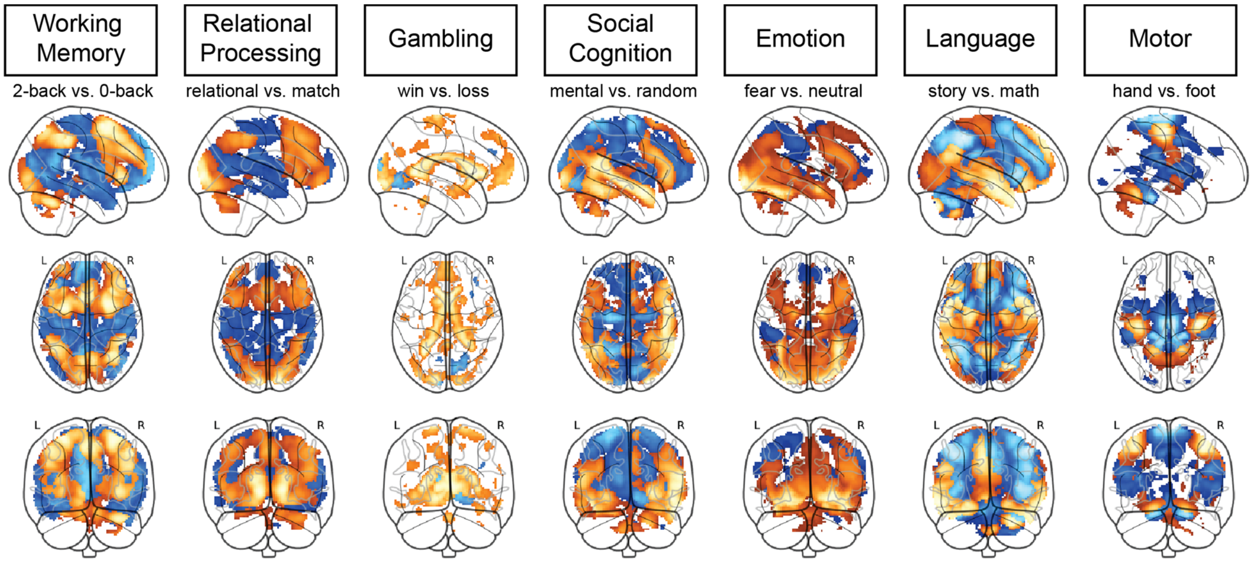

**Figure S1** Illustration of whole brain activation map in each of the 7 HCP tasks. Each general linear model was constructed with the main task conditions (e.g. 2-back and 0-back in the Working Memory task. Hot color represents significantly greater activation in the first than second conditions (e.g. 2-back > 0-back) whereas cool color represents significantly greater activation in the second than first conditions (e.g. 2-back < 0-back) ($p$<0.01, FWE corrected).

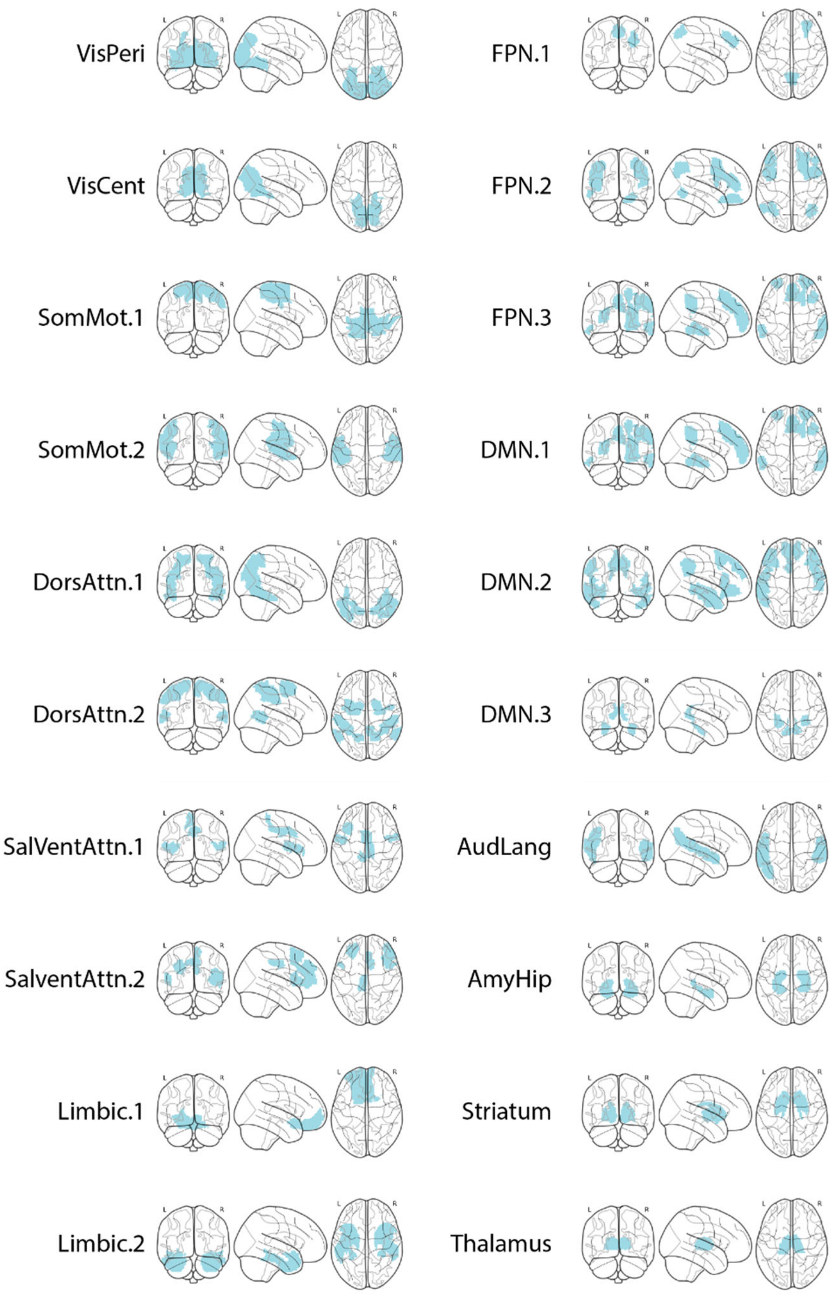

**Figure S2** Illustration of 20 networks in the Brainnetome atlas. The networks were labels as: AudLang = auditory language network; VisCent = visual central network; VisPeri = visual peripheral network; SomMot-1 = somatomotor network; SomMot-2 = somatomotor network; SalVentAttn-1 = salience/ventral attention network; SAlVentAttn-2 = salience/ventral attention network; DMN-1= dorsal default mode network; DMN-2 = ventral default mode network; DMN-3 = default mode network; Limbic-1 = limbic network; Limbic-2 = limbic network; FPN-1 = frontoparietal network; FPN-2 = frontoparietal network; FPN-3 = frontoparietal network; DorsAttn-1 = dorsal attention network; DorsAttn-2 = dorsal attention network; Amy-Hip = amygdala–hippocampus network; Striatum and Thalamus.

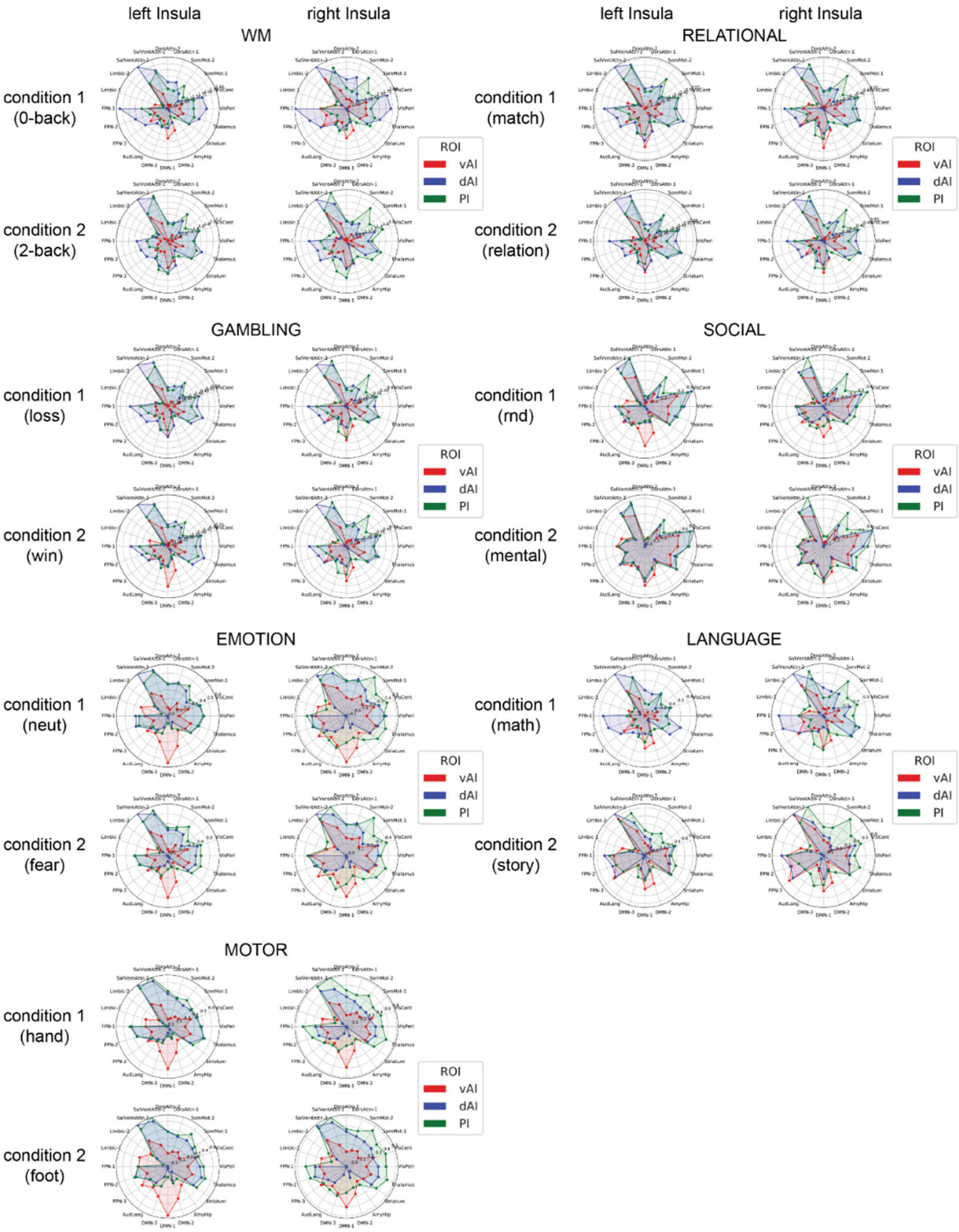

**Figure S3** Illustration of task-dependent connectivity between insular subregions (Deen atlas) and other brain regions (Brainnectome atlas). The insular-brain connectivities were group by brain regions' network membership. The networks were labels as: AudLang = auditory language network; VisCent = visual central network; VisPeri = visual peripheral network; SomMot-1 = somatomotor network; SomMot-2 = somatomotor network; SalVentAttn-1 = salience/ventral attention network; SAlVentAttn-2 = salience/ventral attention network; DMN-1= dorsal default mode network; DMN-2 = ventral default mode network; DMN-3 = default mode network; Limbic-1 = limbic network; Limbic-2 = limbic network; FPN-1 = frontoparietal network; FPN-2 = frontoparietal network; FPN-3 = frontoparietal network; DorsAttn-1 = dorsal attention network; DorsAttn-2 = dorsal attention network; Amy-Hip = amygdala–hippocampus network; Striatum and Thalamus.

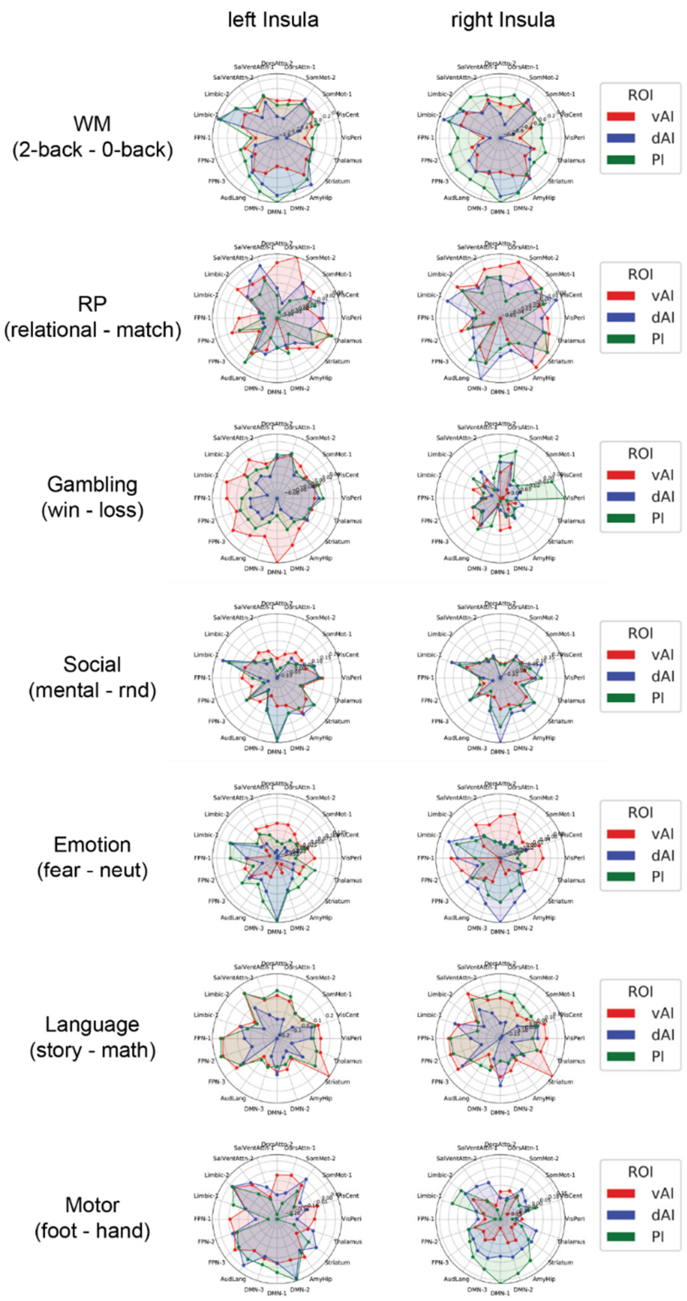

**Figure S4** Illustration of experimental modulation of task-dependent connectivity between insular subregions (Deen atlas) and other brain regions (Brainnectome atlas). The insular-brain connectivities were group by brain regions' network membership. The networks were labels as: AudLang = auditory language network; VisCent = visual central network; VisPeri = visual peripheral network; SomMot-1 = somatomotor network; SomMot-2 = somatomotor network; SalVentAttn-1 = salience/ventral attention network; SAlVentAttn-2 = salience/ventral attention network; DMN-1= dorsal default mode network; DMN-2 = ventral default mode network; DMN-3 = default mode network; Limbic-1 = limbic network; Limbic-2 = limbic network; FPN-1 = frontoparietal network; FPN-2 = frontoparietal network; FPN-3 = frontoparietal network; DorsAttn-1 = dorsal attention network; DorsAttn-2 = dorsal attention network; Amy-Hip = amygdala–hippocampus network; Striatum and Thalamus.

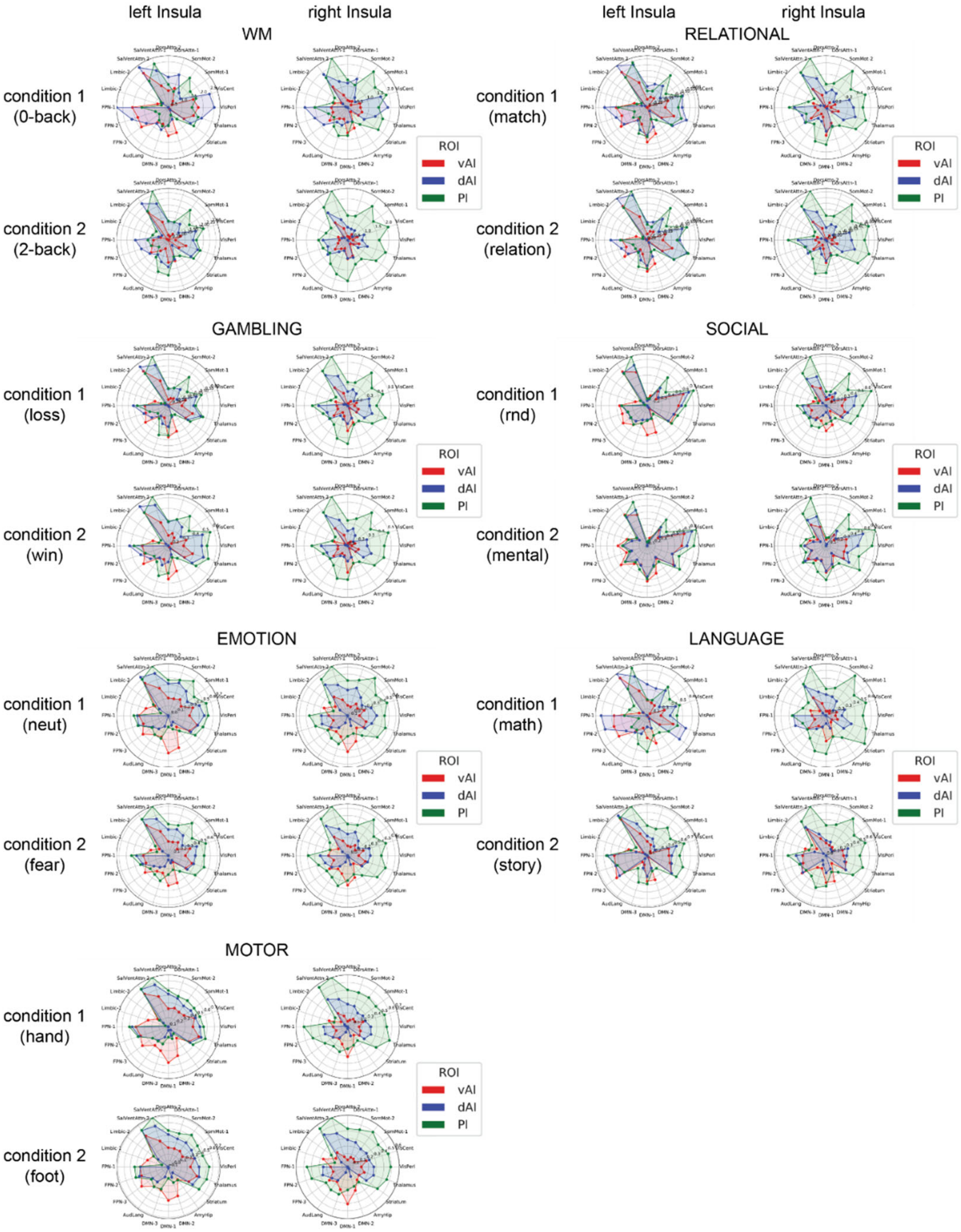

**Figure S5** Illustration of task-dependent connectivity between insular subregions (Ryali atlas) and other brain regions (Brainnectome atlas). The insular-brain connectivities were group by brain regions' network membership. The networks were labels as: AudLang = auditory language network; VisCent = visual central network; VisPeri = visual peripheral network; SomMot-1 = somatomotor network; SomMot-2 = somatomotor network; SalVentAttn-1 = salience/ventral attention network; SAlVentAttn-2 = salience/ventral attention network; DMN-1= dorsal default mode network; DMN-2 = ventral default mode network; DMN-3 = default mode network; Limbic-1 = limbic network; Limbic-2 = limbic network; FPN-1 = frontoparietal network; FPN-2 = frontoparietal network; FPN-3 = frontoparietal network; DorsAttn-1 = dorsal attention network; DorsAttn-2 = dorsal attention network; Amy-Hip = amygdala–hippocampus network; Striatum and Thalamus.

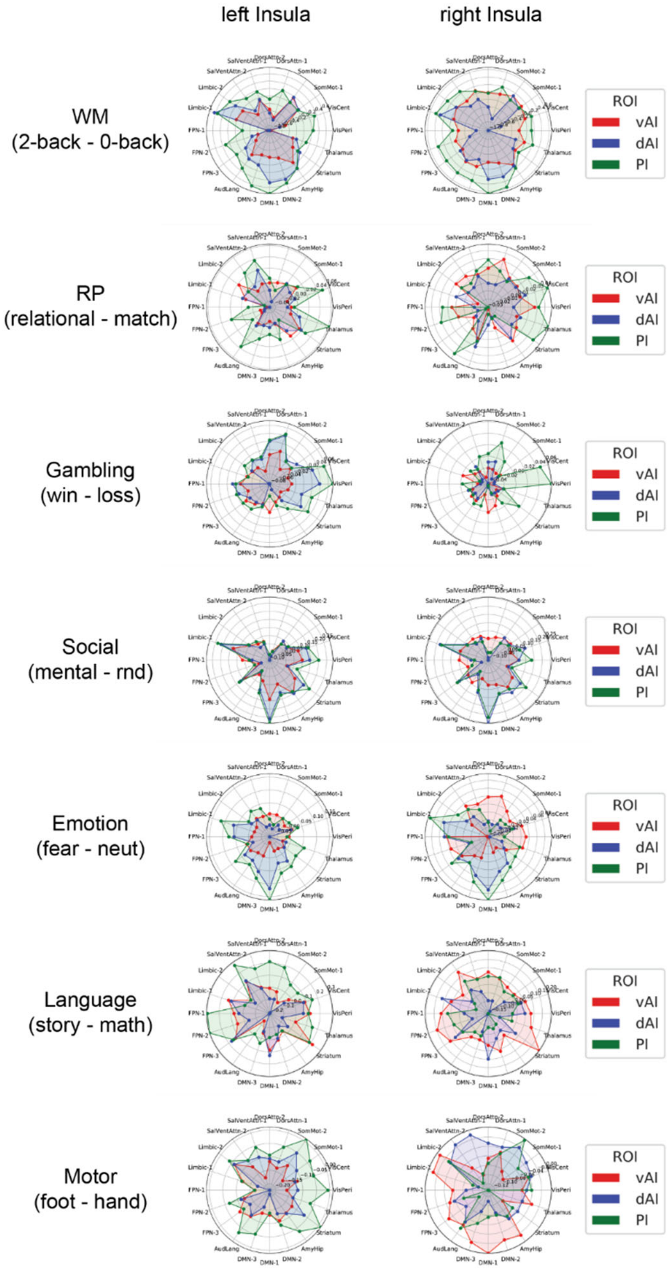

**Figure S6** Illustration of experimental modulation of task-dependent connectivity between insular subregions (Ryali atlas) and other brain regions (Brainnectome atlas). The insular-brain connectivities were group by brain regions' network membership. The networks were labels as: AudLang = auditory language network; VisCent = visual central network; VisPeri = visual peripheral network; SomMot-1 = somatomotor network; SomMot-2 = somatomotor network; SalVentAttn-1 = salience/ventral attention network; SAlVentAttn-2 = salience/ventral attention network; DMN-1= dorsal default mode network; DMN-2 = ventral default mode network; DMN-3 = default mode network; Limbic-1 = limbic network; Limbic-2 = limbic network; FPN-1 = frontoparietal network; FPN-2 = frontoparietal network; FPN-3 = frontoparietal network; DorsAttn-1 = dorsal attention network; DorsAttn-2 = dorsal attention network; Amy-Hip = amygdala–hippocampus network; Striatum and Thalamus.

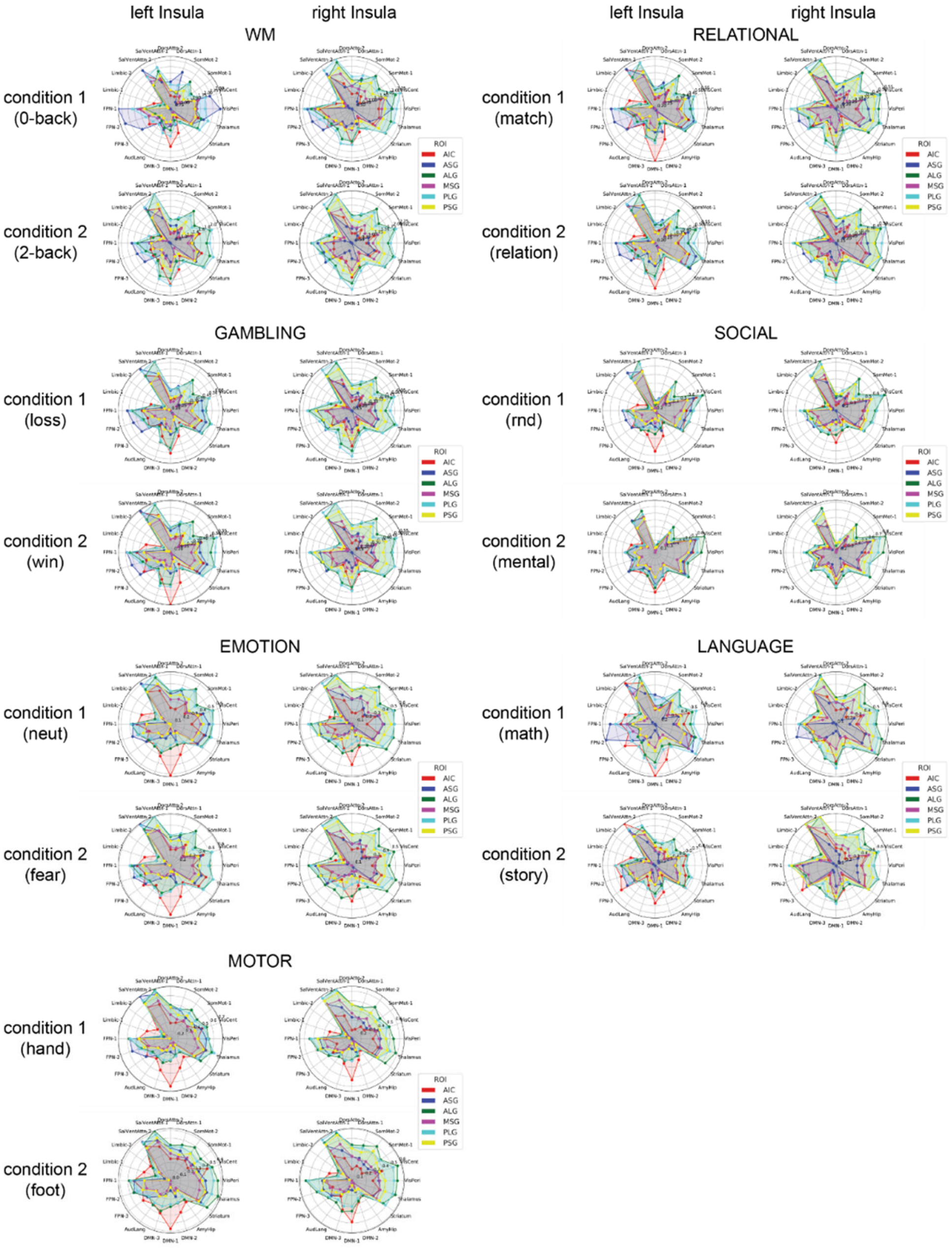

**Figure S7** Illustration of task-dependent connectivity between insular subregions (Faillenot atlas) and other brain regions (Brainnectome atlas). The insular-brain connectivities were group by brain regions' network membership. The networks were labels as: AudLang = auditory language network; VisCent = visual central network; VisPeri = visual peripheral network; SomMot-1 = somatomotor network; SomMot-2 = somatomotor network; SalVentAttn-1 = salience/ventral attention network; SAlVentAttn-2 = salience/ventral attention network; DMN-1= dorsal default mode network; DMN-2 = ventral default mode network; DMN-3 = default mode network; Limbic-1 = limbic network; Limbic-2 = limbic network; FPN-1 = frontoparietal network; FPN-2 = frontoparietal network; FPN-3 = frontoparietal network; DorsAttn-1 = dorsal attention network; DorsAttn-2 = dorsal attention network; Amy-Hip = amygdala–hippocampus network; Striatum and Thalamus.

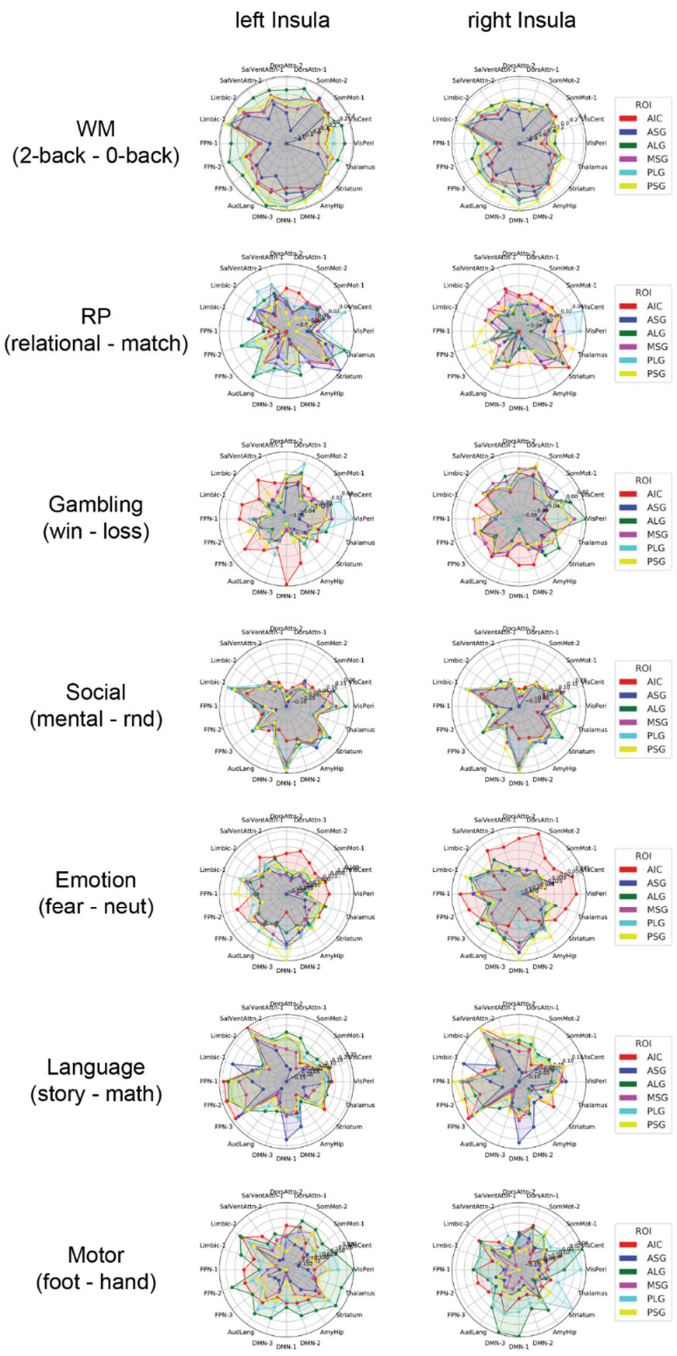

**Figure S8** Illustration of experimental modulation of task-dependent connectivity between insular subregions (Faillenot atlas) and other brain regions (Brainnectome atlas). The insular-brain connectivities were group by brain regions' network membership. The networks were labels as: AudLang = auditory language network; VisCent = visual central network; VisPeri = visual peripheral network; SomMot-1 = somatomotor network; SomMot-2 = somatomotor network; SalVentAttn-1 = salience/ventral attention network; SAlVentAttn-2 = salience/ventral attention network; DMN-1= dorsal default mode network; DMN-2 = ventral default mode network; DMN-3 = default mode network; Limbic-1 = limbic network; Limbic-2 = limbic network; FPN-1 = frontoparietal network; FPN-2 = frontoparietal network; FPN-3 = frontoparietal network; DorsAttn-1 = dorsal attention network; DorsAttn-2 = dorsal attention network; Amy-Hip = amygdala–hippocampus network; Striatum and Thalamus.

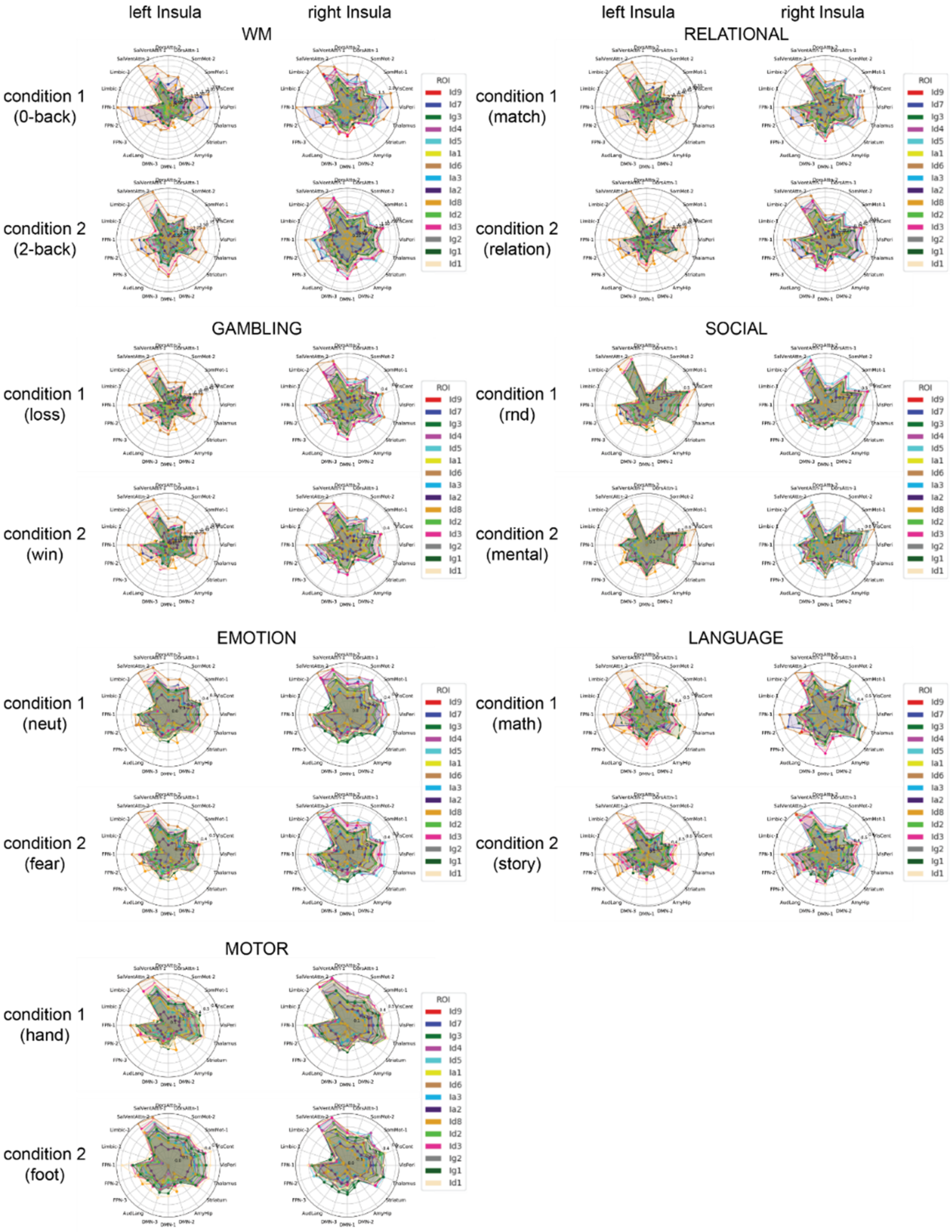

**Figure S9** Illustration of task-dependent connectivity between insular subregions (Julich atlas) and other brain regions (Brainnectome atlas). The insular-brain connectivities were group by brain regions' network membership. The networks were labels as: AudLang = auditory language network; VisCent = visual central network; VisPeri = visual peripheral network; SomMot-1 = somatomotor network; SomMot-2 = somatomotor network; SalVentAttn-1 = salience/ventral attention network; SAIVentAttn-2 = salience/ventral attention network; DMN-1= dorsal default mode network; DMN-2 = ventral default mode network; DMN-3 = default mode network; Limbic-1 = limbic network; Limbic-2 = limbic network; FPN-1 = frontoparietal network; FPN-2 = frontoparietal network; FPN-3 = frontoparietal network; DorsAttn-1 = dorsal attention network; DorsAttn-2 = dorsal attention network; Amy-Hip = amygdala–hippocampus network; Striatum and Thalamus.

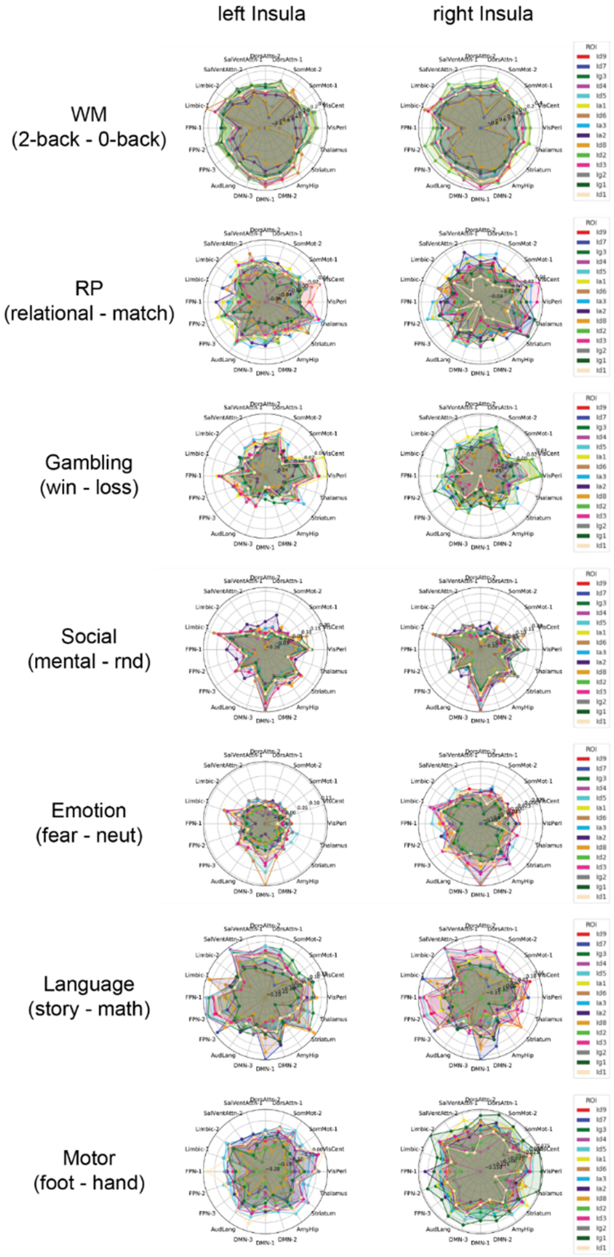

**Figure S10** Illustration of experimental modulation of task-dependent connectivity between insular subregions (Julich atlas) and other brain regions (Brainnectome atlas). The insular-brain connectivities were group by brain regions' network membership. The networks were labels as: AudLang = auditory language network; VisCent = visual central network; VisPeri = visual peripheral network; SomMot-1 = somatomotor network; SomMot-2 = somatomotor network; SalVentAttn-1 = salience/ventral attention network; SAlVentAttn-2 = salience/ventral attention network; DMN-1= dorsal default mode network; DMN-2 = ventral default mode network; DMN-3 = default mode network; Limbic-1 = limbic network; Limbic-2 = limbic network; FPN-1 = frontoparietal network; FPN-2 = frontoparietal network; FPN-3 = frontoparietal network; DorsAttn-1 = dorsal attention network; DorsAttn-2 = dorsal attention network; Amy-Hip = amygdala–hippocampus network; Striatum and Thalamus.

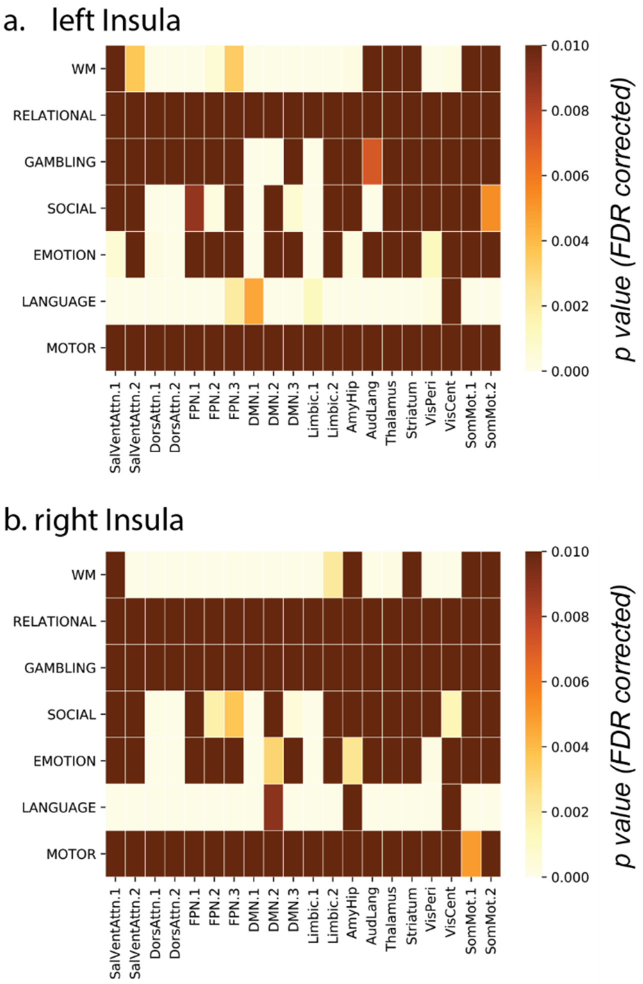

**Figure S11** The statistical significance of interaction effect between the SUBREGION and CONDITION factors in the gPPI weights (Deen atlas) was thresholded at $p<0.05$, corrected. Multiple comparisons (7 tasks x 20 networks x 2 left/right insula) were corrected using FDR method.

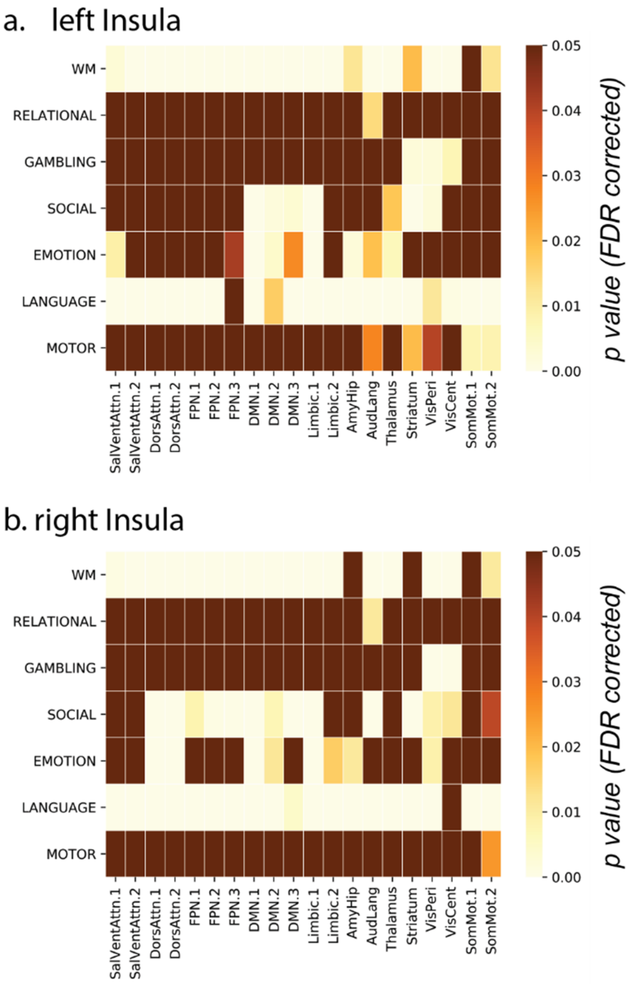

**Figure S12** The statistical significance of interaction effect between the SUBREGION and CONDITION factors in the gPPI weights (Ryali atlas) was thresholded at $p<0.05$, corrected. Multiple comparisons (7 tasks x 20 networks x 2 left/right insula) were corrected using FDR method.

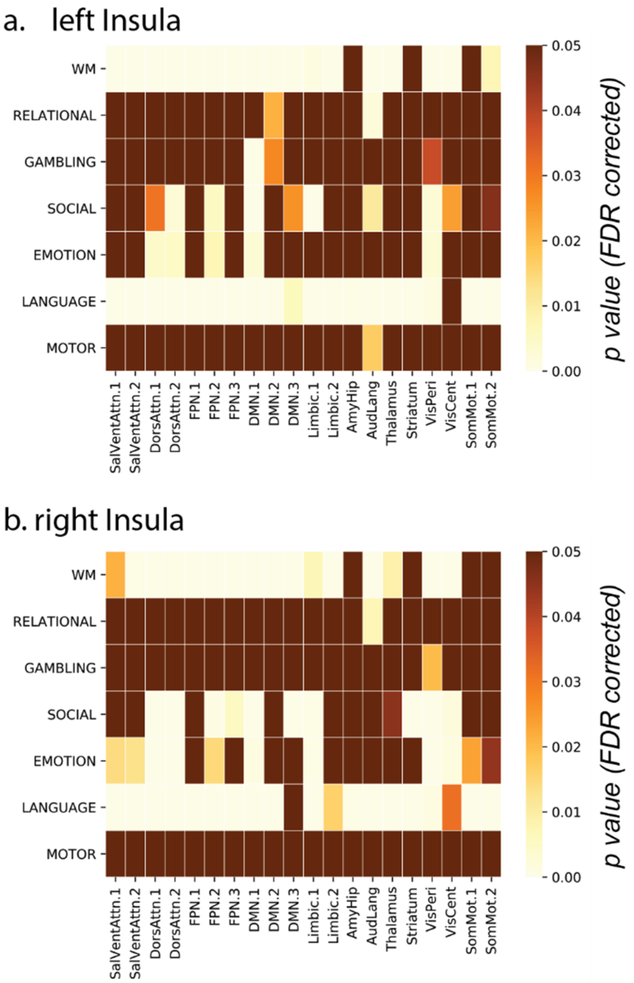

**Figure S13** The statistical significance of interaction effect between the SUBREGION and CONDITION factors in the gPPI weights (Faillenot atlas) was thresholded at $p<0.05$, corrected. Multiple comparisons (7 tasks x 20 networks x 2 left/right insula) were corrected using FDR method.

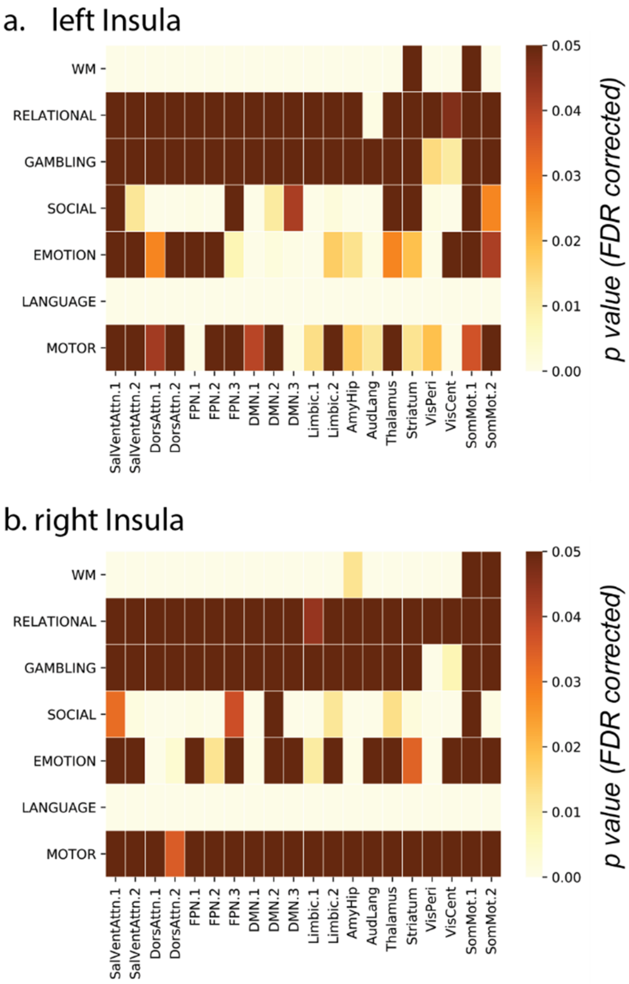

**Figure S14** The statistical significance of interaction effect between the SUBREGION and CONDITION factors in the gPPI weights (Julich atlas) was thresholded at $p<0.05$, corrected. Multiple comparisons (7 tasks x 20 networks x 2 left/right insula) were corrected using FDR method.

**Supplementary Table S1** Percentage of overlapping voxels between ROIs and Atlases

| | | left Insula | | | | | | | | | | | |
|---|---|---|---|---|---|---|---|---|---|---|---|---|---|
| | | Deen | | | Ryali | | | Faillenot | | | | | |
| | | vAI | dAI | PI | vAI | dAI | PI | AIC | ALG | ASG | MSG | PLG | PSG |
| Ryali | vAI | **28** | 13 | 0 | | | | | | | | | |
| | dAI | 3 | **25** | 1 | 0 | | | | | | | | |
| | PI | 0 | 7 | **24** | 0 | 0 | | | | | | | |
| Faillenot | AIC | **31** | 10 | 2 | **18** | 10 | 1 | | | | | | |
| | ALG | 0 | 4 | 18 | 0 | 0 | **25** | 0 | | | | | |
| | ASG | 3 | **17** | 0 | 17 | **12** | 0 | 0 | 0 | | | | |
| | MSG | 0 | 8 | 3 | 0 | 10 | 2 | 0 | 0 | 0 | | | |
| | PLG | 0 | 4 | 0 | 0 | 0 | 10 | 0 | 0 | 0 | 0 | | |
| | PSG | 0 | 2 | **24** | 0 | 2 | 17 | 0 | 0 | 0 | 0 | 0 | |
| Julich | Ig3 | 0 | 0 | 9 | 0 | 0 | 5 | 0 | 6 | 0 | 0 | 0 | 0 |
| | Id4 | 0 | 0 | 10 | 0 | 0 | 11 | 0 | 1 | 0 | 0 | 0 | **24** |
| | Id5 | 0 | 3 | **30** | 0 | 0 | **13** | 0 | **15** | 0 | 0 | 0 | 12 |
| | Ia1 | 0 | 4 | 0 | 0 | 0 | 3 | 0 | 2 | 0 | 0 | 3 | 0 |
| | Id6 | 0 | **28** | 7 | 7 | **31** | 6 | 3 | 0 | 11 | **24** | 0 | 9 |
| | Id9 | 2 | 11 | 3 | 0 | 12 | 1 | **16** | 0 | 0 | 0 | 0 | 1 |
| | Ia3 | 0 | 0 | 1 | 0 | 2 | 1 | 1 | 1 | 0 | 0 | 3 | 0 |
| | Ia2 | 0 | 0 | 0 | 0 | 0 | 0 | 0 | 0 | 0 | 0 | 0 | 0 |
| | Id8 | **11** | 9 | 0 | **24** | 1 | 0 | 7 | 0 | **15** | 0 | 0 | 0 |
| | Id2 | 0 | 5 | 9 | 0 | 0 | 8 | 0 | 10 | 0 | 0 | 1 | 0 |
| | Id3 | 0 | 4 | 1 | 0 | 0 | 6 | 0 | 1 | 0 | 0 | **18** | 0 |
| | Ig2 | 0 | 1 | 1 | 0 | 0 | 6 | 0 | 4 | 0 | 0 | 8 | 0 |
| | Ig1 | 0 | 0 | 0 | 0 | 0 | 1 | 0 | 3 | 0 | 0 | 4 | 0 |
| | Id1 | 0 | 0 | 0 | 0 | 0 | 0 | 0 | 0 | 0 | 0 | 5 | 0 |
| | Id7 | 0 | 8 | 0 | 2 | 17 | 0 | 0 | 0 | 11 | 0 | 0 | 0 |

| | | right Insula | | | | | | | | | | | |
|---|---|---|---|---|---|---|---|---|---|---|---|---|---|
| | | Deen | | | Ryali | | | Faillenot | | | | | |
| | | vAI | dAI | PI | vAI | dAI | PI | AIC | ALG | ASG | MSG | PLG | PSG |
| Ryali | vAI | **29** | 0 | 0 | | | | | | | | | |
| | dAI | 6 | **33** | 0 | 0 | | | | | | | | |
| | PI | 3 | 5 | **23** | 0 | 0 | | | | | | | |
| Faillenot | AIC | **30** | 8 | 0 | **18** | 9 | 0 | | | | | | |
| | ALG | 1 | 1 | **24** | 0 | 0 | **23** | 0 | | | | | |
| | ASG | 5 | **13** | 0 | 1 | **12** | 0 | 0 | 0 | | | | |
| | MSG | 0 | 11 | 0 | 0 | 7 | 0 | 0 | 0 | 0 | | | |
| | PLG | 3 | 0 | 4 | 0 | 0 | 13 | 0 | 0 | 0 | 0 | | |
| | PSG | 0 | 13 | 6 | 0 | 4 | 11 | 0 | 0 | 0 | 0 | 0 | |
| Julich | Ig3 | 0 | 0 | **15** | 0 | 0 | 7 | 0 | 11 | 0 | 0 | 0 | 0 |
| | Id4 | 0 | 4 | 7 | 0 | 0 | 12 | 0 | 2 | 0 | 0 | 0 | **34** |
| | Id5 | 0 | 9 | 14 | 0 | 2 | 8 | 1 | **16** | 0 | 0 | 0 | 19 |
| | Ia1 | 3 | 1 | 3 | 0 | 1 | 3 | 0 | 4 | 0 | 0 | 2 | 0 |
| | Id6 | 0 | **32** | 0 | 0 | **22** | 0 | 1 | 0 | 13 | **30** | 0 | 10 |
| | Id9 | **14** | 12 | 0 | **4** | 14 | 0 | **25** | 0 | 1 | 0 | 0 | 0 |
| | Ia3 | 5 | 0 | 0 | 0 | 0 | 1 | 1 | 1 | 0 | 0 | 2 | 0 |
| | Ia2 | 0 | 0 | 0 | 0 | 0 | 0 | 0 | 0 | 0 | 0 | 1 | 0 |
| | Id8 | 9 | 5 | 0 | 4 | 10 | 0 | 3 | 0 | **18** | 0 | 0 | 0 |
| | Id2 | 2 | 0 | 15 | 0 | 0 | 11 | 0 | 9 | 0 | 0 | 2 | 0 |
| | Id3 | 2 | 0 | 4 | 0 | 0 | 9 | 0 | 1 | 0 | 0 | **26** | 0 |
| | Ig2 | 0 | 0 | 12 | 0 | 0 | **13** | 0 | 9 | 0 | 0 | 9 | 0 |
| | Ig1 | 0 | 0 | 0 | 0 | 0 | 3 | 0 | 3 | 0 | 0 | 6 | 0 |
| | Id1 | 0 | 0 | 0 | 0 | 0 | 2 | 0 | 0 | 0 | 0 | 7 | 0 |
| | Id7 | 0 | 3 | 0 | 0 | 7 | 0 | 0 | 0 | 6 | 0 | 0 | 0 |

**Supplementary Table S2** Cross validation accuracies in task-dependent and ROI-specific voxel-wise activation patterns to differentiate task conditions (Deen atlas).

| left insula | | | |
|---|---|---|---|
| | vAI | dAI | PI |
| WM | 0.70 | 0.75 | 0.66 |
| RELATIONAL | 0.60 | 0.64 | 0.59 |
| GAMBLING | 0.54 | 0.55 | 0.52 |
| SOCIAL | 0.69 | 0.69 | 0.65 |
| EMOTION | 0.58 | 0.57 | 0.54 |
| LANGUAGE | 0.76 | 0.81 | 0.70 |
| MOTOR | 0.55 | 0.65 | 0.77 |
| right insula | | | |
| | vAI | dAI | PI |
| WM | 0.74 | 0.75 | 0.66 |
| RELATIONAL | 0.64 | 0.65 | 0.56 |
| GAMBLING | 0.58 | 0.56 | 0.53 |
| SOCIAL | 0.73 | 0.72 | 0.64 |
| EMOTION | 0.60 | 0.56 | 0.56 |
| LANGUAGE | 0.82 | 0.78 | 0.66 |
| MOTOR | 0.60 | 0.67 | 0.76 |

**Supplementary Table S3** Cross validation accuracies in task-dependent and ROI-specific voxel-wise activation patterns to differentiate task conditions (Ryali atlas).

| left insula | | | |
|---|---|---|---|
| | vAI | dAI | PI |
| WM | 0.74 | 0.71 | 0.65 |
| RELATIONAL | 0.60 | 0.61 | 0.58 |
| GAMBLING | 0.56 | 0.56 | 0.51 |
| SOCIAL | 0.71 | 0.71 | 0.67 |
| EMOTION | 0.60 | 0.59 | 0.55 |
| LANGUAGE | 0.80 | 0.80 | 0.70 |
| MOTOR | 0.57 | 0.64 | 0.81 |
| right insula | | | |
| | vAI | dAI | PI |
| WM | 0.72 | 0.72 | 0.64 |
| RELATIONAL | 0.60 | 0.61 | 0.57 |
| GAMBLING | 0.56 | 0.55 | 0.54 |
| SOCIAL | 0.68 | 0.74 | 0.69 |
| EMOTION | 0.59 | 0.59 | 0.57 |
| LANGUAGE | 0.80 | 0.81 | 0.72 |
| MOTOR | 0.54 | 0.65 | 0.83 |

**Supplementary Table S4** Cross validation accuracies in task-dependent and ROI-specific voxel-wise activation patterns to differentiate task conditions (Faillenot atlas).

| left insula | | | | | | |
|---|---|---|---|---|---|---|
| | AIC | ALG | ASG | MSG | PLG | PSG |
| WM | 0.72 | 0.65 | 0.71 | 0.69 | 0.61 | 0.64 |
| RELATIONAL | 0.62 | 0.56 | 0.59 | 0.59 | 0.59 | 0.59 |
| GAMBLING | 0.56 | 0.53 | 0.60 | 0.51 | 0.58 | 0.49 |
| SOCIAL | 0.71 | 0.68 | 0.66 | 0.59 | 0.70 | 0.63 |
| EMOTION | 0.60 | 0.58 | 0.57 | 0.52 | 0.59 | 0.52 |
| LANGUAGE | 0.80 | 0.67 | 0.75 | 0.75 | 0.73 | 0.65 |
| MOTOR | 0.63 | 0.85 | 0.55 | 0.68 | 0.77 | 0.80 |
| right insula | | | | | | |
| | AIC | ALG | ASG | MSG | PLG | PSG |
| WM | 0.75 | 0.67 | 0.74 | 0.71 | 0.64 | 0.64 |
| RELATIONAL | 0.61 | 0.55 | 0.63 | 0.59 | 0.56 | 0.59 |
| GAMBLING | 0.58 | 0.54 | 0.62 | 0.52 | 0.56 | 0.53 |
| SOCIAL | 0.70 | 0.68 | 0.67 | 0.62 | 0.72 | 0.65 |
| EMOTION | 0.58 | 0.59 | 0.56 | 0.54 | 0.61 | 0.52 |
| LANGUAGE | 0.79 | 0.67 | 0.77 | 0.72 | 0.69 | 0.68 |
| MOTOR | 0.63 | 0.82 | 0.57 | 0.70 | 0.75 | 0.81 |

**Supplementary Table S5** Cross validation accuracies in task-dependent and ROI-specific voxel-wise activation patterns to differentiate task conditions (Julich atlas).

| | left insula | | | | | | | | | | | | | | |
|---|---|---|---|---|---|---|---|---|---|---|---|---|---|---|---|
| | Ig3 | Id4 | Id5 | Ia1 | Id6 | Id9 | Ia3 | Ia2 | Id8 | Id2 | Id3 | Ig2 | Ig1 | Id1 | Id7 |
| WM | 0.64 | 0.63 | 0.64 | 0.64 | 0.74 | 0.67 | 0.64 | 0.63 | 0.72 | 0.65 | 0.65 | 0.66 | 0.65 | 0.63 | 0.69 |
| RELATIONAL | 0.58 | 0.59 | 0.59 | 0.59 | 0.63 | 0.59 | 0.55 | 0.57 | 0.61 | 0.60 | 0.58 | 0.58 | 0.54 | 0.57 | 0.58 |
| GAMBLING | 0.52 | 0.51 | 0.50 | 0.54 | 0.53 | 0.56 | 0.54 | 0.53 | 0.54 | 0.50 | 0.55 | 0.53 | 0.53 | 0.52 | 0.50 |
| SOCIAL | 0.56 | 0.62 | 0.64 | 0.63 | 0.68 | 0.67 | 0.70 | 0.69 | 0.67 | 0.58 | 0.75 | 0.58 | 0.56 | 0.70 | 0.65 |
| EMOTION | 0.50 | 0.54 | 0.53 | 0.52 | 0.57 | 0.58 | 0.59 | 0.58 | 0.57 | 0.52 | 0.58 | 0.54 | 0.53 | 0.55 | 0.55 |
| LANGUAGE | 0.62 | 0.61 | 0.67 | 0.59 | 0.77 | 0.77 | 0.72 | 0.79 | 0.79 | 0.65 | 0.80 | 0.63 | 0.63 | 0.77 | 0.74 |
| MOTOR | 0.71 | 0.74 | 0.74 | 0.58 | 0.69 | 0.60 | 0.55 | 0.53 | 0.54 | 0.72 | 0.70 | 0.77 | 0.74 | 0.63 | 0.53 |
| | right insula | | | | | | | | | | | | | | |
| | Ig3 | Id4 | Id5 | Ia1 | Id6 | Id9 | Ia3 | Ia2 | Id8 | Id2 | Id3 | Ig2 | Ig1 | Id1 | Id7 |
| WM | 0.66 | 0.60 | 0.64 | 0.65 | 0.73 | 0.72 | 0.64 | 0.62 | 0.74 | 0.66 | 0.64 | 0.66 | 0.61 | 0.59 | 0.72 |
| RELATIONAL | 0.54 | 0.55 | 0.58 | 0.57 | 0.63 | 0.63 | 0.58 | 0.56 | 0.63 | 0.56 | 0.55 | 0.55 | 0.55 | 0.56 | 0.61 |
| GAMBLING | 0.51 | 0.54 | 0.51 | 0.55 | 0.53 | 0.56 | 0.55 | 0.52 | 0.56 | 0.54 | 0.56 | 0.53 | 0.53 | 0.53 | 0.54 |
| SOCIAL | 0.56 | 0.64 | 0.67 | 0.67 | 0.65 | 0.70 | 0.71 | 0.69 | 0.69 | 0.64 | 0.78 | 0.62 | 0.60 | 0.74 | 0.68 |
| EMOTION | 0.52 | 0.53 | 0.54 | 0.57 | 0.56 | 0.57 | 0.60 | 0.58 | 0.61 | 0.53 | 0.60 | 0.54 | 0.53 | 0.59 | 0.55 |
| LANGUAGE | 0.62 | 0.69 | 0.70 | 0.63 | 0.78 | 0.78 | 0.69 | 0.69 | 0.78 | 0.64 | 0.73 | 0.63 | 0.62 | 0.69 | 0.77 |
| MOTOR | 0.75 | 0.76 | 0.79 | 0.60 | 0.70 | 0.59 | 0.57 | 0.53 | 0.54 | 0.71 | 0.70 | 0.80 | 0.72 | 0.64 | 0.54 |

**Supplementary Table S6** Repeated ANOVA with factors of SUBREGION and CONDITION reveals significant main effects and interaction effect on task-dependent condition-specific activation in the HCP tasks (Deen atlas). Table shows p values of main effects and interaction effects.

| left insula | | | |
|---|---|---|---|
| Task | SUBREGION | CONDITION | SUBREGIONxCONDITION |
| WM | 8.45E-76 | 2.00E-06 | 7.13E-45 |
| RELATIONAL | 2.57E-137 | 5.39E-02 | 3.94E-29 |
| GAMBLING | 9.08E-72 | 8.67E-01 | 2.82E-02 |
| SOCIAL | 2.26E-10 | 3.34E-02 | 4.17E-05 |
| EMOTION | 2.84E-07 | 8.16E-01 | 3.00E-01 |
| LANGUAGE | 1.12E-55 | 3.57E-21 | 2.77E-185 |
| MOTOR | 5.29E-26 | 7.83E-05 | 2.40E-11 |
| right insula | | | |
| Task | SUBREGION | CONDITION | SUBREGIONxCONDITION |
| WM | 5.16E-223 | 2.28E-02 | 3.57E-95 |
| RELATIONAL | 5.42E-242 | 6.06E-01 | 4.78E-31 |
| GAMBLING | 4.18E-259 | 3.99E-01 | 2.39E-03 |
| SOCIAL | 1.07E-63 | 3.20E-04 | 1.71E-06 |
| EMOTION | 4.12E-26 | 7.00E-02 | 1.23E-01 |
| LANGUAGE | 1.48E-50 | 1.61E-51 | 1.02E-297 |
| MOTOR | 1.28E-58 | 4.79E-07 | 2.41E-21 |

**Supplementary Table S7** Repeated ANOVA with factors of SUBREGION and CONDITION reveals significant main effects and interaction effect on task-dependent condition-specific activation in the HCP tasks (Ryali atlas). Table shows p values of main effects and interaction effects.

| left insula | | | |
|---|---|---|---|
| Task | SUBREGION | CONDITION | SUBREGIONxCONDITION |
| WM | 4.01E-160 | 7.38E-01 | 2.72E-115 |
| RELATIONAL | 5.84E-145 | 5.56E-01 | 3.51E-77 |
| GAMBLING | 3.04E-208 | 3.11E-01 | 2.45E-01 |
| SOCIAL | 1.85E-52 | 5.71E-02 | 2.87E-22 |
| EMOTION | 8.22E-01 | 5.84E-01 | 2.35E-06 |
| LANGUAGE | 1.65E-13 | 2.53E-56 | 1.81E-268 |
| MOTOR | 2.82E-21 | 3.18E-10 | 1.79E-05 |
| right insula | | | |
| Task | SUBREGION | CONDITION | SUBREGIONxCONDITION |
| WM | 2.30E-213 | 2.46E-01 | 9.74E-92 |
| RELATIONAL | 1.95E-222 | 5.01E-02 | 2.76E-36 |
| GAMBLING | 1.50E-238 | 3.21E-01 | 6.37E-03 |
| SOCIAL | 2.15E-72 | 8.60E-01 | 1.14E-29 |
| EMOTION | 3.36E-10 | 5.50E-03 | 7.74E-03 |
| LANGUAGE | 5.02E-13 | 6.95E-32 | 3.17E-247 |
| MOTOR | 4.40E-16 | 1.71E-09 | 8.71E-18 |

**Supplementary Table S8** Repeated ANOVA with factors of SUBREGION and CONDITION reveals significant main effects and interaction effect on task-dependent condition-specific activation in the HCP tasks (Faillenot atlas). Table shows p values of main effects and interaction effects.

| left insula | | | |
|---|---|---|---|
| Task | SUBREGION | CONDITION | SUBREGIONxCONDITION |
| WM | 0.00E+00 | 8.22E-06 | 3.71E-241 |
| RELATIONAL | 0.00E+00 | 3.26E-04 | 2.47E-119 |
| GAMBLING | 0.00E+00 | 4.54E-01 | 7.22E-04 |
| SOCIAL | 1.13E-157 | 4.11E-01 | 3.55E-25 |
| EMOTION | 1.61E-25 | 2.26E-01 | 2.43E-17 |
| LANGUAGE | 8.62E-170 | 1.67E-09 | 0.00E+00 |
| MOTOR | 4.47E-44 | 3.35E-10 | 1.97E-33 |
| right insula | | | |
| Task | SUBREGION | CONDITION | SUBREGIONxCONDITION |
| WM | 0.00E+00 | 4.95E-01 | 5.11E-311 |
| RELATIONAL | 0.00E+00 | 7.70E-01 | 1.59E-160 |
| GAMBLING | 0.00E+00 | 4.36E-01 | 1.45E-04 |
| SOCIAL | 1.47E-246 | 9.81E-01 | 2.58E-20 |
| EMOTION | 6.83E-23 | 1.51E-04 | 2.20E-13 |
| LANGUAGE | 1.99E-150 | 8.87E-40 | 0.00E+00 |
| MOTOR | 5.67E-34 | 1.71E-12 | 1.99E-37 |

**Supplementary Table S9** Repeated ANOVA with factors of SUBREGION and CONDITION reveals significant main effects and interaction effect on task-dependent condition-specific activation in the HCP tasks (Julich atlas). Table shows p values of main effects and interaction effects.

| left insula | | | |
|---|---|---|---|
| Task | SUBREGION | CONDITION | SUBREGIONxCONDITION |
| WM | 0.00E+00 | 2.94E-21 | 0.00E+00 |
| RELATIONAL | 0.00E+00 | 4.64E-08 | 3.56E-295 |
| GAMBLING | 0.00E+00 | 6.96E-01 | 1.14E-11 |
| SOCIAL | 0.00E+00 | 7.76E-01 | 3.73E-148 |
| EMOTION | 1.48E-52 | 2.30E-02 | 3.69E-58 |
| LANGUAGE | 0.00E+00 | 2.92E-21 | 0.00E+00 |
| MOTOR | 1.25E-219 | 4.30E-07 | 1.93E-121 |
| right insula | | | |
| Task | SUBREGION | CONDITION | SUBREGIONxCONDITION |
| WM | 0.00E+00 | 8.10E-11 | 0.00E+00 |
| RELATIONAL | 0.00E+00 | 9.94E-03 | 0.00E+00 |
| GAMBLING | 0.00E+00 | 2.15E-01 | 8.41E-12 |
| SOCIAL | 0.00E+00 | 6.03E-01 | 1.29E-209 |
| EMOTION | 3.73E-90 | 2.62E-06 | 2.29E-85 |
| LANGUAGE | 0.00E+00 | 1.61E-01 | 0.00E+00 |
| MOTOR | 6.62E-183 | 1.50E-09 | 3.62E-54 |

**Supplementary Table S10** Repeated ANOVA with factors of SUBREGION and CONDITION reveals significant main effects and interaction effect on task-dependent condition-specific connectivity in the HCP tasks (Deen atlas). Table shows p values of main effects and interaction effects for insula's connectivity to brain networks.

left insular

main effect of SUBREGION

| | SalVentAttn | SalVentAttn | DorsAttn.1 | DorsAttn.2 | FPN.1 | FPN.2 | FPN.3 | DMN.1 | DMN.2 | DMN.3 | Limbic.1 | Limbic.2 | AmyHip | AudLang | Thalamus | Striatum | VisPeri | VisCent | SomMot.1 | SomMot.2 |
|---|---|---|---|---|---|---|---|---|---|---|---|---|---|---|---|---|---|---|---|---|
| WM | 9.67E-112 | 2.76E-74 | 1.34E-68 | 1.58E-97 | 5.04E-53 | 8.71E-45 | 2.85E-31 | 8.19E-05 | 2.06E-25 | 3.81E-22 | 3.12E-08 | 1.27E-15 | 3.70E-32 | 2.78E-15 | 4.11E-42 | 3.01E-54 | 5.28E-53 | 1.95E-46 | 4.44E-109 | 5.84E-129 |
| RELATIONAL | 3.12E-53 | 4.45E-31 | 7.66E-30 | 2.51E-46 | 1.20E-25 | 2.12E-17 | 2.18E-11 | 1.09E-07 | 3.61E-14 | 5.41E-07 | 1.50E-06 | 2.15E-02 | 5.39E-11 | 1.39E-05 | 7.94E-31 | 8.84E-21 | 1.85E-23 | 1.32E-33 | 8.80E-49 | 7.28E-71 |
| GAMBLING | 8.76E-73 | 6.41E-45 | 1.99E-56 | 1.52E-68 | 1.48E-31 | 8.30E-21 | 3.05E-08 | 1.78E-11 | 8.40E-14 | 1.21E-09 | 6.52E-09 | 3.67E-02 | 1.49E-07 | 1.03E-07 | 2.90E-33 | 1.35E-28 | 1.08E-45 | 3.61E-40 | 2.40E-62 | 8.77E-76 |
| SOCIAL | 3.90E-46 | 1.77E-10 | 1.36E-03 | 8.59E-09 | 3.35E-01 | 5.03E-03 | 3.30E-05 | 8.09E-35 | 2.37E-26 | 3.75E-01 | 4.86E-35 | 1.18E-07 | 4.51E-11 | 5.21E-01 | 2.02E-09 | 2.29E-08 | 5.07E-24 | 1.10E-28 | 1.85E-35 | 4.67E-55 |
| EMOTION | 8.07E-104 | 4.23E-57 | 5.39E-95 | 6.30E-117 | 1.88E-09 | 2.46E-47 | 3.04E-05 | 3.19E-100 | 4.16E-51 | 1.92E-19 | 4.24E-68 | 1.86E-11 | 5.00E-24 | 3.02E-05 | 2.66E-14 | 1.33E-12 | 1.25E-55 | 2.02E-23 | 8.53E-68 | 1.98E-96 |
| LANGUAGE | 1.67E-99 | 8.57E-57 | 4.45E-77 | 9.11E-93 | 1.23E-48 | 2.01E-52 | 3.76E-39 | 3.69E-37 | 1.61E-59 | 6.08E-24 | 5.37E-34 | 1.02E-44 | 1.87E-64 | 2.09E-02 | 2.37E-33 | 2.80E-34 | 1.93E-22 | 1.63E-29 | 1.47E-131 | 8.96E-122 |
| MOTOR | 4.79E-51 | 1.85E-26 | 2.18E-34 | 4.63E-61 | 2.54E-03 | 3.06E-16 | 4.55E-05 | 6.82E-56 | 4.22E-29 | 5.83E-13 | 3.49E-39 | 3.75E-08 | 2.53E-18 | 6.93E-03 | 1.71E-11 | 1.44E-11 | 4.31E-14 | 1.93E-01 | 1.76E-34 | 3.58E-41 |

main effect of CONDITION

| | SalVentAttn | SalVentAttn | DorsAttn.1 | DorsAttn.2 | FPN.1 | FPN.2 | FPN.3 | DMN.1 | DMN.2 | DMN.3 | Limbic.1 | Limbic.2 | AmyHip | AudLang | Thalamus | Striatum | VisPeri | VisCent | SomMot.1 | SomMot.2 |
|---|---|---|---|---|---|---|---|---|---|---|---|---|---|---|---|---|---|---|---|---|
| WM | 3.91E-01 | 3.15E-04 | 5.00E-04 | 8.62E-09 | 3.14E-15 | 3.20E-04 | 8.08E-04 | 3.00E-05 | 1.99E-05 | 3.76E-03 | 6.75E-10 | 8.90E-07 | 1.96E-09 | 5.51E-02 | 1.72E-01 | 4.43E-01 | 3.59E-07 | 4.34E-02 | 1.24E-01 | 1.61E-03 |
| RELATIONAL | 3.99E-01 | 7.79E-01 | 6.15E-01 | 9.30E-01 | 4.27E-01 | 9.12E-01 | 2.08E-01 | 2.77E-01 | 7.74E-01 | 7.17E-01 | 1.21E-01 | 6.46E-01 | 5.65E-01 | 3.16E-01 | 3.70E-01 | 4.74E-01 | 7.20E-01 | 7.47E-01 | 7.39E-01 | 9.37E-01 |
| GAMBLING | 9.81E-02 | 5.94E-01 | 3.25E-01 | 7.93E-01 | 7.02E-01 | 5.40E-01 | 7.15E-01 | 9.33E-02 | 1.29E-01 | 4.68E-02 | 1.20E-01 | 1.39E-01 | 6.62E-02 | 3.21E-01 | 4.51E-01 | 4.45E-01 | 8.43E-01 | 9.87E-01 | 1.87E-01 | 1.88E-01 |
| SOCIAL | 2.77E-01 | 1.18E-01 | 3.98E-02 | 1.27E-04 | 8.70E-02 | 4.94E-01 | 2.49E-09 | 5.12E-19 | 8.41E-09 | 2.63E-03 | 3.90E-20 | 2.93E-06 | 2.64E-18 | 6.46E-01 | 3.81E-02 | 1.26E-17 | 3.77E-11 | 3.55E-06 | 2.00E-03 | 9.74E-03 |
| EMOTION | 7.67E-01 | 1.31E-01 | 6.24E-01 | 9.99E-01 | 2.22E-02 | 3.25E-01 | 2.45E-03 | 1.38E-03 | 9.59E-02 | 2.27E-02 | 8.52E-03 | 7.81E-01 | 4.55E-01 | 8.09E-02 | 1.24E-01 | 1.19E-01 | 8.29E-01 | 1.52E-01 | 8.85E-01 | 6.59E-01 |
| LANGUAGE | 1.09E-10 | 2.30E-41 | 1.64E-09 | 5.63E-17 | 1.08E-24 | 2.08E-32 | 1.30E-29 | 3.54E-04 | 9.16E-01 | 4.39E-01 | 7.09E-09 | 2.23E-04 | 1.43E-06 | 5.86E-16 | 4.00E-06 | 2.49E-27 | 2.31E-11 | 2.71E-14 | 5.14E-01 | 1.85E-04 |
| MOTOR | 2.05E-05 | 4.30E-04 | 3.98E-04 | 3.94E-06 | 3.15E-05 | 3.32E-04 | 4.73E-04 | 1.09E-03 | 1.05E-03 | 3.69E-04 | 2.83E-04 | 1.63E-02 | 1.19E-03 | 5.82E-04 | 1.00E-04 | 1.53E-03 | 3.71E-04 | 2.31E-03 | 4.05E-06 | 1.01E-02 |

interaction effect of SUBREGIONxCONDITION

| | SalVentAttn | SalVentAttn | DorsAttn.1 | DorsAttn.2 | FPN.1 | FPN.2 | FPN.3 | DMN.1 | DMN.2 | DMN.3 | Limbic.1 | Limbic.2 | AmyHip | AudLang | Thalamus | Striatum | VisPeri | VisCent | SomMot.1 | SomMot.2 |
|---|---|---|---|---|---|---|---|---|---|---|---|---|---|---|---|---|---|---|---|---|
| WM | 4.63E-02 | 1.22E-03 | 2.26E-06 | 2.09E-08 | 9.83E-08 | 1.87E-04 | 1.15E-03 | 1.10E-23 | 9.06E-21 | 3.75E-06 | 5.07E-18 | 2.13E-08 | 5.03E-05 | 9.71E-03 | 9.09E-01 | 5.11E-01 | 5.77E-07 | 5.76E-05 | 5.63E-01 | 6.46E-02 |
| RELATIONAL | 6.37E-01 | 6.30E-01 | 5.01E-03 | 4.47E-02 | 5.06E-01 | 2.40E-01 | 9.19E-01 | 2.81E-01 | 8.53E-01 | 6.08E-01 | 9.76E-01 | 2.61E-01 | 4.87E-01 | 3.52E-01 | 5.25E-01 | 9.37E-01 | 1.52E-01 | 7.12E-01 | 7.08E-01 | 2.10E-01 |
| GAMBLING | 2.30E-01 | 8.42E-02 | 9.65E-01 | 8.13E-01 | 1.13E-01 | 2.34E-01 | 6.55E-03 | 2.31E-09 | 1.53E-05 | 2.42E-02 | 8.46E-06 | 1.36E-02 | 1.65E-01 | 2.54E-03 | 9.70E-01 | 9.06E-01 | 6.97E-01 | 8.76E-01 | 8.64E-01 | 2.07E-01 |
| SOCIAL | 3.38E-02 | 5.41E-03 | 1.89E-07 | 2.02E-07 | 3.19E-03 | 7.67E-05 | 5.94E-01 | 2.15E-15 | 2.44E-02 | 2.02E-04 | 3.64E-10 | 5.38E-01 | 7.47E-02 | 4.34E-06 | 8.51E-01 | 1.78E-02 | 6.97E-01 | 2.82E-01 | 5.64E-02 | 1.88E-03 |
| EMOTION | 1.85E-04 | 5.81E-02 | 5.01E-05 | 2.35E-06 | 1.53E-01 | 5.41E-02 | 3.00E-01 | 6.27E-09 | 9.02E-03 | 2.16E-02 | 5.34E-08 | 4.02E-02 | 5.96E-05 | 2.95E-02 | 1.28E-02 | 7.01E-02 | 3.79E-04 | 4.41E-01 | 2.39E-02 | 1.07E-02 |
| LANGUAGE | 3.30E-20 | 2.42E-17 | 8.65E-37 | 1.91E-59 | 6.51E-18 | 9.91E-47 | 6.46E-04 | 1.59E-03 | 3.04E-10 | 1.55E-08 | 3.63E-04 | 6.34E-21 | 1.48E-08 | 6.66E-60 | 5.16E-41 | 5.48E-32 | 6.32E-05 | 1.94E-02 | 6.63E-21 | 2.44E-58 |
| MOTOR | 8.65E-01 | 5.27E-01 | 5.33E-01 | 3.09E-01 | 5.26E-03 | 7.82E-01 | 8.38E-02 | 3.21E-02 | 4.66E-01 | 3.81E-02 | 1.05E-01 | 3.79E-01 | 2.55E-01 | 4.84E-01 | 6.92E-01 | 7.05E-01 | 5.89E-01 | 1.30E-01 | 3.20E-01 | 4.98E-01 |

right insular

main effect of SUBREGION

| | SalVentAttn | SalVentAttn | DorsAttn.1 | DorsAttn.2 | FPN.1 | FPN.2 | FPN.3 | DMN.1 | DMN.2 | DMN.3 | Limbic.1 | Limbic.2 | AmyHip | AudLang | Thalamus | Striatum | VisPeri | VisCent | SomMot.1 | SomMot.2 |
|---|---|---|---|---|---|---|---|---|---|---|---|---|---|---|---|---|---|---|---|---|
| WM | 4.84E-97 | 4.61E-54 | 2.08E-50 | 6.65E-81 | 2.87E-56 | 4.16E-45 | 4.95E-48 | 3.81E-07 | 7.36E-13 | 5.44E-26 | 1.72E-21 | 9.57E-36 | 8.29E-61 | 5.25E-11 | 1.89E-30 | 9.00E-46 | 1.75E-38 | 2.54E-45 | 7.04E-148 | 3.85E-182 |
| RELATIONAL | 4.32E-33 | 8.45E-16 | 1.13E-21 | 1.40E-35 | 1.97E-14 | 5.69E-11 | 1.93E-07 | 1.45E-07 | 3.48E-06 | 1.09E-12 | 2.45E-09 | 2.94E-11 | 4.59E-27 | 1.10E-03 | 6.05E-15 | 1.49E-18 | 1.29E-18 | 3.33E-23 | 2.85E-56 | 9.42E-69 |
| GAMBLING | 7.06E-53 | 7.99E-23 | 1.50E-34 | 1.40E-48 | 1.66E-21 | 1.39E-15 | 4.15E-08 | 2.71E-11 | 2.11E-08 | 7.35E-13 | 1.44E-14 | 4.32E-17 | 8.15E-26 | 5.23E-08 | 3.93E-17 | 3.47E-12 | 3.34E-28 | 1.07E-29 | 2.83E-79 | 1.75E-98 |
| SOCIAL | 2.20E-52 | 1.75E-05 | 2.18E-08 | 1.83E-17 | 4.78E-01 | 7.58E-04 | 3.93E-04 | 7.56E-23 | 4.15E-10 | 1.17E-14 | 3.39E-19 | 4.31E-24 | 1.73E-38 | 4.30E-05 | 3.38E-08 | 8.65E-10 | 8.52E-17 | 5.76E-24 | 1.88E-78 | 8.89E-114 |
| EMOTION | 6.45E-56 | 2.58E-14 | 6.88E-65 | 5.35E-74 | 2.08E-04 | 2.81E-14 | 9.78E-16 | 1.30E-109 | 1.60E-45 | 4.95E-55 | 1.68E-85 | 5.83E-42 | 8.41E-71 | 5.39E-28 | 1.20E-19 | 3.29E-27 | 3.55E-35 | 5.42E-25 | 1.11E-86 | 2.83E-113 |
| LANGUAGE | 6.32E-66 | 4.84E-30 | 4.81E-62 | 6.90E-59 | 1.91E-25 | 1.04E-36 | 1.83E-49 | 4.04E-48 | 6.41E-31 | 9.14E-59 | 3.12E-41 | 2.08E-59 | 7.07E-114 | 7.80E-32 | 8.41E-23 | 4.51E-27 | 5.77E-31 | 1.00E-35 | 2.18E-186 | 8.25E-188 |
| MOTOR | 6.90E-52 | 7.06E-24 | 2.13E-37 | 3.72E-59 | 1.40E-10 | 1.53E-17 | 3.38E-05 | 1.33E-51 | 8.05E-25 | 2.87E-24 | 1.28E-39 | 3.51E-18 | 8.73E-28 | 1.81E-10 | 2.52E-16 | 1.37E-13 | 1.34E-18 | 5.12E-11 | 3.01E-42 | 2.66E-58 |

main effect of CONDITION

| | SalVentAttn | SalVentAttn | DorsAttn.1 | DorsAttn.2 | FPN.1 | FPN.2 | FPN.3 | DMN.1 | DMN.2 | DMN.3 | Limbic.1 | Limbic.2 | AmyHip | AudLang | Thalamus | Striatum | VisPeri | VisCent | SomMot.1 | SomMot.2 |
|---|---|---|---|---|---|---|---|---|---|---|---|---|---|---|---|---|---|---|---|---|
| WM | 6.34E-01 | 5.38E-02 | 1.35E-07 | 4.45E-09 | 6.30E-12 | 1.84E-03 | 5.63E-01 | 4.73E-07 | 8.34E-10 | 9.39E-01 | 3.32E-08 | 5.46E-04 | 4.38E-04 | 2.62E-02 | 1.17E-01 | 8.08E-03 | 3.04E-15 | 1.45E-06 | 7.97E-01 | 1.20E-01 |
| RELATIONAL | 8.08E-01 | 5.80E-01 | 6.17E-01 | 8.99E-01 | 2.59E-01 | 5.18E-01 | 3.05E-02 | 1.12E-01 | 1.41E-01 | 3.86E-01 | 3.99E-01 | 4.29E-01 | 5.55E-01 | 9.89E-01 | 7.17E-01 | 4.48E-01 | 6.92E-01 | 6.79E-01 | 5.40E-01 | 9.97E-01 |
| GAMBLING | 3.32E-03 | 6.00E-02 | 6.21E-01 | 2.14E-01 | 6.94E-02 | 1.82E-01 | 1.50E-01 | 5.27E-02 | 5.92E-02 | 2.17E-02 | 3.70E-02 | 8.22E-03 | 1.73E-02 | 1.62E-01 | 2.85E-02 | 1.68E-02 | 1.95E-01 | 1.50E-01 | 2.29E-03 | 3.96E-03 |
| SOCIAL | 6.48E-01 | 5.76E-01 | 1.26E-05 | 7.18E-08 | 6.25E-01 | 3.56E-02 | 5.03E-08 | 2.89E-19 | 6.32E-07 | 5.13E-03 | 1.69E-19 | 1.46E-03 | 3.42E-12 | 1.15E-01 | 9.36E-01 | 2.50E-13 | 3.14E-07 | 1.73E-05 | 1.03E-02 | 1.85E-02 |
| EMOTION | 4.01E-01 | 3.54E-01 | 4.73E-01 | 3.85E-01 | 8.67E-02 | 9.82E-01 | 8.19E-02 | 1.45E-01 | 2.92E-01 | 1.07E-01 | 2.22E-01 | 7.51E-01 | 6.63E-01 | 4.59E-01 | 9.28E-01 | 4.86E-01 | 2.25E-01 | 6.37E-01 | 1.01E-01 | 2.52E-01 |
| LANGUAGE | 6.82E-05 | 4.48E-19 | 6.88E-02 | 6.53E-03 | 3.85E-10 | 9.26E-08 | 9.45E-18 | 6.17E-03 | 1.54E-01 | 5.22E-02 | 6.65E-06 | 3.54E-06 | 6.57E-09 | 7.49E-21 | 9.72E-01 | 2.36E-05 | 1.82E-05 | 1.03E-07 | 5.90E-01 | 3.22E-08 |
| MOTOR | 4.86E-03 | 6.40E-03 | 4.78E-02 | 2.02E-03 | 2.59E-03 | 1.50E-02 | 5.14E-03 | 3.76E-02 | 8.29E-03 | 3.52E-02 | 1.95E-03 | 2.91E-01 | 8.21E-02 | 2.85E-02 | 4.91E-03 | 9.22E-04 | 1.56E-02 | 6.04E-02 | 2.48E-03 | 5.60E-02 |

interaction effect of SUBREGIONxCONDITION

| | SalVentAttn | SalVentAttn | DorsAttn.1 | DorsAttn.2 | FPN.1 | FPN.2 | FPN.3 | DMN.1 | DMN.2 | DMN.3 | Limbic.1 | Limbic.2 | AmyHip | AudLang | Thalamus | Striatum | VisPeri | VisCent | SomMot.1 | SomMot.2 |
|---|---|---|---|---|---|---|---|---|---|---|---|---|---|---|---|---|---|---|---|---|
| WM | 2.64E-02 | 1.66E-10 | 6.79E-16 | 3.79E-09 | 5.55E-12 | 3.54E-16 | 4.15E-13 | 6.79E-17 | 1.35E-13 | 1.84E-06 | 3.00E-08 | 7.00E-04 | 1.27E-01 | 1.17E-05 | 4.96E-05 | 4.92E-03 | 1.38E-16 | 7.52E-13 | 8.88E-01 | 4.21E-03 |
| RELATIONAL | 8.67E-01 | 8.71E-01 | 6.47E-02 | 5.55E-01 | 9.83E-01 | 8.42E-01 | 2.64E-01 | 1.10E-01 | 7.63E-01 | 5.56E-01 | 5.59E-02 | 8.13E-01 | 1.15E-01 | 4.69E-01 | 9.59E-01 | 6.48E-01 | 5.42E-01 | 6.69E-01 | 8.26E-01 | 2.66E-01 |
| GAMBLING | 4.88E-01 | 5.29E-01 | 6.87E-01 | 4.64E-01 | 9.85E-01 | 9.96E-01 | 8.07E-01 | 4.55E-01 | 7.20E-01 | 4.55E-01 | 6.17E-01 | 4.10E-01 | 7.86E-01 | 8.68E-01 | 7.26E-01 | 8.99E-01 | 1.22E-02 | 4.50E-02 | 5.77E-01 | 7.13E-01 |
| SOCIAL | 7.18E-01 | 6.12E-01 | 2.44E-05 | 1.90E-06 | 6.66E-01 | 5.78E-04 | 1.26E-03 | 5.88E-15 | 2.05E-02 | 1.20E-04 | 4.67E-06 | 9.34E-01 | 4.10E-02 | 8.99E-02 | 7.50E-02 | 3.88E-03 | 1.04E-01 | 4.52E-04 | 7.39E-01 | 2.08E-01 |
| EMOTION | 3.11E-02 | 1.32E-01 | 4.62E-06 | 3.20E-07 | 5.99E-02 | 2.02E-02 | 6.38E-01 | 2.33E-06 | 1.03E-03 | 7.08E-02 | 1.85E-09 | 1.54E-02 | 7.88E-04 | 6.34E-02 | 1.38E-01 | 7.51E-01 | 3.83E-05 | 9.04E-02 | 1.68E-01 | 2.45E-02 |
| LANGUAGE | 4.11E-11 | 1.53E-10 | 2.92E-36 | 1.65E-53 | 3.35E-11 | 7.08E-28 | 4.36E-08 | 3.46E-16 | 3.29E-03 | 2.85E-06 | 1.29E-18 | 8.53E-08 | 3.81E-03 | 4.34E-49 | 2.36E-26 | 1.71E-33 | 3.53E-06 | 1.63E-02 | 3.01E-21 | 3.74E-60 |
| MOTOR | 4.33E-02 | 4.51E-02 | 9.00E-01 | 3.63E-01 | 6.69E-01 | 8.28E-01 | 4.66E-01 | 1.83E-01 | 7.42E-01 | 4.98E-01 | 4.96E-01 | 9.02E-01 | 2.88E-01 | 6.54E-01 | 7.98E-01 | 8.49E-02 | 6.67E-01 | 7.15E-01 | 1.71E-03 | 6.48E-02 |

**Supplementary Table S11** Repeated ANOVA with factors of SUBREGION and CONDITION reveals significant main effects and interaction effect on task-dependent condition-specific connectivity in the HCP tasks (Ryali atlas). Table shows p values of main effects and interaction effects for insula’s connectivity to brain networks.

| | | | | | | | | | | | | | | | | | | | | |
|---|---|---|---|---|---|---|---|---|---|---|---|---|---|---|---|---|---|---|---|---|
| | | | | | | | | | | | left insular | | | | | | | | | |
| main effect of SUBREGION | | | | | | | | | | | | | | | | | | | | |
| | SalVentAttn.1 | SalVentAttn.2 | DorsAttn.1 | DorsAttn.2 | FPN.1 | FPN.2 | FPN.3 | DMN.1 | DMN.2 | DMN.3 | Limbic.1 | Limbic.2 | AmyHip | AudLang | Thalamus | Striatum | VisPeri | VisCent | SomMot.1 | SomMot.2 |
| WM | 1.89E-125 | 3.90E-32 | 5.47E-34 | 1.01E-58 | 5.88E-38 | 6.73E-44 | 1.13E-48 | 5.24E-04 | 2.03E-39 | 1.85E-27 | 3.67E-11 | 3.81E-25 | 1.16E-65 | 8.36E-04 | 4.54E-21 | 6.71E-44 | 7.15E-30 | 4.83E-38 | 2.43E-154 | 2.72E-193 |
| RELATIONAL | 8.09E-43 | 5.41E-12 | 1.63E-10 | 4.86E-21 | 1.48E-08 | 7.70E-16 | 1.63E-15 | 3.53E-03 | 4.59E-20 | 1.46E-10 | 2.67E-05 | 8.18E-06 | 3.27E-22 | 3.59E-01 | 3.17E-14 | 2.30E-15 | 1.13E-13 | 2.28E-24 | 3.93E-54 | 7.56E-77 |
| GAMBLING | 2.83E-69 | 5.54E-07 | 3.52E-25 | 1.52E-37 | 1.18E-09 | 2.00E-03 | 9.82E-18 | 1.46E-13 | 3.99E-36 | 1.17E-27 | 9.75E-19 | 1.51E-09 | 1.75E-25 | 5.03E-05 | 6.84E-18 | 1.19E-11 | 4.22E-29 | 1.50E-34 | 6.92E-82 | 8.79E-93 |
| SOCIAL | 3.63E-45 | 6.84E-02 | 4.73E-06 | 5.76E-05 | 2.14E-03 | 8.50E-09 | 4.39E-18 | 4.35E-19 | 3.51E-44 | 3.18E-05 | 3.06E-23 | 1.32E-10 | 1.58E-19 | 4.64E-10 | 5.19E-09 | 6.59E-10 | 1.21E-17 | 6.02E-22 | 3.80E-50 | 4.76E-71 |
| EMOTION | 3.80E-99 | 3.65E-10 | 6.43E-60 | 5.53E-87 | 1.95E-07 | 2.09E-07 | 5.87E-25 | 3.36E-68 | 1.09E-82 | 1.66E-25 | 2.46E-59 | 3.12E-29 | 2.33E-35 | 2.81E-23 | 2.65E-16 | 1.50E-22 | 1.42E-41 | 8.47E-37 | 3.41E-93 | 5.01E-132 |
| LANGUAGE | 2.06E-143 | 1.56E-19 | 5.84E-52 | 7.87E-52 | 2.13E-18 | 3.41E-52 | 4.12E-36 | 4.04E-46 | 1.81E-53 | 1.80E-91 | 1.09E-34 | 2.68E-89 | 8.42E-117 | 1.09E-14 | 1.18E-19 | 4.87E-24 | 2.37E-38 | 9.39E-48 | 2.69E-188 | 5.77E-166 |
| MOTOR | 1.02E-38 | 3.26E-05 | 2.68E-21 | 1.04E-38 | 1.78E-02 | 2.07E-02 | 9.04E-20 | 2.05E-42 | 1.36E-47 | 1.07E-12 | 2.86E-33 | 4.83E-11 | 2.12E-11 | 2.92E-06 | 5.77E-06 | 4.67E-08 | 5.08E-14 | 1.03E-04 | 1.81E-33 | 3.17E-38 |
| main effect of CONDITION | | | | | | | | | | | | | | | | | | | | |
| | SalVentAttn.1 | SalVentAttn.2 | DorsAttn.1 | DorsAttn.2 | FPN.1 | FPN.2 | FPN.3 | DMN.1 | DMN.2 | DMN.3 | Limbic.1 | Limbic.2 | AmyHip | AudLang | Thalamus | Striatum | VisPeri | VisCent | SomMot.1 | SomMot.2 |
| WM | 8.03E-01 | 3.29E-04 | 1.04E-08 | 1.48E-14 | 3.74E-19 | 1.49E-05 | 4.45E-03 | 1.07E-09 | 1.45E-11 | 1.17E-02 | 4.00E-16 | 6.63E-09 | 4.00E-09 | 5.09E-02 | 3.41E-02 | 6.27E-01 | 1.06E-12 | 6.38E-05 | 5.60E-01 | 2.48E-03 |
| RELATIONAL | 3.16E-01 | 9.14E-01 | 2.73E-01 | 5.56E-01 | 1.21E-01 | 3.88E-01 | 1.44E-01 | 4.97E-01 | 5.32E-01 | 6.79E-01 | 3.81E-01 | 7.47E-01 | 8.80E-01 | 7.54E-01 | 3.49E-01 | 2.72E-01 | 6.29E-01 | 6.22E-01 | 7.12E-01 | 9.60E-01 |
| GAMBLING | 6.09E-02 | 2.67E-01 | 6.97E-02 | 3.78E-01 | 9.06E-01 | 5.43E-01 | 4.49E-01 | 2.19E-02 | 7.72E-03 | 1.18E-01 | 7.54E-03 | 3.09E-02 | 6.21E-02 | 1.02E-01 | 8.03E-01 | 8.70E-01 | 1.81E-01 | 4.36E-01 | 2.74E-01 | 9.26E-02 |
| SOCIAL | 4.02E-01 | 2.56E-01 | 3.38E-03 | 2.18E-07 | 1.51E-01 | 1.12E-01 | 7.75E-12 | 8.46E-27 | 1.93E-10 | 2.44E-05 | 1.82E-25 | 3.96E-07 | 7.86E-23 | 4.79E-01 | 5.44E-02 | 4.27E-22 | 2.09E-12 | 8.34E-10 | 9.79E-03 | 6.10E-03 |
| EMOTION | 8.73E-01 | 3.99E-01 | 1.66E-01 | 2.79E-01 | 1.06E-01 | 9.18E-01 | 2.88E-02 | 5.86E-03 | 2.16E-01 | 3.33E-02 | 7.87E-03 | 8.99E-01 | 1.45E-01 | 5.23E-02 | 7.85E-01 | 7.21E-01 | 1.23E-01 | 4.15E-01 | 3.73E-01 | 6.99E-01 |
| LANGUAGE | 7.11E-08 | 3.57E-34 | 6.05E-02 | 1.62E-04 | 5.94E-16 | 1.31E-15 | 8.50E-28 | 3.41E-06 | 5.58E-01 | 5.55E-02 | 1.53E-10 | 9.57E-07 | 3.35E-08 | 8.15E-27 | 1.82E-01 | 1.46E-13 | 2.92E-11 | 5.22E-13 | 4.01E-01 | 2.09E-13 |
| MOTOR | 4.49E-04 | 3.17E-03 | 3.14E-04 | 4.38E-05 | 3.67E-05 | 1.00E-03 | 4.60E-04 | 4.87E-04 | 1.20E-04 | 3.07E-03 | 1.04E-04 | 6.91E-03 | 1.47E-03 | 4.16E-04 | 2.45E-03 | 5.76E-02 | 9.16E-03 | 1.09E-02 | 1.07E-04 | 3.58E-02 |
| interaction effect of SUBREGIONxCONDITION | | | | | | | | | | | | | | | | | | | | |
| | SalVentAttn.1 | SalVentAttn.2 | DorsAttn.1 | DorsAttn.2 | FPN.1 | FPN.2 | FPN.3 | DMN.1 | DMN.2 | DMN.3 | Limbic.1 | Limbic.2 | AmyHip | AudLang | Thalamus | Striatum | VisPeri | VisCent | SomMot.1 | SomMot.2 |
| WM | 8.88E-04 | 2.47E-12 | 2.07E-22 | 1.64E-09 | 3.25E-11 | 7.07E-27 | 1.40E-15 | 1.78E-27 | 3.97E-32 | 7.52E-21 | 2.34E-18 | 4.21E-17 | 4.67E-03 | 1.04E-14 | 1.58E-05 | 8.44E-03 | 8.00E-28 | 2.49E-24 | 8.72E-01 | 5.00E-03 |
| RELATIONAL | 1.47E-01 | 2.30E-01 | 3.55E-01 | 4.54E-01 | 4.21E-01 | 3.97E-02 | 5.28E-01 | 7.12E-01 | 2.64E-01 | 4.91E-01 | 1.88E-01 | 3.11E-01 | 2.24E-01 | 5.83E-03 | 2.29E-02 | 8.78E-01 | 9.38E-01 | 3.87E-02 | 5.81E-01 | 9.92E-01 |
| GAMBLING | 1.63E-01 | 5.18E-01 | 5.49E-02 | 4.40E-02 | 7.99E-01 | 3.98E-01 | 6.97E-01 | 5.47E-02 | 1.35E-01 | 9.50E-01 | 2.95E-02 | 1.60E-01 | 1.82E-01 | 4.05E-01 | 3.75E-02 | 6.93E-04 | 6.41E-04 | 2.44E-03 | 4.68E-01 | 2.28E-01 |
| SOCIAL | 7.32E-01 | 9.52E-02 | 2.34E-01 | 8.88E-02 | 6.63E-02 | 1.06E-01 | 9.43E-02 | 2.04E-08 | 5.72E-04 | 1.05E-03 | 1.57E-06 | 7.74E-01 | 6.86E-01 | 1.88E-01 | 7.55E-03 | 4.03E-07 | 7.68E-04 | 3.87E-02 | 7.51E-01 | 3.24E-01 |
| EMOTION | 3.42E-03 | 2.39E-01 | 9.19E-02 | 7.76E-02 | 2.85E-02 | 5.93E-01 | 1.87E-02 | 2.50E-09 | 1.51E-03 | 1.18E-02 | 4.58E-06 | 4.86E-02 | 7.28E-04 | 8.00E-03 | 2.23E-03 | 2.41E-02 | 5.65E-02 | 5.64E-01 | 2.97E-02 | 1.72E-01 |
| LANGUAGE | 2.86E-33 | 1.45E-21 | 2.48E-74 | 1.84E-99 | 6.24E-33 | 1.21E-65 | 6.63E-02 | 3.25E-14 | 6.96E-03 | 3.37E-11 | 2.52E-17 | 1.75E-06 | 4.42E-05 | 1.29E-48 | 1.21E-42 | 3.99E-36 | 4.35E-03 | 2.98E-04 | 2.19E-30 | 9.36E-77 |
| MOTOR | 3.15E-02 | 7.40E-01 | 1.37E-01 | 8.17E-01 | 6.04E-01 | 3.51E-01 | 5.38E-01 | 4.13E-01 | 4.65E-01 | 1.23E-01 | 3.49E-01 | 6.34E-01 | 3.30E-02 | 1.22E-02 | 2.87E-01 | 8.40E-03 | 1.78E-02 | 3.40E-01 | 2.66E-03 | 2.90E-03 |
| | | | | | | | | | | | right insular | | | | | | | | | |
| main effect of SEED | | | | | | | | | | | | | | | | | | | | |
| | SalVentAttn.1 | SalVentAttn.2 | DorsAttn.1 | DorsAttn.2 | FPN.1 | FPN.2 | FPN.3 | DMN.1 | DMN.2 | DMN.3 | Limbic.1 | Limbic.2 | AmyHip | AudLang | Thalamus | Striatum | VisPeri | VisCent | SomMot.1 | SomMot.2 |
| WM | 2.62E-207 | 2.79E-49 | 4.23E-71 | 3.42E-122 | 4.85E-57 | 1.20E-39 | 1.32E-18 | 3.67E-51 | 3.38E-05 | 3.25E-85 | 1.59E-48 | 7.27E-92 | 3.32E-115 | 2.71E-79 | 8.53E-84 | 3.24E-113 | 1.18E-60 | 3.65E-113 | 5.87E-242 | 7.08E-308 |
| RELATIONAL | 2.86E-86 | 5.18E-28 | 9.80E-55 | 1.41E-82 | 2.84E-38 | 5.86E-16 | 2.05E-05 | 2.35E-13 | 1.16E-01 | 1.74E-58 | 2.02E-11 | 1.38E-40 | 3.91E-62 | 6.42E-34 | 1.34E-56 | 6.20E-64 | 9.50E-51 | 7.56E-73 | 3.46E-128 | 3.23E-149 |
| GAMBLING | 3.84E-140 | 8.25E-44 | 2.56E-71 | 4.71E-100 | 3.67E-47 | 3.61E-28 | 5.42E-04 | 7.14E-32 | 2.99E-07 | 6.17E-69 | 1.22E-25 | 5.45E-62 | 1.42E-69 | 1.95E-53 | 3.06E-65 | 1.63E-53 | 5.70E-66 | 1.73E-88 | 2.96E-155 | 9.50E-197 |
| SOCIAL | 4.96E-143 | 4.52E-38 | 2.24E-07 | 1.46E-16 | 3.20E-08 | 8.88E-11 | 1.21E-03 | 8.59E-24 | 1.68E-07 | 1.30E-38 | 1.38E-26 | 1.50E-41 | 2.90E-62 | 1.36E-21 | 5.98E-46 | 1.06E-62 | 2.42E-70 | 1.13E-92 | 1.07E-140 | 2.76E-200 |
| EMOTION | 1.39E-181 | 1.71E-83 | 6.54E-120 | 1.21E-154 | 8.19E-44 | 5.20E-53 | 6.18E-07 | 6.60E-94 | 2.98E-46 | 2.72E-73 | 3.52E-86 | 2.64E-88 | 1.18E-100 | 3.75E-65 | 2.54E-86 | 6.21E-99 | 3.79E-77 | 4.77E-94 | 3.80E-179 | 1.39E-249 |
| LANGUAGE | 8.17E-192 | 8.45E-33 | 9.53E-172 | 2.77E-154 | 3.78E-53 | 6.73E-22 | 2.52E-02 | 1.32E-100 | 1.63E-28 | 1.93E-166 | 1.18E-67 | 1.11E-118 | 2.84E-178 | 2.63E-97 | 1.10E-107 | 3.52E-109 | 2.34E-112 | 5.80E-143 | 1.35E-284 | 0.00E+00 |
| MOTOR | 1.24E-105 | 2.71E-69 | 2.12E-80 | 2.02E-102 | 9.50E-34 | 4.48E-50 | 7.85E-09 | 8.47E-42 | 5.46E-20 | 2.63E-28 | 1.07E-35 | 2.36E-31 | 8.99E-36 | 3.07E-26 | 4.15E-47 | 5.24E-41 | 6.24E-52 | 3.06E-39 | 5.66E-89 | 4.21E-105 |
| main effect of CONDITION | | | | | | | | | | | | | | | | | | | | |
| | SalVentAttn.1 | SalVentAttn.2 | DorsAttn.1 | DorsAttn.2 | FPN.1 | FPN.2 | FPN.3 | DMN.1 | DMN.2 | DMN.3 | Limbic.1 | Limbic.2 | AmyHip | AudLang | Thalamus | Striatum | VisPeri | VisCent | SomMot.1 | SomMot.2 |
| WM | 9.66E-01 | 7.87E-01 | 5.46E-05 | 5.26E-07 | 1.62E-09 | 4.56E-02 | 6.55E-01 | 3.12E-08 | 1.75E-12 | 3.47E-01 | 1.77E-09 | 2.61E-06 | 1.94E-05 | 4.99E-04 | 6.38E-01 | 2.89E-03 | 6.83E-11 | 1.62E-03 | 6.49E-01 | 1.36E-02 |
| RELATIONAL | 3.13E-01 | 5.64E-01 | 2.54E-01 | 1.38E-01 | 5.82E-01 | 6.29E-01 | 2.88E-01 | 2.29E-01 | 3.39E-01 | 2.63E-01 | 4.18E-01 | 2.75E-01 | 5.26E-01 | 2.68E-01 | 1.70E-01 | 2.39E-01 | 2.91E-01 | 1.01E-01 | 6.74E-02 | 2.43E-01 |
| GAMBLING | 3.13E-02 | 1.98E-01 | 9.69E-01 | 3.80E-01 | 1.85E-01 | 2.89E-01 | 2.74E-01 | 8.71E-02 | 8.98E-02 | 6.08E-02 | 8.09E-02 | 1.27E-02 | 1.06E-01 | 2.08E-01 | 1.75E-01 | 3.29E-01 | 6.13E-01 | 5.79E-01 | 1.05E-02 | 4.21E-02 |
| SOCIAL | 5.38E-01 | 4.44E-01 | 1.98E-05 | 1.00E-07 | 7.27E-01 | 1.43E-02 | 2.57E-08 | 1.77E-22 | 1.93E-07 | 2.50E-03 | 8.77E-23 | 4.59E-04 | 4.21E-15 | 4.45E-01 | 6.69E-01 | 1.23E-16 | 2.60E-07 | 3.04E-05 | 1.43E-03 | 1.14E-02 |
| EMOTION | 9.37E-01 | 1.62E-01 | 4.10E-01 | 7.05E-01 | 2.52E-02 | 9.06E-01 | 1.93E-02 | 7.85E-02 | 1.64E-01 | 8.88E-02 | 4.61E-02 | 3.29E-01 | 5.50E-01 | 3.20E-01 | 6.07E-01 | 9.80E-01 | 3.35E-01 | 4.71E-01 | 2.90E-01 | 5.62E-01 |
| LANGUAGE | 4.84E-02 | 3.03E-11 | 8.32E-01 | 1.98E-01 | 3.32E-05 | 2.48E-04 | 2.59E-11 | 3.99E-01 | 2.84E-01 | 1.31E-04 | 6.14E-03 | 4.47E-09 | 4.33E-13 | 3.58E-18 | 3.44E-01 | 1.04E-01 | 1.63E-03 | 7.52E-04 | 1.27E-01 | 1.21E-11 |
| MOTOR | 9.45E-04 | 1.36E-03 | 1.50E-02 | 3.87E-04 | 4.16E-03 | 3.88E-03 | 1.66E-03 | 6.37E-02 | 4.15E-03 | 5.91E-02 | 3.68E-03 | 1.02E-01 | 3.10E-02 | 1.25E-02 | 2.25E-03 | 7.14E-04 | 4.58E-02 | 1.21E-01 | 3.18E-04 | 2.62E-02 |
| interaction effect of SUBREGIONxCONDITION | | | | | | | | | | | | | | | | | | | | |
| | SalVentAttn.1 | SalVentAttn.2 | DorsAttn.1 | DorsAttn.2 | FPN.1 | FPN.2 | FPN.3 | DMN.1 | DMN.2 | DMN.3 | Limbic.1 | Limbic.2 | AmyHip | AudLang | Thalamus | Striatum | VisPeri | VisCent | SomMot.1 | SomMot.2 |
| WM | 2.94E-04 | 4.92E-10 | 1.08E-26 | 1.14E-13 | 1.47E-09 | 4.78E-24 | 2.16E-13 | 5.25E-22 | 8.07E-27 | 1.32E-12 | 2.61E-13 | 4.11E-13 | 1.69E-01 | 1.72E-15 | 1.38E-08 | 5.52E-02 | 3.66E-26 | 2.32E-21 | 1.60E-01 | 3.99E-03 |
| RELATIONAL | 9.43E-01 | 7.40E-01 | 2.91E-01 | 1.68E-01 | 3.28E-01 | 1.63E-01 | 6.69E-01 | 6.86E-01 | 7.57E-01 | 8.87E-01 | 1.20E-01 | 4.24E-01 | 7.32E-02 | 4.14E-03 | 1.64E-01 | 3.32E-01 | 5.32E-01 | 2.73E-01 | 3.49E-01 | 9.40E-01 |
| GAMBLING | 2.69E-02 | 8.62E-01 | 2.55E-02 | 2.00E-01 | 4.11E-01 | 9.32E-01 | 9.24E-01 | 1.37E-01 | 4.54E-01 | 1.96E-01 | 2.13E-01 | 1.36E-01 | 4.43E-01 | 7.24E-01 | 9.99E-01 | 2.87E-02 | 8.04E-07 | 6.83E-05 | 9.19E-01 | 3.11E-02 |
| SOCIAL | 2.37E-01 | 1.80E-01 | 2.63E-10 | 2.01E-10 | 3.00E-03 | 2.18E-04 | 9.39E-05 | 1.65E-21 | 2.51E-03 | 3.59E-05 | 5.30E-12 | 8.90E-01 | 4.15E-02 | 7.56E-07 | 3.47E-01 | 8.37E-07 | 3.29E-03 | 4.48E-03 | 5.83E-01 | 1.73E-02 |
| EMOTION | 2.75E-02 | 3.09E-01 | 4.55E-05 | 2.92E-05 | 9.41E-01 | 3.67E-02 | 3.49E-01 | 3.96E-06 | 4.50E-03 | 6.77E-02 | 2.85E-10 | 6.91E-03 | 4.09E-03 | 3.21E-01 | 3.16E-02 | 1.62E-01 | 3.26E-03 | 5.47E-01 | 1.43E-01 | 3.26E-02 |
| LANGUAGE | 1.00E-08 | 3.65E-12 | 1.76E-32 | 5.47E-47 | 5.91E-11 | 6.94E-31 | 5.92E-07 | 4.39E-25 | 1.33E-16 | 1.55E-03 | 1.40E-38 | 3.50E-18 | 5.63E-11 | 2.79E-37 | 1.20E-27 | 1.87E-61 | 1.53E-09 | 6.65E-02 | 3.52E-13 | 1.72E-46 |
| MOTOR | 3.02E-02 | 3.65E-02 | 9.09E-01 | 3.64E-01 | 3.85E-01 | 6.98E-01 | 1.02E-01 | 1.37E-01 | 1.26E-01 | 3.40E-01 | 3.16E-02 | 4.50E-01 | 2.62E-01 | 8.23E-01 | 6.23E-02 | 6.00E-01 | 9.07E-01 | 9.89E-01 | 1.01E-01 | 1.09E-02 |

**Supplementary Table S12** Repeated ANOVA with factors of SUBREGION and CONDITION reveals significant main effects and interaction effect on task-dependent condition-specific connectivity in the HCP tasks (Faillenot atlas). Table shows p values of main effects and interaction effects for insula's connectivity to brain networks.

left insular

main effect of SUBREGION

| | SalVentAttn | SalVentAttn | DorsAttn.1 | DorsAttn.2 | FPN.1 | FPN.2 | FPN.3 | DMN.1 | DMN.2 | DMN.3 | Limbic.1 | Limbic.2 | AmyHip | AudLang | Thalamus | Striatum | VisPeri | VisCent | SomMot.1 | SomMot.2 |
|---|---|---|---|---|---|---|---|---|---|---|---|---|---|---|---|---|---|---|---|---|
| WM | 7.44E-91 | 2.70E-47 | 3.39E-59 | 5.42E-58 | 1.67E-56 | 1.29E-91 | 1.45E-87 | 3.99E-88 | 2.77E-79 | 9.27E-38 | 2.62E-72 | 1.89E-51 | 3.43E-108 | 2.18E-34 | 1.95E-41 | 9.56E-40 | 5.55E-42 | 8.01E-26 | 2.84E-191 | 1.64E-218 |
| RELATIONAL | 1.74E-22 | 2.56E-19 | 2.79E-22 | 2.48E-20 | 2.39E-16 | 3.52E-35 | 9.69E-30 | 8.37E-58 | 5.97E-43 | 8.97E-24 | 1.04E-39 | 1.22E-28 | 3.34E-30 | 3.66E-21 | 4.28E-18 | 4.37E-12 | 1.30E-16 | 8.30E-20 | 4.00E-41 | 3.02E-65 |
| GAMBLING | 1.18E-43 | 1.42E-29 | 4.06E-44 | 7.84E-44 | 2.17E-30 | 8.36E-43 | 1.15E-48 | 1.69E-73 | 1.96E-47 | 6.16E-48 | 2.01E-47 | 1.15E-35 | 3.14E-58 | 6.97E-35 | 1.35E-34 | 1.05E-32 | 5.01E-41 | 8.59E-45 | 1.93E-93 | 3.96E-91 |
| SOCIAL | 2.48E-40 | 1.16E-40 | 8.26E-20 | 3.69E-18 | 4.71E-28 | 9.83E-42 | 2.64E-54 | 1.02E-97 | 1.44E-73 | 1.33E-48 | 4.26E-68 | 6.36E-44 | 1.07E-62 | 1.30E-49 | 7.44E-39 | 8.88E-42 | 1.49E-35 | 3.09E-45 | 4.82E-71 | 2.91E-79 |
| EMOTION | 2.42E-48 | 5.61E-20 | 3.51E-59 | 3.03E-64 | 7.08E-25 | 5.06E-44 | 7.32E-51 | 7.41E-146 | 2.77E-78 | 1.69E-81 | 1.05E-100 | 1.25E-67 | 1.53E-80 | 2.25E-71 | 7.59E-43 | 4.74E-47 | 4.78E-40 | 1.70E-47 | 2.99E-87 | 1.50E-124 |
| LANGUAGE | 2.15E-109 | 4.27E-88 | 1.35E-50 | 1.02E-41 | 1.69E-50 | 3.64E-105 | 7.47E-123 | 3.59E-223 | 6.91E-142 | 9.90E-144 | 8.80E-124 | 3.41E-147 | 2.22E-197 | 2.16E-122 | 1.25E-65 | 2.61E-32 | 2.35E-89 | 5.66E-70 | 5.28E-260 | 7.80E-213 |
| MOTOR | 1.86E-14 | 3.64E-08 | 1.57E-22 | 8.19E-25 | 2.52E-10 | 1.19E-17 | 2.60E-22 | 8.97E-70 | 9.55E-46 | 6.43E-35 | 8.61E-43 | 4.42E-26 | 2.30E-36 | 6.19E-25 | 1.14E-14 | 1.79E-10 | 6.44E-17 | 3.90E-18 | 1.60E-19 | 4.96E-24 |

main effect of CONDITION

| | SalVentAttn | SalVentAttn | DorsAttn.1 | DorsAttn.2 | FPN.1 | FPN.2 | FPN.3 | DMN.1 | DMN.2 | DMN.3 | Limbic.1 | Limbic.2 | AmyHip | AudLang | Thalamus | Striatum | VisPeri | VisCent | SomMot.1 | SomMot.2 |
|---|---|---|---|---|---|---|---|---|---|---|---|---|---|---|---|---|---|---|---|---|
| WM | 6.04E-01 | 4.49E-04 | 8.39E-06 | 1.12E-09 | 5.76E-11 | 4.86E-04 | 1.15E-02 | 2.08E-05 | 8.19E-05 | 3.53E-02 | 9.78E-10 | 2.35E-05 | 1.26E-04 | 3.73E-01 | 2.30E-02 | 8.49E-01 | 1.16E-07 | 2.79E-04 | 8.96E-01 | 1.83E-01 |
| RELATIONAL | 4.74E-01 | 8.06E-01 | 2.01E-01 | 5.85E-01 | 6.61E-02 | 5.61E-01 | 9.59E-02 | 7.07E-01 | 5.40E-01 | 6.29E-01 | 3.44E-01 | 6.96E-01 | 8.30E-01 | 3.06E-01 | 1.50E-01 | 3.54E-02 | 4.37E-01 | 5.87E-01 | 7.80E-01 | 6.24E-01 |
| GAMBLING | 7.46E-02 | 2.86E-01 | 2.77E-01 | 6.69E-01 | 6.94E-01 | 5.16E-01 | 3.84E-01 | 4.49E-02 | 3.54E-02 | 1.74E-01 | 3.16E-02 | 9.47E-02 | 4.72E-02 | 2.42E-01 | 6.16E-01 | 3.29E-01 | 6.31E-01 | 7.36E-01 | 2.15E-01 | 1.47E-01 |
| SOCIAL | 3.23E-01 | 9.72E-01 | 5.72E-04 | 2.42E-09 | 3.87E-01 | 1.16E-01 | 2.89E-07 | 9.58E-19 | 1.63E-07 | 1.26E-03 | 3.49E-21 | 5.84E-06 | 6.21E-18 | 7.09E-01 | 9.86E-02 | 2.41E-17 | 1.86E-10 | 3.12E-07 | 1.63E-01 | 2.58E-01 |
| EMOTION | 4.63E-01 | 6.57E-01 | 7.06E-02 | 5.84E-02 | 5.64E-01 | 9.98E-01 | 5.99E-02 | 4.03E-02 | 1.24E-01 | 1.54E-01 | 8.55E-02 | 5.50E-01 | 1.34E-01 | 1.80E-02 | 7.16E-01 | 6.79E-01 | 1.95E-02 | 8.82E-01 | 1.75E-01 | 3.48E-01 |
| LANGUAGE | 6.70E-16 | 3.17E-47 | 1.15E-08 | 7.62E-18 | 2.84E-27 | 6.85E-33 | 1.95E-39 | 5.81E-07 | 1.18E-07 | 7.71E-01 | 1.01E-09 | 1.10E-01 | 2.76E-06 | 2.82E-04 | 2.20E-05 | 4.12E-06 | 1.63E-13 | 1.65E-14 | 5.84E-02 | 1.10E-03 |
| MOTOR | 1.81E-04 | 3.84E-04 | 6.16E-03 | 1.23E-04 | 1.48E-03 | 6.00E-03 | 9.22E-04 | 1.39E-03 | 3.39E-04 | 8.50E-03 | 2.26E-04 | 9.98E-03 | 1.41E-03 | 7.22E-04 | 6.26E-03 | 1.32E-01 | 6.65E-02 | 2.19E-02 | 1.98E-04 | 3.98E-03 |

interaction effect of SUBREGIONxCONDITION

| | SalVentAttn | SalVentAttn | DorsAttn.1 | DorsAttn.2 | FPN.1 | FPN.2 | FPN.3 | DMN.1 | DMN.2 | DMN.3 | Limbic.1 | Limbic.2 | AmyHip | AudLang | Thalamus | Striatum | VisPeri | VisCent | SomMot.1 | SomMot.2 |
|---|---|---|---|---|---|---|---|---|---|---|---|---|---|---|---|---|---|---|---|---|
| WM | 1.11E-05 | 2.22E-14 | 1.14E-37 | 1.22E-20 | 1.21E-17 | 2.69E-31 | 1.52E-09 | 3.44E-13 | 9.67E-09 | 1.09E-20 | 2.77E-04 | 9.51E-10 | 4.68E-01 | 7.85E-10 | 2.60E-08 | 9.29E-01 | 4.13E-41 | 3.55E-32 | 4.14E-01 | 2.37E-03 |
| RELATIONAL | 6.63E-01 | 3.27E-01 | 3.58E-01 | 3.15E-01 | 9.61E-01 | 9.70E-02 | 1.82E-01 | 1.48E-01 | 8.06E-03 | 2.06E-01 | 4.06E-01 | 6.31E-01 | 2.10E-01 | 7.32E-04 | 2.79E-01 | 6.78E-01 | 8.20E-01 | 2.36E-01 | 5.50E-01 | 5.03E-01 |
| GAMBLING | 1.89E-01 | 1.49E-01 | 5.09E-01 | 7.06E-01 | 7.64E-01 | 9.50E-01 | 1.34E-01 | 1.21E-04 | 1.08E-02 | 5.92E-01 | 5.29E-02 | 7.32E-01 | 3.24E-01 | 6.45E-01 | 7.82E-01 | 8.81E-01 | 1.54E-02 | 9.42E-02 | 8.70E-01 | 6.15E-01 |
| SOCIAL | 3.01E-01 | 6.94E-02 | 1.23E-02 | 8.87E-04 | 2.69E-01 | 1.76E-03 | 8.33E-01 | 2.02E-10 | 1.04E-01 | 1.01E-02 | 6.33E-06 | 4.76E-01 | 4.18E-02 | 3.80E-03 | 7.61E-01 | 3.12E-02 | 7.92E-04 | 9.27E-03 | 2.45E-01 | 1.91E-02 |
| EMOTION | 3.25E-02 | 2.71E-01 | 1.46E-03 | 1.66E-03 | 3.40E-01 | 2.50E-03 | 5.54E-01 | 9.50E-04 | 4.10E-01 | 1.02E-01 | 7.40E-02 | 5.21E-01 | 3.97E-01 | 4.36E-01 | 9.08E-02 | 9.30E-02 | 9.00E-04 | 3.89E-01 | 9.85E-02 | 9.19E-02 |
| LANGUAGE | 6.14E-07 | 4.10E-09 | 2.75E-64 | 7.60E-84 | 2.37E-19 | 6.96E-62 | 7.56E-11 | 5.94E-32 | 1.16E-18 | 2.07E-03 | 8.11E-31 | 1.26E-05 | 5.89E-05 | 2.41E-59 | 3.40E-26 | 1.45E-19 | 3.25E-04 | 1.02E-01 | 2.49E-21 | 7.10E-75 |
| MOTOR | 7.27E-01 | 5.29E-01 | 4.94E-01 | 5.56E-01 | 4.41E-01 | 1.83E-01 | 4.72E-01 | 3.93E-01 | 1.31E-01 | 5.61E-01 | 1.40E-01 | 1.95E-01 | 4.85E-01 | 6.29E-03 | 3.79E-01 | 6.44E-01 | 1.63E-01 | 1.81E-01 | 5.13E-01 | 4.53E-01 |

right insular

main effect of SUBREGION

| | SalVentAttn | SalVentAttn | DorsAttn.1 | DorsAttn.2 | FPN.1 | FPN.2 | FPN.3 | DMN.1 | DMN.2 | DMN.3 | Limbic.1 | Limbic.2 | AmyHip | AudLang | Thalamus | Striatum | VisPeri | VisCent | SomMot.1 | SomMot.2 |
|---|---|---|---|---|---|---|---|---|---|---|---|---|---|---|---|---|---|---|---|---|
| WM | 2.26E-215 | 1.66E-15 | 9.47E-75 | 9.22E-108 | 4.45E-39 | 5.57E-38 | 1.43E-36 | 1.70E-116 | 5.15E-39 | 5.47E-96 | 8.98E-87 | 1.61E-119 | 5.21E-178 | 8.30E-84 | 7.89E-66 | 3.28E-107 | 4.18E-56 | 1.15E-98 | 683641518 | 0.00E+00 |
| RELATIONAL | 6.35E-54 | 4.72E-09 | 5.51E-34 | 2.64E-46 | 4.29E-23 | 3.46E-21 | 1.79E-15 | 4.60E-38 | 1.00E-12 | 1.44E-44 | 3.33E-27 | 7.04E-41 | 4.66E-72 | 3.68E-27 | 1.04E-23 | 3.08E-19 | 9.83E-31 | 5.27E-42 | 8.22E-95 | 7.89E-120 |
| GAMBLING | 7.44E-130 | 1.44E-21 | 2.60E-67 | 4.85E-88 | 1.52E-39 | 4.09E-31 | 2.20E-20 | 8.66E-62 | 2.54E-23 | 2.47E-78 | 1.19E-38 | 3.68E-87 | 5.72E-99 | 1.12E-75 | 4.19E-68 | 2.35E-47 | 4.97E-70 | 1.16E-88 | 4.76E-179 | 1.43E-200 |
| SOCIAL | 1.42E-88 | 1.45E-14 | 2.35E-45 | 8.67E-54 | 2.01E-17 | 8.14E-26 | 2.65E-19 | 4.58E-52 | 5.30E-18 | 4.63E-49 | 1.10E-34 | 3.71E-51 | 9.70E-112 | 9.76E-41 | 2.41E-48 | 2.82E-49 | 1.31E-59 | 5.82E-55 | 1.19E-146 | 4.74E-175 |
| EMOTION | 7.38E-102 | 5.40E-23 | 3.51E-87 | 7.07E-106 | 2.06E-19 | 2.55E-39 | 1.45E-27 | 7.91E-93 | 3.15E-39 | 2.20E-63 | 7.61E-69 | 8.46E-78 | 1.32E-103 | 3.88E-69 | 4.05E-60 | 3.97E-65 | 9.62E-53 | 2.10E-52 | 5.33E-148 | 2.62E-187 |
| LANGUAGE | 1.01E-167 | 4.80E-07 | 7.44E-130 | 2.22E-91 | 1.60E-11 | 2.89E-12 | 4.70E-17 | 7.41E-142 | 1.20E-31 | 1.03E-156 | 1.28E-70 | 8.27E-137 | 4.15E-223 | 2.31E-116 | 2.76E-71 | 2.22E-45 | 2.45E-124 | 5.52E-111 | 0.00E+00 | 0.00E+00 |
| MOTOR | 7.36E-48 | 1.38E-22 | 2.96E-42 | 4.44E-58 | 8.72E-13 | 2.06E-22 | 1.43E-09 | 5.56E-46 | 4.05E-16 | 5.93E-30 | 1.71E-29 | 7.46E-20 | 4.31E-23 | 1.06E-12 | 5.33E-16 | 1.14E-14 | 1.20E-22 | 4.54E-18 | 2.67E-50 | 3.31E-60 |

main effect of CONDITION

| | SalVentAttn | SalVentAttn | DorsAttn.1 | DorsAttn.2 | FPN.1 | FPN.2 | FPN.3 | DMN.1 | DMN.2 | DMN.3 | Limbic.1 | Limbic.2 | AmyHip | AudLang | Thalamus | Striatum | VisPeri | VisCent | SomMot.1 | SomMot.2 |
|---|---|---|---|---|---|---|---|---|---|---|---|---|---|---|---|---|---|---|---|---|
| WM | 4.40E-02 | 2.07E-03 | 3.62E-09 | 2.13E-12 | 1.40E-10 | 9.26E-04 | 3.09E-01 | 6.78E-06 | 8.01E-08 | 4.12E-01 | 8.58E-10 | 5.97E-03 | 2.58E-02 | 4.02E-01 | 2.04E-02 | 1.34E-02 | 5.17E-14 | 1.26E-09 | 4.30E-02 | 9.19E-01 |
| RELATIONAL | 7.61E-01 | 4.19E-01 | 7.28E-01 | 8.16E-01 | 2.62E-01 | 6.22E-01 | 4.76E-02 | 1.27E-01 | 7.57E-02 | 9.60E-01 | 1.03E-01 | 6.54E-01 | 9.92E-01 | 9.80E-01 | 5.44E-01 | 7.39E-02 | 8.60E-01 | 3.08E-01 | 9.44E-01 | 2.72E-01 |
| GAMBLING | 5.37E-02 | 1.33E-01 | 8.37E-01 | 2.99E-01 | 5.66E-02 | 1.23E-01 | 2.75E-02 | 2.38E-03 | 5.77E-03 | 8.88E-03 | 2.86E-04 | 2.96E-02 | 1.92E-02 | 3.45E-02 | 1.02E-01 | 7.02E-02 | 4.35E-01 | 2.22E-01 | 1.02E-02 | 3.58E-02 |
| SOCIAL | 9.47E-01 | 9.56E-01 | 4.58E-06 | 6.25E-11 | 7.71E-01 | 1.98E-02 | 4.17E-07 | 1.79E-24 | 6.70E-08 | 4.17E-04 | 1.93E-23 | 1.37E-04 | 1.19E-13 | 7.43E-01 | 3.17E-01 | 6.79E-15 | 2.72E-08 | 1.99E-06 | 3.75E-02 | 1.79E-01 |
| EMOTION | 9.94E-02 | 8.34E-01 | 5.05E-02 | 2.74E-02 | 4.35E-01 | 3.68E-01 | 2.02E-01 | 1.19E-01 | 3.03E-01 | 3.10E-01 | 7.80E-02 | 8.45E-01 | 8.56E-02 | 1.58E-01 | 6.86E-01 | 6.56E-01 | 6.79E-04 | 4.20E-01 | 7.07E-02 | 1.05E-01 |
| LANGUAGE | 1.10E-13 | 2.21E-35 | 7.81E-06 | 2.30E-08 | 9.60E-21 | 2.26E-14 | 4.02E-32 | 7.88E-06 | 9.97E-03 | 9.74E-01 | 1.99E-04 | 6.46E-03 | 7.66E-08 | 3.70E-08 | 2.18E-01 | 6.13E-01 | 1.26E-08 | 9.32E-12 | 3.17E-02 | 3.77E-04 |
| MOTOR | 1.82E-02 | 1.02E-02 | 1.26E-01 | 1.60E-02 | 1.12E-01 | 3.02E-02 | 1.53E-02 | 2.56E-01 | 6.53E-03 | 4.79E-01 | 4.62E-02 | 2.25E-01 | 1.99E-01 | 2.54E-02 | 5.73E-02 | 1.06E-01 | 2.13E-01 | 5.76E-01 | 1.94E-02 | 6.35E-02 |

interaction effect of SUBREGIONxCONDITION

| | SalVentAttn | SalVentAttn | DorsAttn.1 | DorsAttn.2 | FPN.1 | FPN.2 | FPN.3 | DMN.1 | DMN.2 | DMN.3 | Limbic.1 | Limbic.2 | AmyHip | AudLang | Thalamus | Striatum | VisPeri | VisCent | SomMot.1 | SomMot.2 |
|---|---|---|---|---|---|---|---|---|---|---|---|---|---|---|---|---|---|---|---|---|
| WM | 8.05E-03 | 7.40E-05 | 1.28E-11 | 1.41E-04 | 2.89E-05 | 1.47E-12 | 6.00E-07 | 5.07E-11 | 5.40E-07 | 5.09E-12 | 2.28E-03 | 7.22E-07 | 8.58E-02 | 1.63E-05 | 3.07E-03 | 4.83E-01 | 1.01E-12 | 5.31E-11 | 5.04E-01 | 3.36E-01 |
| RELATIONAL | 4.47E-01 | 9.09E-01 | 7.87E-01 | 9.57E-01 | 7.60E-01 | 2.27E-01 | 1.65E-01 | 2.93E-01 | 1.53E-01 | 9.68E-01 | 7.70E-02 | 7.44E-01 | 2.38E-01 | 2.41E-03 | 3.69E-01 | 8.28E-01 | 2.01E-01 | 2.41E-01 | 5.84E-01 | 2.16E-01 |
| GAMBLING | 8.25E-01 | 9.71E-01 | 7.35E-01 | 8.15E-01 | 6.07E-01 | 3.05E-01 | 3.29E-01 | 7.72E-02 | 1.08E-01 | 5.25E-01 | 6.85E-02 | 6.38E-01 | 5.01E-01 | 6.72E-02 | 8.41E-01 | 5.92E-01 | 7.38E-03 | 3.50E-01 | 3.91E-01 | 3.84E-01 |
| SOCIAL | 6.74E-01 | 2.37E-01 | 2.02E-05 | 1.21E-05 | 8.06E-01 | 1.92E-04 | 1.74E-03 | 8.95E-15 | 2.08E-01 | 5.17E-06 | 1.51E-04 | 8.56E-01 | 3.15E-01 | 5.20E-01 | 1.86E-02 | 2.20E-05 | 5.19E-06 | 5.19E-04 | 5.05E-01 | 3.62E-01 |
| EMOTION | 5.01E-03 | 4.58E-03 | 2.06E-09 | 1.07E-09 | 3.29E-02 | 5.23E-03 | 4.86E-01 | 3.24E-04 | 2.08E-01 | 1.13E-01 | 1.30E-04 | 5.49E-01 | 1.55E-01 | 7.07E-01 | 2.95E-02 | 2.22E-01 | 9.43E-08 | 4.48E-04 | 8.96E-03 | 1.81E-02 |
| LANGUAGE | 7.72E-06 | 2.19E-10 | 4.64E-49 | 2.10E-55 | 4.01E-17 | 3.28E-33 | 1.05E-08 | 3.31E-27 | 6.15E-12 | 4.97E-02 | 4.97E-31 | 5.89E-03 | 1.54E-04 | 4.67E-28 | 4.92E-12 | 2.54E-17 | 2.98E-04 | 1.26E-02 | 7.31E-15 | 6.12E-47 |
| MOTOR | 9.28E-01 | 6.31E-01 | 5.85E-01 | 7.07E-01 | 2.73E-01 | 6.38E-01 | 6.39E-01 | 2.78E-01 | 2.89E-01 | 2.53E-01 | 5.28E-01 | 5.16E-01 | 6.11E-01 | 2.01E-01 | 8.78E-01 | 2.83E-02 | 2.76E-01 | 1.61E-01 | 4.95E-01 | 2.17E-01 |

**Supplementary Table S13** Repeated ANOVA with factors of SUBREGION and CONDITION reveals significant main effects and interaction effect on task-dependent condition-specific connectivity in the HCP tasks (Julich atlas). Table shows p values of main effects and interaction effects for insula's connectivity to brain networks.

left insular

main effect of SUBREGION

| | SalVentAttn | SalVentAttn | DorsAttn.1 | DorsAttn.2 | FPN.1 | FPN.2 | FPN.3 | DMN.1 | DMN.2 | DMN.3 | Limbic.1 | Limbic.2 | AmyHip | AudLang | Thalamus | Striatum | VisPeri | VisCent | SomMot.1 | SomMot.2 |
|---|---|---|---|---|---|---|---|---|---|---|---|---|---|---|---|---|---|---|---|---|
| WM | 0.00E+00 | 0.00E+00 | 1.49E-264 | 0.00E+00 | 0.00E+00 | 0.00E+00 | 0.00E+00 | 1.64E-149 | 1.08E-219 | 8.35E-109 | 1.08E-78 | 1.61E-107 | 7.67E-179 | 1.77E-197 | 5.36E-231 | 1.19E-245 | 3.89E-227 | 2.34E-215 | 0.00E+00 | 0.00E+00 |
| RELATIONAL | 3.81E-217 | 4.59E-287 | 3.25E-128 | 1.84E-175 | 5.84E-154 | 4.79E-224 | 1.39E-197 | 5.19E-91 | 1.87E-175 | 1.29E-64 | 9.58E-44 | 3.91E-37 | 8.66E-58 | 1.54E-121 | 5.16E-159 | 1.72E-123 | 2.10E-96 | 5.64E-144 | 9.13E-157 | 1.35E-227 |
| GAMBLING | 5.98E-265 | 0.00E+00 | 1.66E-157 | 1.08E-224 | 3.78E-155 | 1.54E-217 | 5.63E-211 | 9.27E-75 | 6.25E-137 | 3.85E-80 | 4.07E-54 | 1.84E-55 | 5.42E-80 | 3.06E-126 | 1.94E-165 | 1.04E-166 | 5.14E-126 | 8.61E-136 | 4.39E-178 | 1.65E-229 |
| SOCIAL | 5.61E-263 | 3.59E-281 | 7.78E-72 | 1.71E-77 | 6.87E-122 | 6.68E-166 | 5.41E-254 | 3.59E-114 | 1.39E-189 | 5.47E-57 | 6.15E-77 | 1.12E-50 | 4.52E-93 | 3.68E-125 | 8.87E-126 | 1.22E-222 | 4.51E-174 | 1.41E-238 | 2.40E-181 | 7.71E-228 |
| EMOTION | 2.58E-278 | 2.31E-294 | 1.55E-146 | 2.21E-244 | 2.37E-81 | 3.75E-143 | 6.10E-120 | 8.53E-147 | 2.74E-128 | 6.57E-98 | 2.35E-122 | 3.96E-74 | 2.16E-128 | 1.02E-92 | 2.63E-105 | 1.33E-134 | 6.02E-111 | 6.88E-116 | 4.94E-208 | 5.84E-314 |
| LANGUAGE | 0.00E+00 | 0.00E+00 | 1.45E-258 | 1597930119 | 0.00E+00 | 0.00E+00 | 0.00E+00 | 1.05E-260 | 6.26E-161 | 7.01E-272 | 3.33E-125 | 2.60E-230 | 0.00E+00 | 9.06E-263 | 5.99E-207 | 3.32E-283 | 2.93E-187 | 2.14E-228 | 0.00E+00 | 0.00E+00 |
| MOTOR | 1.29E-119 | 2.08E-106 | 2.47E-67 | 1.85E-114 | 1.90E-39 | 3.06E-57 | 8.64E-39 | 2.32E-54 | 5.20E-41 | 1.65E-37 | 1.16E-40 | 9.71E-16 | 1.34E-35 | 2.76E-32 | 4.88E-57 | 1.75E-57 | 1.05E-55 | 4.75E-48 | 1.13E-93 | 3.30E-99 |

main effect of CONDITION

| | SalVentAttn | SalVentAttn | DorsAttn.1 | DorsAttn.2 | FPN.1 | FPN.2 | FPN.3 | DMN.1 | DMN.2 | DMN.3 | Limbic.1 | Limbic.2 | AmyHip | AudLang | Thalamus | Striatum | VisPeri | VisCent | SomMot.1 | SomMot.2 |
|---|---|---|---|---|---|---|---|---|---|---|---|---|---|---|---|---|---|---|---|---|
| WM | 3.55E-01 | 4.29E-01 | 6.04E-02 | 1.14E-05 | 1.25E-08 | 6.25E-01 | 9.61E-01 | 9.13E-14 | 3.24E-14 | 1.95E-05 | 1.04E-14 | 4.59E-11 | 5.80E-08 | 2.55E-04 | 5.68E-01 | 5.50E-01 | 4.66E-03 | 2.04E-01 | 5.65E-01 | 2.12E-01 |
| RELATIONAL | 6.37E-01 | 8.05E-01 | 3.41E-01 | 8.77E-01 | 2.34E-01 | 9.68E-01 | 3.58E-01 | 8.73E-01 | 8.18E-01 | 6.31E-01 | 2.81E-01 | 9.39E-01 | 5.32E-01 | 2.25E-01 | 1.55E-01 | 4.29E-01 | 4.66E-01 | 7.84E-01 | 7.88E-01 | 6.88E-01 |
| GAMBLING | 2.73E-01 | 3.20E-01 | 1.26E-01 | 5.11E-01 | 9.26E-01 | 3.16E-01 | 2.94E-01 | 1.43E-02 | 1.18E-02 | 1.15E-01 | 1.29E-02 | 1.18E-01 | 1.17E-01 | 7.65E-02 | 5.65E-01 | 9.17E-01 | 9.88E-02 | 1.89E-01 | 1.68E-01 | 3.31E-01 |
| SOCIAL | 6.16E-01 | 3.41E-01 | 2.38E-01 | 5.12E-04 | 1.74E-01 | 6.68E-01 | 1.00E-08 | 3.55E-20 | 5.29E-10 | 1.92E-03 | 1.00E-21 | 7.73E-06 | 1.59E-14 | 9.84E-01 | 4.24E-02 | 5.11E-19 | 2.88E-13 | 8.03E-05 | 2.42E-02 | 4.30E-01 |
| EMOTION | 3.39E-01 | 9.36E-01 | 6.33E-02 | 4.97E-02 | 8.09E-01 | 8.29E-01 | 1.27E-01 | 2.11E-02 | 2.66E-01 | 2.06E-01 | 1.49E-02 | 6.95E-01 | 1.79E-01 | 2.83E-01 | 6.99E-01 | 7.42E-01 | 1.16E-01 | 8.16E-01 | 2.65E-01 | 1.71E-01 |
| LANGUAGE | 5.64E-09 | 4.04E-22 | 2.74E-07 | 2.05E-12 | 3.61E-14 | 4.95E-17 | 2.09E-13 | 2.57E-01 | 3.30E-01 | 6.55E-01 | 1.25E-02 | 1.53E-02 | 3.40E-05 | 2.12E-03 | 2.23E-02 | 7.86E-04 | 6.32E-07 | 1.14E-08 | 4.31E-02 | 5.32E-01 |
| MOTOR | 1.37E-05 | 2.26E-05 | 2.85E-04 | 1.17E-06 | 6.87E-05 | 4.79E-05 | 3.26E-05 | 3.49E-04 | 2.23E-05 | 1.91E-03 | 1.31E-04 | 1.85E-03 | 2.86E-03 | 2.66E-04 | 6.33E-05 | 4.58E-03 | 4.44E-03 | 1.25E-02 | 9.92E-06 | 6.52E-03 |

interaction effect of SUBREGIONxCONDITION

| | SalVentAttn | SalVentAttn | DorsAttn.1 | DorsAttn.2 | FPN.1 | FPN.2 | FPN.3 | DMN.1 | DMN.2 | DMN.3 | Limbic.1 | Limbic.2 | AmyHip | AudLang | Thalamus | Striatum | VisPeri | VisCent | SomMot.1 | SomMot.2 |
|---|---|---|---|---|---|---|---|---|---|---|---|---|---|---|---|---|---|---|---|---|
| WM | 8.97E-07 | 2.04E-32 | 2.62E-98 | 4.32E-49 | 1.53E-47 | 8.16E-79 | 1.14E-20 | 4.32E-21 | 1.74E-22 | 1.30E-39 | 3.90E-20 | 1.39E-20 | 8.10E-11 | 5.24E-24 | 5.30E-14 | 7.55E-02 | 2.37E-111 | 1.24E-72 | 1.50E-01 | 2.50E-06 |
| RELATIONAL | 7.84E-01 | 5.68E-01 | 3.05E-01 | 1.35E-01 | 1.30E-01 | 5.36E-02 | 3.32E-01 | 8.21E-01 | 5.56E-01 | 3.79E-01 | 4.81E-01 | 6.87E-01 | 5.42E-01 | 2.87E-04 | 2.28E-01 | 7.07E-01 | 1.90E-01 | 2.48E-02 | 9.25E-01 | 7.06E-01 |
| GAMBLING | 3.02E-01 | 8.58E-01 | 3.95E-01 | 6.88E-01 | 2.99E-01 | 5.48E-01 | 8.84E-01 | 9.84E-01 | 7.56E-01 | 7.58E-01 | 2.85E-01 | 4.51E-01 | 5.34E-01 | 8.01E-01 | 9.88E-01 | 2.20E-01 | 6.60E-03 | 4.40E-03 | 6.54E-01 | 6.15E-01 |
| SOCIAL | 7.77E-02 | 5.10E-03 | 1.58E-19 | 1.88E-26 | 1.89E-04 | 2.22E-09 | 1.69E-01 | 3.47E-16 | 4.48E-03 | 2.17E-02 | 6.99E-10 | 7.45E-04 | 2.62E-14 | 5.15E-21 | 7.00E-01 | 5.06E-02 | 4.31E-04 | 2.28E-06 | 4.69E-02 | 1.35E-02 |
| EMOTION | 5.15E-02 | 2.36E-01 | 1.37E-02 | 5.56E-02 | 4.32E-02 | 6.58E-02 | 3.07E-03 | 2.21E-11 | 6.67E-04 | 1.50E-04 | 3.81E-06 | 8.07E-03 | 5.87E-03 | 4.47E-04 | 1.38E-02 | 9.30E-03 | 1.31E-04 | 4.23E-01 | 5.71E-02 | 2.14E-02 |
| LANGUAGE | 4.18E-39 | 5.44E-49 | 1.00E-143 | 2.86E-192 | 1.31E-58 | 7.22E-124 | 2.67E-59 | 1.56E-118 | 8.21E-126 | 7.15E-15 | 1.88E-130 | 1.79E-56 | 1.46E-34 | 1.33E-214 | 3.82E-58 | 1.83E-105 | 2.04E-29 | 1.55E-23 | 3.52E-46 | 1.28E-200 |
| MOTOR | 7.10E-02 | 4.12E-01 | 2.23E-02 | 1.99E-01 | 1.28E-04 | 3.50E-01 | 2.47E-01 | 2.06E-02 | 3.36E-02 | 4.26E-04 | 6.23E-03 | 4.20E-01 | 7.89E-03 | 4.91E-03 | 1.38E-01 | 5.66E-03 | 9.32E-03 | 1.41E-05 | 1.87E-02 | 2.36E-01 |

right insular

main effect of SUBREGION

| | SalVentAttn | SalVentAttn | DorsAttn.1 | DorsAttn.2 | FPN.1 | FPN.2 | FPN.3 | DMN.1 | DMN.2 | DMN.3 | Limbic.1 | Limbic.2 | AmyHip | AudLang | Thalamus | Striatum | VisPeri | VisCent | SomMot.1 | SomMot.2 |
|---|---|---|---|---|---|---|---|---|---|---|---|---|---|---|---|---|---|---|---|---|
| WM | 0.00E+00 | 0.00E+00 | 2.82E-253 | 0.00E+00 | 3146153672 | 0.00E+00 | 0.00E+00 | 2.59E-240 | 1.83E-173 | 1.63E-203 | 1.30E-156 | 6.72E-251 | 0.00E+00 | 1.79E-238 | 3.40E-264 | 0.00E+00 | 3.66E-207 | 9.63E-278 | 0.00E+00 | 0.00E+00 |
| RELATIONAL | 9.36E-168 | 4.21E-201 | 5.24E-127 | 3.23E-182 | 4.32E-126 | 1.08E-142 | 3.50E-147 | 7.52E-85 | 3.80E-84 | 3.24E-94 | 1.62E-45 | 9.81E-69 | 8.23E-118 | 2.84E-107 | 1.17E-138 | 4.92E-168 | 9.16E-99 | 4.06E-154 | 1.05E-222 | 1.61E-290 |
| GAMBLING | 3282292062 | 2500022741 | 3.67E-192 | 3.45E-268 | 3.27E-184 | 2.67E-233 | 1.45E-220 | 1.40E-124 | 2.06E-105 | 3.06E-146 | 1.44E-76 | 1.59E-154 | 1.59E-164 | 2.68E-183 | 9.19E-182 | 3.88E-214 | 2.12E-149 | 9.13E-196 | 0.00E+00 | 0.00E+00 |
| SOCIAL | 0.00E+00 | 1.20E-269 | 4.95E-123 | 2.01E-165 | 3.43E-129 | 6.40E-158 | 1.85E-196 | 6.36E-132 | 1.85E-112 | 2.65E-131 | 1.01E-99 | 3.56E-139 | 6.47E-248 | 1.24E-161 | 3.40E-195 | 4.65E-298 | 3.96E-187 | 5.00E-245 | 0.00E+00 | 0.00E+00 |
| EMOTION | 1.73E-259 | 1.56E-262 | 1.16E-215 | 2.36E-298 | 8.07E-92 | 1.47E-116 | 7.09E-102 | 5.71E-162 | 1.07E-59 | 8.01E-143 | 5.18E-158 | 1.52E-134 | 1.11E-232 | 1.57E-130 | 4.68E-154 | 1.02E-215 | 9.23E-160 | 1.14E-168 | 0.00E+00 | 0.00E+00 |
| LANGUAGE | 0.00E+00 | 0.00E+00 | 0.00E+00 | 0.00E+00 | 0.00E+00 | 0.00E+00 | 0.00E+00 | 0.00E+00 | 1.05E-104 | 0.00E+00 | 2.58E-179 | 2.48E-299 | 0.00E+00 | 1.96E-252 | 3.09E-260 | 0.00E+00 | 2.86E-275 | 0.00E+00 | 0.00E+00 | 0.00E+00 |
| MOTOR | 2.54E-141 | 6.62E-110 | 1.05E-89 | 6.48E-135 | 8.08E-46 | 9.25E-67 | 3.18E-35 | 7.97E-55 | 2.23E-21 | 9.17E-56 | 2.95E-34 | 1.18E-56 | 3.28E-73 | 5.20E-46 | 3.77E-65 | 4.10E-83 | 1.41E-59 | 4.24E-63 | 2.63E-125 | 4.96E-155 |

main effect of CONDITION

| | SalVentAttn | SalVentAttn | DorsAttn.1 | DorsAttn.2 | FPN.1 | FPN.2 | FPN.3 | DMN.1 | DMN.2 | DMN.3 | Limbic.1 | Limbic.2 | AmyHip | AudLang | Thalamus | Striatum | VisPeri | VisCent | SomMot.1 | SomMot.2 |
|---|---|---|---|---|---|---|---|---|---|---|---|---|---|---|---|---|---|---|---|---|
| WM | 7.07E-01 | 9.74E-01 | 3.36E-03 | 9.83E-06 | 2.65E-06 | 9.15E-01 | 4.01E-02 | 3.91E-12 | 7.81E-16 | 5.14E-02 | 5.83E-13 | 3.33E-07 | 5.10E-04 | 8.13E-04 | 7.90E-01 | 1.01E-01 | 8.66E-07 | 6.13E-04 | 5.60E-01 | 5.93E-01 |
| RELATIONAL | 8.51E-01 | 7.51E-01 | 9.36E-01 | 7.66E-01 | 5.26E-01 | 9.67E-01 | 1.82E-01 | 4.58E-01 | 3.72E-01 | 6.48E-01 | 5.11E-01 | 6.17E-01 | 8.59E-01 | 4.33E-01 | 2.77E-01 | 1.45E-01 | 9.08E-01 | 2.71E-01 | 5.28E-01 | 5.02E-01 |
| GAMBLING | 7.25E-02 | 1.72E-01 | 8.62E-01 | 4.25E-01 | 2.05E-01 | 1.64E-01 | 4.65E-02 | 7.47E-04 | 7.76E-03 | 1.42E-02 | 4.33E-04 | 1.63E-02 | 1.05E-02 | 6.30E-02 | 8.81E-02 | 1.64E-01 | 8.61E-01 | 5.34E-01 | 9.70E-04 | 3.11E-02 |
| SOCIAL | 9.24E-01 | 3.82E-01 | 7.32E-04 | 7.91E-07 | 6.85E-01 | 3.66E-01 | 5.15E-10 | 2.83E-25 | 1.21E-09 | 1.42E-04 | 1.68E-23 | 7.53E-05 | 2.21E-11 | 8.83E-01 | 1.64E-01 | 7.99E-16 | 7.55E-10 | 6.92E-05 | 6.95E-02 | 4.33E-01 |
| EMOTION | 1.51E-01 | 8.45E-01 | 7.32E-03 | 6.08E-03 | 7.18E-01 | 3.16E-01 | 1.50E-01 | 1.90E-02 | 1.40E-01 | 2.12E-01 | 8.56E-03 | 3.97E-01 | 1.90E-01 | 1.95E-01 | 3.94E-01 | 2.96E-01 | 4.37E-03 | 3.18E-01 | 3.61E-02 | 6.26E-02 |
| LANGUAGE | 1.61E-05 | 1.79E-12 | 1.81E-03 | 7.50E-04 | 1.10E-08 | 1.82E-04 | 1.20E-09 | 2.26E-01 | 5.32E-01 | 3.47E-01 | 3.20E-01 | 1.03E-04 | 4.73E-08 | 2.78E-06 | 5.06E-01 | 2.90E-03 | 1.98E-03 | 1.16E-05 | 1.34E-01 | 2.13E-02 |
| MOTOR | 1.16E-02 | 8.56E-03 | 6.28E-02 | 3.99E-03 | 1.47E-02 | 1.07E-02 | 2.00E-03 | 4.32E-02 | 4.00E-03 | 9.02E-02 | 3.72E-03 | 8.47E-02 | 6.20E-02 | 3.53E-02 | 5.59E-03 | 3.94E-02 | 1.05E-01 | 3.79E-01 | 1.20E-02 | 2.00E-01 |

interaction effect of SUBREGIONxCONDITION

| | SalVentAttn | SalVentAttn | DorsAttn.1 | DorsAttn.2 | FPN.1 | FPN.2 | FPN.3 | DMN.1 | DMN.2 | DMN.3 | Limbic.1 | Limbic.2 | AmyHip | AudLang | Thalamus | Striatum | VisPeri | VisCent | SomMot.1 | SomMot.2 |
|---|---|---|---|---|---|---|---|---|---|---|---|---|---|---|---|---|---|---|---|---|
| WM | 8.81E-14 | 6.72E-29 | 5.63E-89 | 3.95E-42 | 1.22E-35 | 1.44E-70 | 7.45E-20 | 2.39E-23 | 9.72E-19 | 2.03E-35 | 1.15E-14 | 1.31E-19 | 5.51E-03 | 1.44E-28 | 2.15E-18 | 4.23E-09 | 3.83E-95 | 3.26E-68 | 4.08E-02 | 3.11E-02 |
| RELATIONAL | 7.87E-01 | 6.37E-01 | 3.24E-01 | 5.24E-01 | 1.83E-01 | 2.77E-01 | 6.38E-02 | 8.93E-02 | 4.67E-01 | 7.80E-01 | 2.33E-02 | 9.88E-01 | 5.89E-01 | 4.75E-02 | 3.55E-01 | 8.74E-01 | 7.20E-01 | 5.73E-01 | 9.06E-01 | 8.24E-01 |
| GAMBLING | 2.16E-01 | 2.19E-01 | 4.55E-01 | 4.98E-01 | 5.95E-01 | 4.38E-01 | 7.41E-01 | 5.03E-01 | 4.23E-01 | 3.00E-01 | 3.13E-01 | 1.54E-01 | 2.09E-01 | 3.94E-01 | 9.53E-01 | 2.92E-01 | 2.46E-07 | 3.11E-03 | 3.90E-02 | 5.36E-01 |
| SOCIAL | 1.59E-02 | 5.30E-04 | 8.87E-19 | 9.34E-20 | 2.33E-04 | 1.45E-12 | 1.92E-02 | 1.69E-17 | 4.45E-02 | 2.06E-05 | 4.36E-07 | 5.07E-03 | 3.02E-07 | 9.96E-11 | 6.04E-03 | 7.93E-04 | 7.63E-10 | 2.80E-11 | 2.82E-01 | 2.57E-04 |
| EMOTION | 9.08E-02 | 1.41E-01 | 1.80E-07 | 1.39E-03 | 7.07E-02 | 5.63E-03 | 1.05E-01 | 1.78E-04 | 3.77E-02 | 2.07E-01 | 4.24E-03 | 9.05E-02 | 1.49E-04 | 9.51E-02 | 1.11E-01 | 1.70E-02 | 8.07E-10 | 3.78E-01 | 1.09E-01 | 6.27E-02 |
| LANGUAGE | 6.48E-30 | 1.26E-74 | 2.55E-116 | 3.22E-145 | 5.55E-72 | 2.18E-92 | 4.00E-87 | 3.20E-120 | 2.80E-93 | 2.41E-08 | 1.32E-139 | 1.21E-32 | 8.83E-23 | 3.55E-141 | 2.44E-34 | 8.03E-118 | 2.91E-34 | 3.95E-33 | 7.03E-29 | 1.21E-118 |
| MOTOR | 1.30E-01 | 2.92E-01 | 2.57E-01 | 1.76E-02 | 8.69E-02 | 6.20E-01 | 8.72E-01 | 9.67E-01 | 6.96E-01 | 9.36E-01 | 9.93E-01 | 2.79E-01 | 8.50E-01 | 7.52E-01 | 1.48E-01 | 4.00E-02 | 7.56E-01 | 4.85E-01 | 2.41E-01 | 8.21E-01 |
